# FGF signalling is involved in cumulus migration in the common house spider *Parasteatoda tepidariorum*

**DOI:** 10.1101/2021.10.01.462731

**Authors:** Ruixun Wang, Linda Karadas, Philipp Schiffer, Matthias Pechmann

## Abstract

Cell migration is a fundamental component during the development of most multicellular organisms. In spiders, the collective migration of a signalling centre, known as the cumulus, is required to set the dorsoventral body axis of the embryo.

Here, we show that FGF signalling plays an important role during cumulus migration in the spider *Parasteatoda tepidariorum*. Spider embryos with reduced FGF signalling lack cumulus migration and display dorsoventral patterning defects. Our study reveals that cumulus expression of several FGF signalling components is regulated by the transcription factor *Ets4*. In conjunction with a previous study, we show that the expression of *fgf8* in the germ-disc is regulated via the Hedgehog signalling pathway. We also demonstrate that FGF signalling influences the BMP signalling pathway activity in the region around cumulus cells.

Finally, we show that FGF signalling might also influence cumulus migration in basally branching spiders and we propose a hypothetical model in which *fgf8* acts a chemo-attractant to guide cumulus cells towards the future dorsal pole of the spider embryo.

## Introduction

In vertebrates like mice and humans, almost two dozen of ligands and four receptors are responsible for the regulation of one of the most complex signalling pathways, the Fibroblast-Growth-Factor-Receptor (FGFR) signalling pathway. Differential splicing of the receptors and different ligand/receptor combinations are able to regulate a wealth of functions including axis establishment, cell proliferation, differentiation and survival (reviewed in [1,2]). In contrast, the genomes of invertebrates such as beetles or flies only have a handful of fibroblast growth factor receptors and ligands ([3–5] and reviewed in [6]). Nevertheless, also in these animals FGFR signalling is involved in a variety of important developmental processes, which include the formation of the mesoderm or the branching of the tracheal system (summarized in [4,6]). In *Drosophila*, these two processes involve the regulation of directed cell migration, another widely conserved feature of the FGFR-signalling pathway. By acting as chemo-attractants, FGFs are able to attract cells by altering components of the cytoskeleton, which in turn leads to the formation of filo- and lammellopodia (e.g. [7,8], reviewed in [6]).

In the nematode *Caenorhabditis elegans*, EGL-15 and EGL-17 (the FGF receptor and FGF orthologs of *C. elegans*) are involved in a chemo-attractive mechanism to guide sex myoblasts to their final destination [9–11].

Also in vertebrates FGF signalling controls cell migration in various tissues. In mouse embryos, endoderm- and mesoderm-derived tissues depend on the outward migration of cells from the primitive streak, and this process is controlled by a small subset of FGFs [12].

Setting up the dorsoventral (DV) body axis of chelicerate species, like spiders, involves the migration of the cumulus (a cluster of cells that possesses organizing capacities). Cumulus cells are inducing the DV body axis via the secretion of Dpp (the BMP2/4 ortholog) and the subsequent activation of the BMP signalling pathway in a subset of germ-disc cells [13–19]. As the name implies, the germ-disc is radially symmetric and gives rise to the embryo proper. Cumulus transplantation or the local activation of the BMP signalling pathway demonstrated that all germ-disc cells have the capacity to develop either into ectodermal embryonic or into dorsal field cells [18,20,21]. For this reason, the decision if a germ-disc cell develops into ectoderm or changes its fate towards a dorsal field cell depends on the signalling and the direction of cumulus migration.

It is still unclear if the direction of cumulus migration in the radially symmetric field of germ-disc cells is actively guided via unknown factors, or if cumulus migration is a completely stochastic process. Hedgehog signalling was found to play a role in cumulus migration [22,23]. However, it is unknown if Hedgehog is directly or indirectly involved during this important developmental process.

Here we show that cumulus migration in the common-house spider *Parasteatoda tepidariorum* is highly dependent on FGF signalling. We show that a FGF ligand is asymmetrically expressed in the early germ-disc of many spider embryos and we propose a hypothetical model in which FGF signalling is involved in the guidance of the cumulus cells towards the periphery of the germ-disc.

## Methods

### Spider husbandry and embryology

For all experiments we used our Cologne laboratory culture of *Parasteatoda tepidariorum*. Adults and juveniles of *Parasteatoda tepidariorum* were kept in plastic vials at room temperature and were fed with *Drosophila melanogaster* and crickets (*Gryllus bimaculatus*). Embryos were collected and fixed as described previously [19] and staged according to [24]. *Acanthoscurria geniculata* embryos were kept as described previously [18]. Adults and juveniles of *Steatoda grossa* were collected near Cologne (Hürth, Germany) and were kept under the same conditions as *Parasteatoda tepidariorum*.

### Gene cloning

CLC Main Workbench 7 (QIAGEN Aarhus A/S) was used to perform a local tBLASTn [25] against the *Parasteatoda tepidariorum* official AUGUSTUS gene set (https://i5k.nal.usda.gov/Parasteatoda_tepidariorum, [26]). For this, protein sequences of known homology were downloaded from FlyBase [27] and NCBI. Putatively identified genes were reciprocally BLASTed against the online NCBI nr database using BLASTx to confirm their identity.

PCR amplification and cloning of genes was performed using standard techniques. Genes were cloned into the pCR4 and the pCRII vector (ThermoFisher scientific). *Pt-Ets4, Pt-fork-head, Pt-orthodenticle, Pt-short-gastrulation, Pt-decapentaplegic, Pt-hedgehog* and *Pt-caudal* were isolated previously [19,28,29].

A *Pt-fgf8* DNA fragment of approximately 2kb, including the full coding sequence and a portion of the 5’UTR and the 3’UTR, was amplified using Pt-fgf8-Fw (5’-CATCTCTTCGCTCTCCGCGC-3’) and Pt-fgf8-Rev (5’-GAATGCTCGTGCAAAGAGAGTG-3’). *Pt-dof, Pt-fgf1, Pt-FGFR1* and *Pt-FGFR2* were amplified using Pt-dof–Fw (5’-GAAATGGCTCCTGTCGACGTTAC-3’), Pt-dof–Rev (5’-CAATACTGGAACAGGTTGAGCTG-3’), Pt-fgf1-Fw (5’-GTGGATAGAGGCATACCGAGT-3’), Pt-fgf1-Rev (5’-CGGAACACCTCTACGGAACG-3’), Pt-FGFR1-Fw (5’-GACATATGCTGAGGAAGATAATAG-3’), Pt-FGFR1-Rev (5’-CAAACGTTATTTGAATCTGAATC-3’), Pt-FGFR2-Fw (5’-GTCACAGTCATTTTAGGCTTG-3’) and Pt-FGFR2-Rev (5’-CAGTGTACATCTCCTGAGGTAC-3’) primer, respectively.

The published transcriptome [18] was searched for *Acanthoscurria geniculata fgf1, fgf8* and *fgf17* sequences. Total RNA was isolated and cDNA was produced as described in [18]. *Ag-fgf* gene sequences were isolated using the following primer combinations: Ag-fgf1-Fw (5’-GTG CAA CGA GGA CAA CTA TTC AG-3’) and Ag-fgf1-Rev (5’-GAG TAC TAT ACA TTT CAA AAC ACC G-3’); Ag-fgf8-Fw (5’-CTC CAT GCT TCA CCG TTG ATG-3’) and Ag-fgf8-Rev (5’-CTG CTG TCA TTG GCA CTT GTC-3’); Ag-fgf17-Fw (5’-GAA CAG CCT GAG CAA ATG CCA C-3’) and Ag-fgf17-Rev (5’-CAC CTG AGA ATC TTG CTG GAC AC-3’).

Nucleotide sequences of *fgf8* and *fgf17* of different spider species were blasted against the combined RNA-Seq from 19 Libraries sequenced from False Black Widow (*Steatoda grossa*) (SRR1539523). Blast hits were manually aligned and primer sequences of *Steatoda grossa fgf8* and *fgf17* were designed according to this alignment. Total RNA of *Steatoda grossa* was extracted using TRIzol Reagent (life technologies) and cDNA was produced using the RNA to cDNA EcoDry Premix (TaKaRa Clontec, double primed). *Sg-fgf8* and *Sg-fgf17* were amplified using Sg-fgf8-Fw (5’-CAG GTT ACT ACA GAT TGC CTC C-3’) and Sg-fgf8-Rev (5’-GTA CGT GTG CTC CCT CTT GAT G-3’) and Sg-fgf17-Fw (5’-GTT CAT CTG CAC TTC TGT CAT GG-3’) and Sg-fgf17-Rev (5’-CCG TGT GTC ATT CCA GAG ATG-3’) primer, respective.

### Transcriptome sequencing of cumulus cells

To knock down *Pt-Ets4*, several spider females were injected with *Pt-Ets4* dsRNA (nucleotide 11-697 of aug3.g4238.t1 was used for the knockdown) as described in [19]. While the majority of stage 4 control (dsRed dsRNA was used for the control injections) and *Pt-Ets4* RNAi embryos of 3^rd^ and 4^th^ cocoons were formaldehyde fixed and stored in 100% MeOH at −20**°**C, around 50 embryos of each cocoon were monitored under oil. Subsequently, the primary thickening of fixed control embryos and of fixed embryos where the siblings of the same cocoon showed the characteristic *Pt-Ets4* knockdown phenotype (see [19]) was extracted using sharp needles. The primary thickening of 100-120 embryos was extracted for each replicate. For each condition (control vs. Ets4 RNAi) the embryos of three independently produced cocoons were used. Overall, three biological and technical replicates were produced per condition. The total RNA of the extracted cells was isolated using the Quick-RNA™ FFPE Kit (Zymo Research). Library preparation and sequencing was carried out at the Cologne Center for Genomics, using a paired-end 2×150nt approach on the Illimuna NovaSeq platform. The sequenced reads were subjected to adaptor and quality threshold trimming using fastp [30]. Cleaned reads where mapped against the published *P. tepidariorum* gene set [26] using kallisto [31] and the expression patterns analysed in Degust (David Powell (2015), drpowell/degust v3.2.0 (v3.2.0); Zenodo; https://doi.org/10.5281/zenodo.3258933).

### Phylogenetic analysis

The FGF core sequence (see [3]) was used to align FGF proteins of different species. In case of Dof related proteins, only the conserved DBB (Dof, BCAP, BANK region) motif as well as the region of the ankyrin repeats were used to generate the phylogeny. Alignments were produced using MUSCLE [32] and aligned sequences were trimmed using TrimAl with the GappyOut setting [33]. Maximum likelihood based phylogenies were constructed using PhyML at ‘phylogeny.fr’ [34]. Final phylogenies were generated with the WAG substitution model and 1000 bootstrap replicates [35]. Gene bank accession numbers or sequences can be found in the in supplemental information (figure legends of Fig. S1 and S5 and Supplemental Sequences).

### Parental RNAi experiments

Adult spider females were injected 3-4 times (in a 2-3 day interval) with 2µl [2-3µg/µl] dsRNA solution. *Pt-ptc* and *Pt-Ets4* pRNAi experiments were performed as described in [19]. Double stranded RNA was produced using the T7 MEGAscript Kit (ThermoFisher scientific).

During the initial RNAi screen *Pt-fgf8* (AUGUSTUS prediction: aug3.g5611.t1; Schwager et al. 2017) was amplified using the g5611-Fw (5’-CAGGTTACTACAGATTGCCTCC-3’) and g5611-Rev (5’-GCACTTTCGTTCGTATTCATAG-3’) primer. Template to generate dsRNA was amplified using the T7 overhang primer T7-g5611-Fw (5’-GTAATACGACTCACTATAGGGCTAGCGTACCTGTGT-3’) and T7-g5611-Rev (5’-GTAATACGACTCACTATAGGGGCCAGTCCCCAGC-3’). To test for “off-target-effects”, two non-overlapping DNA fragments coding for *Pt-fgf8* were amplified using T7-Pt-fgf8-off1-Fw (5’-GTAATACGACTCACTATAGGGCTCCGCGCTGCGGC-3’) and T7-Pt-fgf8-off1-Rev (5’-GTAATACGACTCACTATAGGGCGCCTCAATAGTGGAGC-3’) and T7-Pt-fgf8-off2-Fw (5’-GTAATACGACTCACTATAGGGGTGTGTCTATTCAAAGAAGG-3’) and T7-Pt-fgf8-off2-Rev (5’-GTAATACGACTCACTATAGGGGGATGATGAGAGATCTATAG −3’), respectively. These DNA fragments were used as a template to generate dsRNA that targeted independent regions of the *Pt-fgf8* transcript. For the statistics shown in Fig. S2, four spider females were injected with dsRNA of each “off-target” fragment.

The statistics shown in Fig. S5 l (knockdown of *Pt-dof*) is based on three injected spider females.

*Pt-hh* was cloned into pCRII vector using Pt-hh-Fw (5’-GGTACACCCATAAATGCCGTCAGTTGAG-3’) and Pt-hh-Rev (5’-GTATATTCATGACAAGCGCCAGATCACACC-3’). Subsequently, T7 and T7M13R primer were used to generate the template for dsRNA production. Several spiders were injected with the generated *Pt-hh* dsRNA and only severely affected cocoons were used for the experiments shown in this study.

### Embryo fixation, *in situ* hybridisation and pMad antibody staining

To analyse BMP pathway activity in control and *Pt-fgf8* knockdown embryos a pMad antibody staining was performed as described in [19]. Fluorescent pMad antibody staining was performed using the Phospho-Smad1/5 (Ser463/465) (41D10) Rabbit mAb (Cell Signaling Technology, Inc.; antibody concentration: 1:1000) as primary and an Alexa488 anti rabbit AB (Invitrogen; 1:400 concentration) as secondary antibody.

### *In situ* hybridisation

*In situ* hybridisation was performed as described previously [19,36]. Fluorescent *in situ* hybridisation was performed using FastRed [37]. Double *in situ* hybridisation was performed using NBT/BCIP and INT/BCIP as a substrate.

### Imaging and image analysis

Embryos were imaged using an Axio Zoom.V16 that was equipped with an AxioCam 506 colour camera. Confocal imaging was performed on a LSM 700 (Zeiss). Projections of image stacks were carried out using Helicon Focus (HeliconSoft) or Fiji [38]. Live imaging was carried out on the Axio Zoom.V16 and images were processed using Fiji. Living embryos were monitored under Voltalef H10S or Halocarbon 700 (Merck) oil and live imaging was performed at RT.

Movies were created using Fiji. Images have been adjusted for brightness and contrast using Adobe Photoshop CS5.1.

False-colour overlays of *in situ* hybridization images were generated as described in [19].

## Results

### Cumulus migration requires FGF signalling

A RNAi screen, designed to find new genes that are involved in the process of axis specification in *P. tepidariorum* (see [19]) revealed that also FGF signalling might have an important role during axis formation in spiders.

During this RNAi screen, the knockdown of a gene, orthologous to *fgf8* (named *Pt-fgf8* in this study; AUGUSTUS genome prediction: aug3.g5611.t1; see Fig. S1), led to a strong dorsoventral axis patterning defect phenotype in a portion of the analysed embryos. Time lapse imaging of *Pt-fgf8* knockdown embryos revealed that the down regulation of *Pt-fgf8* is preventing cumulus migration at stage 5 of development (Fig. 1 a, b; Movie S1). In control embryos, morphogenetic processes lead to the opening of the germ-disc and to the formation of the bilaterally symmetric germ-band embryo owing a clear dorsoventral polarity (Fig. 1 a, b; Movie S1). This was in stark contrast to *Pt-fgf8* pRNAi embryos in which germ-disc opening did not occur. As a result, the knockdown embryos had a tube-like morphology at stages 7-10 (Fig. 1 a, b; Movie S1). As revealed by an *in situ* hybridisation for the ventral cell fate marker *short gastrulation* (*Pt-sog*) and the anterior marker *orthodenticle* (*Pt-otd*), stage 8 *Pt-fgf8* knockdown embryos were completely ventralized and were lacking any bilateral symmetry (Fig. 1 c-g). However, the staining of *Pt-otd* at the anterior and *caudal* (*Pt-cad*) at the posterior pole indicated that anterior and posterior patterning was not affected by the knockdown of *Pt-fgf8* (Fig. 1 g, g’ and i).

**Fig. 1.**
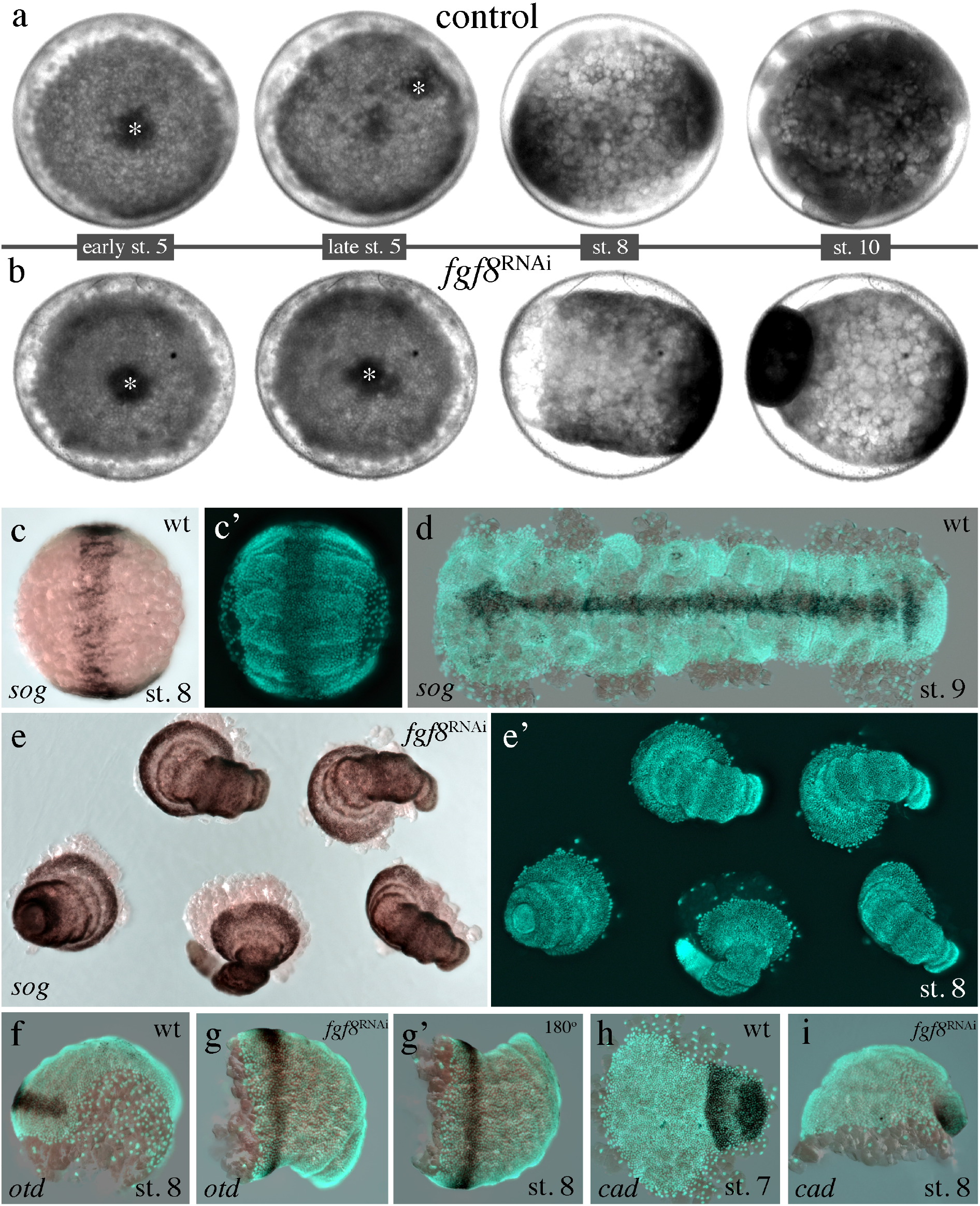
Knockdown of *Pt-fgf8* prevents cumulus migration. (**a, b**) Stills from Movie S1. In contrast to the control embryo, the cumulus (marked by the asterisk in **a** and **b**) of *Pt-fgf8* RNAi embryo is not shifting and stays in the centre of the germ-disc during stage 5 of embryonic development. *Pt-fgf8* knockdown embryos are radially symmetric, ventralized and have a tube-like morphology during later stages of development (**b, e, g, i**). (**c-i**) Expression of a ventral (*Pt-sog*), an anterior (*Pt-otd*) and a posterior (*Pt-cad*) marker gene in control and *Pt-fgf8* RNAi embryos. Whole mounted (**c, e, f, g, i**) and flat mounted (**d, h**) embryos co-stained with the nuclear dye Sytox green. To clearly show the radially symmetry, the embryo shown in g was turned by 180° (**g’**).

As already mentioned, only a portion of the *Pt-fgf8* RNAi embryos showed a DV patterning phenotype in the initial knockdown experiment. In this initial RNAi experiment almost the complete coding sequence of *Pt-fgf8* was used to generate dsRNA for parental injection. To verify and quantify this observation and to rule out that this low knockdown efficiency was fragment specific, we tested two non-overlapping fragments spanning the complete coding sequence (see Fig. S2; *fgf8*^off1-pRNAi^) as well as a large region of the 3’UTR of *Pt-fgf8* (see Fig. S2; *fgf8*^off2-pRNAi^). For both dsRNA fragments we observed a very similar number of embryos showing the described cumulus migration and DV defect phenotype (Fig. S2). The highest number of affected embryos was present in the third cocoon (around 3 weeks after injection, see Fig. S2). However, similar to the initial approach, only around 25% of the embryos were affected after the knockdown of *Pt-fgf8*. To overcome the low knockdown efficiency we were searching for other components of the FGF signalling pathway. Scanning the *Parasteatoda tepidariorum* genome [26], we found two paralogous genes coding for FGF receptors (named *Pt-FGFR1* (aug3.g26565.t1) and *Pt-FGFR2* (aug3.g26271.t1) in this study) and an ortholog of a gene known as *downstream-of-fgf* (*dof*), *stumps* (*sms*) or *heartbroken* (*hbr*) (named *Pt-dof* in this study; aug3.g4286.t1). We analysed the expression of the genes and found that *Pt-FGFR1* and *Pt-dof* were expressed in the migrating cumulus of stage 5 embryos, while *Pt-FGFR2* did not show any expression at germ-disc stage (Fig. S3-S5).

As expected from a previous analysis [22] the knockdown of the two *fgf* receptors did not lead to any detectable defect in cumulus migration. In contrast, some of the *Pt-dof* RNAi embryos showed a phenotype that was identical to the knockdown of *Pt-fgf8* (Fig. S5i). However, as the *Pt-dof* knockdown did not result in a higher frequency of strong FGF signalling dependent cumulus migration defect phenotypes (Fig. S5l), we used the *Pt-fgf8* RNAi embryos for the further analysis.

In conclusion, the knockdown of *Pt-fgf8* as well as of the downstream effector *Pt-dof* prevents cumulus migration in the spider *Parasteatoda tepidariorum*. As cumulus migration is required for the formation of the dorsoventral body axis, FGF signalling is an important player during early spider embryogenesis.

### Dynamic germ-disc expression of *Pt-fgf8*

It is known that FGF ligands can function as a chemo attractant to attract cells that express a FGF receptor (reviewed in e.g. [6]). For this reason, we were very interested to see whether *Pt-fgf8* might be expressed in a subset of germ-disc cells to be able to guide the *Pt-FGFR1* expressing cumulus cells towards the rim of the germ-disc.

At stage 3 and 4, *Pt-fgf8* transcripts were detectable within the central primary thickening of the germ-disc (Fig. 2 a-c). Confocal sectioning of fluorescently labelled stage 4 embryos revealed that *Pt-fgf8* transcripts were present in the gastrulating cells that entered the germ-disc via the blastoporus (see Fig. 2 c).

**Fig. 2.**
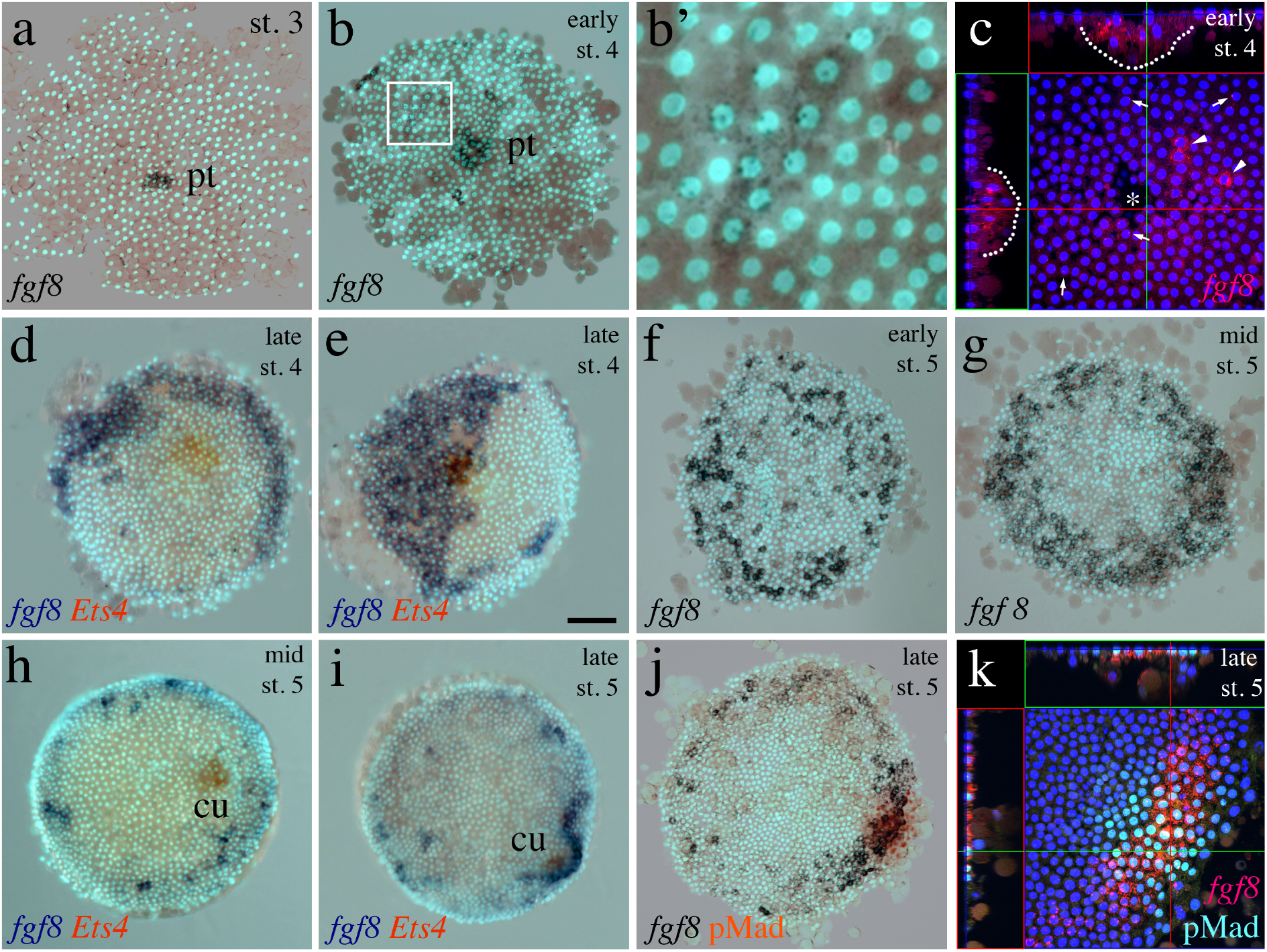
Dynamic expression of *Pt-fgf8* in the germ-disc. (**a-c**) At stage 3 and 4, cells delaminate via the blastopore (see asterisk in **c**) and form the primary thickening (pt in **a** and **b**). The cells of the primary thickening are positive for *Pt-fgf-8* transcripts (outlined by the dotted line in the orthogonal views in **c**). At early stage 4, *Pt-fgf8* is stochastically activated in single or in groups of cells within the ectoderm of the germ-disc (see boxed region in **b** and arrows and arrow heads in **c**). Nascent transcripts in the nuclei (arrow heads) and cells with *Pt-fgf8* transcripts in the cytoplasm (arrows) of the germ-disc cells are indicated in **c**. Boxed region in **b** is magnified in **b’**. (**d-g**) At late stage 4 and early stage 5, embryos show broad, asymmetric (**d, e**), spotted (**f**) or ring like (**g**) *Pt-fgf8* expression within the germ-disc. At mid and late stage 5, most embryos show strong expression of *Pt-fgf8* in the region around the migrating cumulus (cu). (**j** and **k**) Ectodermal expression of *Pt-fgf8* in the germ-disc cells overlaying the cumulus. Double *in situ* hybridisation showing *Pt-fgf8* expression in blue and the cumulus marker *Pt-Ets4* in orange (**d, e, h-j**). **c** and **k** show confocal scans of fluorescently labelled embryos (via pMad antibody staining and *in situ* hybridisation using fast red as a substrate). All embryos were co-stained with the nuclear dye DAPI or Sytox green. *Pt-fgf8* expression beyond stage 5 is depicted in Fig. S10.

Single cells or groups of ectodermal germ-disc cells started to express *Pt-fgf8* (Fig. 2 b’, Fig. 2 c) at the beginning of stage 4. At the end of stage 4 (Fig. 2 d-e, Fig. S6) many embryos showed a clear asymmetric *Pt-fgf8* expression. In extreme cases, half of the germ-disc was positive while the other half was mostly negative for *Pt-fgf8* transcripts (Fig. 2 e, Fig. S6 a, c, e, f, i). However, at this stage every embryo showed a unique pattern and the expression of *Pt-fgf8* was very dynamic and variable. While in some embryos *Pt-fgf8* could be detectable in only a few germ-disc cells (e.g. Fig. S6 b, g) other embryos showed a nearly ubiquitous expression of *Pt-fgf8* (e.g. Fig. S6 d, h). At late stage 4, expression seemed to be switched off from the primary thickening and was restricted to the ectodermal cells of the germ-disc (e.g. Fig. 2 d; e.g. Fig. S6 b, g, j).

A similar dynamic and variable expression was also detectable at early and mid stage 5 embryos when the cumulus cells started to shift towards the rim of the germ-disc. At these stages, some embryos showed little (Fig. S7 c, e, g, h) or a patchy, salt and pepper like expression of *Pt-fgf8* (Fig. 2 f, h e.g. Fig. S7 i), others had a ring-like expression domain (Fig. 2g, Fig. S7 j-m) and single embryos showed no *Pt-fgf8* expression (Fig. S7 f). Importantly, during stage 5, *Pt-fgf8* expression was always switched of from the centre of the germ-disc (Fig. 2 f-j, Fig. S7-S9).

In contrast to this dynamic and variable expression until mid stage 5, we could observe a very consistent feature of *Pt-fgf8* expression at late stage 5. In all analysed embryos (n>30) the cumulus did reach the rim of the germ-disc at high levels of *Pt-fgf8* expression (Fig. 2 i-k, Fig. S8 and S9). Confocal sectioning of fluorescently labelled embryos revealed that *Pt-fgf8* transcripts were detectable in ectodermal, pMad positive, germ-disc cells (Fig. 2 k). For several embryos, we even had the impression, that ectodermal, *Pt-fgf8* expressing cells were directly anterior to the travelling cumulus cells (e.g. Fig. 2i).

Overall, *Pt-fgf8* showed a very dynamic expression during germ-disc stages and every embryo had its unique pattern. However, the very asymmetric expression at late stage 4 and the strong expression of *Pt-fgf8* at the final position of the cumulus at late stage 5 suggested that *Pt-fgf8* might function as a chemo attractant to guide the cumulus cells towards the rim of the germ-disc.

### Possible ancestral function of FGFR-signalling in cumulus migration

Searching for fibroblast growth factor like proteins revealed that the *P. tepidariorum* genome possesses two genes that code for FGFR ligands. This is in contrast to many other spider species where we could find three FGFs. Phylogenetic analysis revealed that the spider FGFs fall into the FGF A and D families (FGF families according to [39], see Fig. S1). All analysed spider species have a clear homolog for Fgf1 and Fgf8. Only *P. tepidariorum* is missing a homolog for Fgf17, which is present in other spider transcriptomes/genomes. As the genome of *P. tepidariorum* is well assembled and several transcriptomic resources are available for this species [26,40], it is very likely, that *fgf17* was lost in the lineage leading to *Parasteatoda*.

To see whether Fgf17 might have a role during cumulus migration in other spider species, we cloned *fgf17* and analysed its expression in *Steatoda grossa* (*S. grossa*, a cobweb spider species, closely related to *P. tepidariorum*) and *Acanthoscurria geniculata* (a basally branching mygalomorph spider species). Live imaging of *S. grossa* embryos revealed, that embryogenesis is very similar to *P. tepidariorum* embryogenesis (see Movie S2). Expression analysis revealed that *S. grossa fgf17* (*Sg-fgf17*) seems not to be expressed at germ-disc stages (Fig. S11 b). This is in contrast to *Sg-fgf8* that shows a similar and variable expression (see Fig. S11a), as was shown for *Pt-fgf8* in germ-disc stage *P. tepidariorum* embryos (e.g. Fig. 2). *A. geniculata fgf8* (*Ag-fgf8*) and *fgf17* (*Ag-fgf17*) showed both a variable (sometimes enhanced expression in the region of the cumulus) expression pattern at stage 5 of embryogenesis (Fig. S12 g and h). Like in *P. tepidariorum*, also *Ag-FGFR1* (but not *Ag-FGFR2*) and *Ag-dof* showed an expression in cells of the migrating cumulus (Fig. S12 c-e). This analysis indicated that also in basally branching spiders FGF signalling might be involved in cumulus migration. For *fgf1* we could not detect any specific embryonic localisation of transcripts, neither in *P. tepidariorum* nor in *A. geniculata* (Fig. S12 f, Fig. S13 a, Fig. S14).

### Cumulus expression of FGF signalling components are regulated by the transcription factor *Pt-Ets4*

The primary thickening of stage 4 embryos is a cluster of gastrulating cells that enter the germ-disc via the blastoporus [13–15,41]. In *P. tepidariorum*, around nine cells of the primary thickening will contribute to the cell cluster of the migrating cumulus [13]. In 2017, we showed that the transcription factor *Pt-Ets4* is expressed in the cells of the primary thickening (see Fig. 3a) and in the migrating cumulus [19]. We identified *Pt-Ets4* as an important factor for maintaining the cohesion of the cumulus cells and could show that Pt-Ets4 is required to activate cumulus specific genes. Furthermore, ectopic expression of *Pt-Ets4* was able to induce cell delamination and migration [19].

**Fig. 3.**
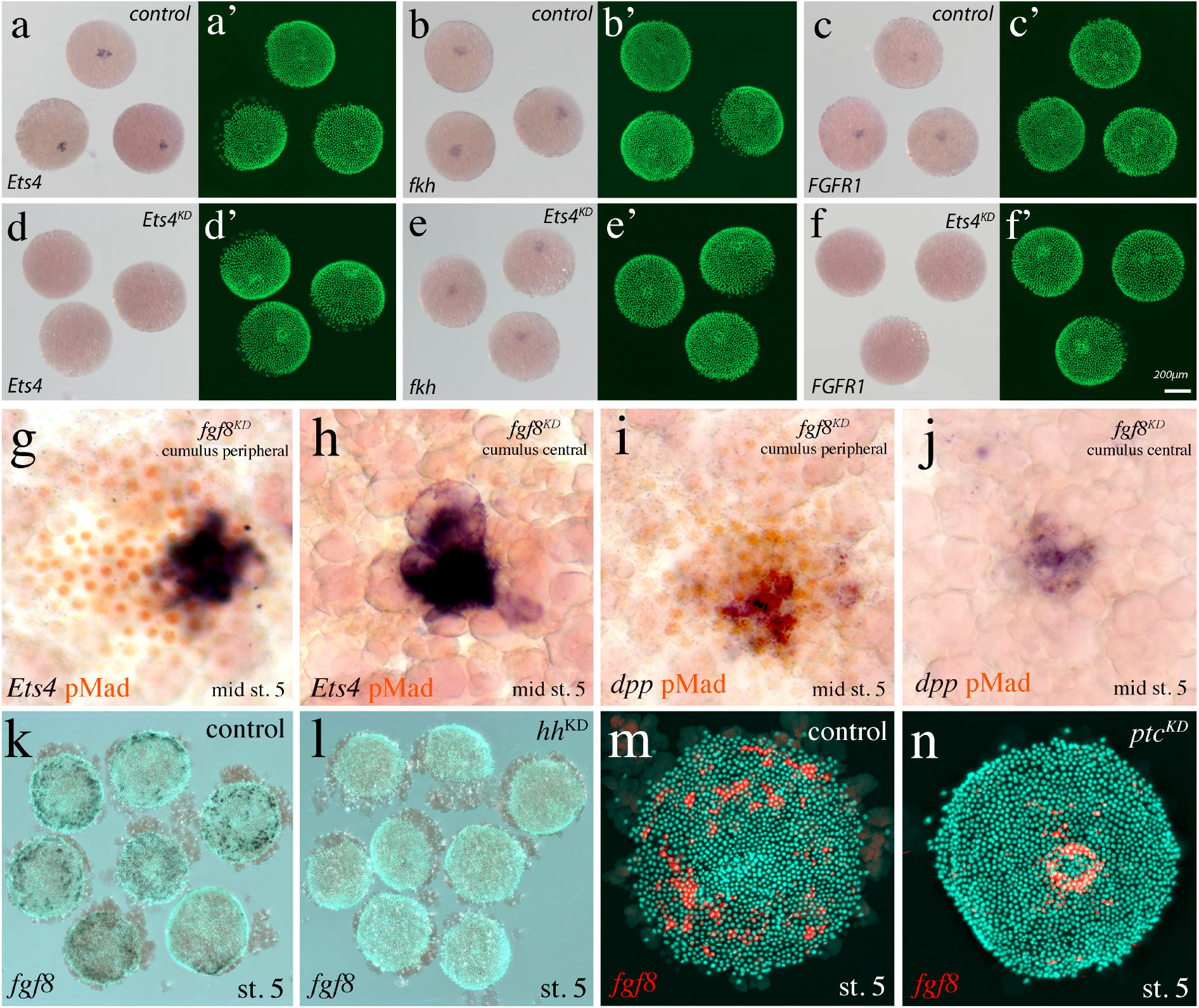
Upstream and downstream of FGF signalling. (**a-f**) FGF signalling components are regulated by *Pt-Ets4* (also compare to Table1 and Fig. S5). (**a, a’** and **d, d’**) *Pt-Ets4* expression in control and *Pt-Ets4* RNAi embryos. (**b, b’** and **e, e’**) Positive control. As shown before (Pechmann et al. 2017), *Pt-Ets4* is not influencing the expression of *Pt-fkh* in the cumulus. (**c, c’** and **f, f’**) The expression of *Pt-FGFR1* is absent in *Pt-Ets4* RNAi embryos. (**g-j**) Knockdown of *Pt-fgf8* is influencing the activation of the BMP signalling pathway. BMP signalling pathway activation visualized via pMad antibody staining (orange nuclear staining) in *Pt-fgf8* RNAi embryos. *Pt-fgf8* knockdown embryos show no obvious reduction in *Pt-Ets4* or *Pt-dpp* expression. In *Pt-fgf8* knockdown embryos that lack cumulus migration (**h, j**) also pMad activation is strongly reduced. (**k-n**) Hedgehog signalling is influencing *Pt-fgf8* expression in germ-disc embryos. *Pt-fgf8* expression is absent in *Pt-hh* RNAi embryos (**k, l**). Knockdown of the receptor *Pt-ptc* is leading to ectopic activation of Pt-*fgf8* in the centre of the germ-disc (**m, n**).

As described above, *Pt-fgf8* (Fig. 2 a-c), *Pt-FGFR1* (Fig. S3 a) and *Pt-dof* (Fig. S5 b) were expressed in the region of the primary thickening and were likely to be co-expressed with *Pt-Ets4*. To see whether *Pt-Ets4* might be involved in the transcriptional activation of the FGF signalling components we analysed the expression of *Pt-FGFR1* and *Pt-dof* in stage 4 *Pt-Ets4* RNAi embryos. Indeed, while the expression of *Pt-fkh*, a gene known being expressed in the primary thickening but not being regulated by *Pt-Ets4*, was unchanged, *Pt-FGFR1* expression was completely absent in *Pt-Ets4* RNAi embryos (Fig. 3 c, f). Also the expression of *Pt-dof* was greatly reduced upon *Pt-Ets4* knockdown (Fig. S5 j, k). In addition, a differential expression analysis of genes being regulated by *Pt-Ets4* at stage 4 confirmed that *Pt-FGFR1, Pt-dof*, and *Pt-fgf8* have a significantly reduced expression profile (see table 1).

**Table1:** Expression changes (TPM values) of FGF signalling components after RNAi with *Pt-Ets4*.

| gene | FDR | dsRed | dsRed | Ets4 RNAi | Ets4 RNAi |
| --- | --- | --- | --- | --- | --- |
| *Ets4* | 2,55E-31 | 161,599 | 196,816 | 2,20942 | 3,25887 |
| *fgf8* | 4,87E-18 | 46,3583 | 80,273 | 6,65353 | 4,35487 |
| *dof* | 1,54E-20 | 9,30084 | 6,15884 | 0,238895 | 0,094327 |
| *fgfr1* | 1,86E-13 | 9,56246 | 8,5299 | 0,82618 | 0,580557 |
| *fgfr2* | 0,64875295 | 2,48446 | 2,67093 | 2,5176 | 1,55443 |

### Knockdown of *Pt-fgf8* is influencing the activation of the BMP signalling pathway

As shown in previous studies, the down regulation of the Hedgehog receptor Patched (*Pt-ptc*) is also able to inhibit cumulus migration [19,22]. However, there is a fundamental difference between the *Pt-ptc* and the *Pt-fgf8* knockdown phenotype. In wild type embryos, the shifting and Dpp secreting cumulus leads to the activation of the BMP signalling pathway at the periphery of the germ-disc. This is leading to the formation of the dorsal field, which is required to transform the radially symmetric germ-disc into a bilaterally symmetric germ-band. The cumulus cells of the *Pt-ptc* RNAi embryos are still able to activate the BMP signalling pathway in nearby ectodermal germ-disc cells [22]. However, as the cumulus of *Pt-ptc* knockdown embryos was not shifting, the dorsal field was established in the centre of the germ-disc. An ectopic formation of the dorsal field in the centre of the germ-disc did never occur in *Pt-fgf8* RNAi embryos (Fig. 1 and Movie S1). This suggested to us that FGFR signalling is required to activate the BMP signalling pathway in stage 5 spider embryos.

To see at what level the FGFR signalling pathway was required to activate the BMP signalling pathway, we performed an *in situ* hybridisation to detect *Pt-dpp* transcripts and performed a pMad antibody staining to analyse BMP pathway activity (Fig. 3 g-j). For this experiment, we took 338 embryos of a third cocoon (produced by a spider female that was injected with *Pt-fgf8* dsRNA) and divided the embryos in four groups. We took advantage of the fact that *P. tepidariorum* embryos of the same cocoon are developing very synchronous [14,24]. This is especially true for early developmental stages, including germ-disc stage embryos (own observation). As only around 20% of the *Pt-fgf8* RNAi embryos showed a cumulus migration defect phenotype (see Fig. S2) we monitored the development of 68 living embryos (see Fig. S15). Of these 68 embryos 16% developed the typical tube like *Pt-fgf8* knockdown phenotype. We fixed the remaining embryos of this cocoon at late stage 5 to perform the aforementioned experiments. In addition to the pMad antibody staining and to the *Pt-dpp in situ* hybridisation we analysed 72 embryos for the expression of the cumulus marker *Pt-Ets4*. This staining revealed that all analysed embryos had developed a regular cumulus. However, in 19 of the 72 embryos (26%) the cumulus did stay in the centre of the germ-disc and these embryos showed a strongly reduced pMad signal (Fig. 3h). The 59 embryos that we analysed for *Pt-dpp* expression did not reveal a reduced expression for *Pt-dpp* transcripts. Again, in a small portion of the analysed embryos (23%) the cumulus did not shift towards the rim of the disc and *Pt-dpp* transcripts were detectable in the centre of the germ-disc (Fig. 3j). Similar to the embryos stained for *Pt-Ets4*, the pMad signal was strongly reduced in embryos with a centrally located cumulus. Finally, we took the remaining embryos (139 embryos; see Fig. S15) of the described *Pt-fgf8* RNAi cocoon and performed a pMad antibody staining. 20 of the 139 embryos (14%) showed a reduced pMad activity (Fig. S15). Overall, the experiment indicated that FGFR signalling is not influencing the expression of the BMP ligand *Pt-dpp* but is blocking the activation of the BMP signalling pathway downstream of Dpp.

### *Pt-fgf8* expression is regulated via the Hh signalling pathway

A genome wide search for genes being regulated by the Hh signalling pathway showed that *Pt-fgf8* expression levels were significantly altered upon *Pt-hh* and *Pt-ptc* RNAi [23]. As our experiments indicate that *Pt-fgf8* might be an important factor controlling cumulus migration, we repeated the published *Pt-ptc* and *Pt-hh* knockdown experiments and analysed the expression of *Pt-fgf8* in stage 5 embryos. Our experiments confirmed the results by Akiyama-Oda and Oda and showed that *Pt-fgf8* transcripts are no longer detectable in *Pt-hh* RNAi embryos (Fig. 3 k, l). In contrast, knockdown of *Pt-ptc* did lead to an ectopic, central activation of *Pt-fgf8* expression at stage 5 of embryonic development. As discussed below (see discussion and Fig. 4) these results are in accordance with published cumulus migration defect phenotypes that were observed in *Pt-ptc* and *Pt-hh* knockdown embryos and favour a hypothetical model in which Fgf8 acts as a chemo attractant to guide cumulus cells towards the rim of the germ-disc.

**Fig. 4.**
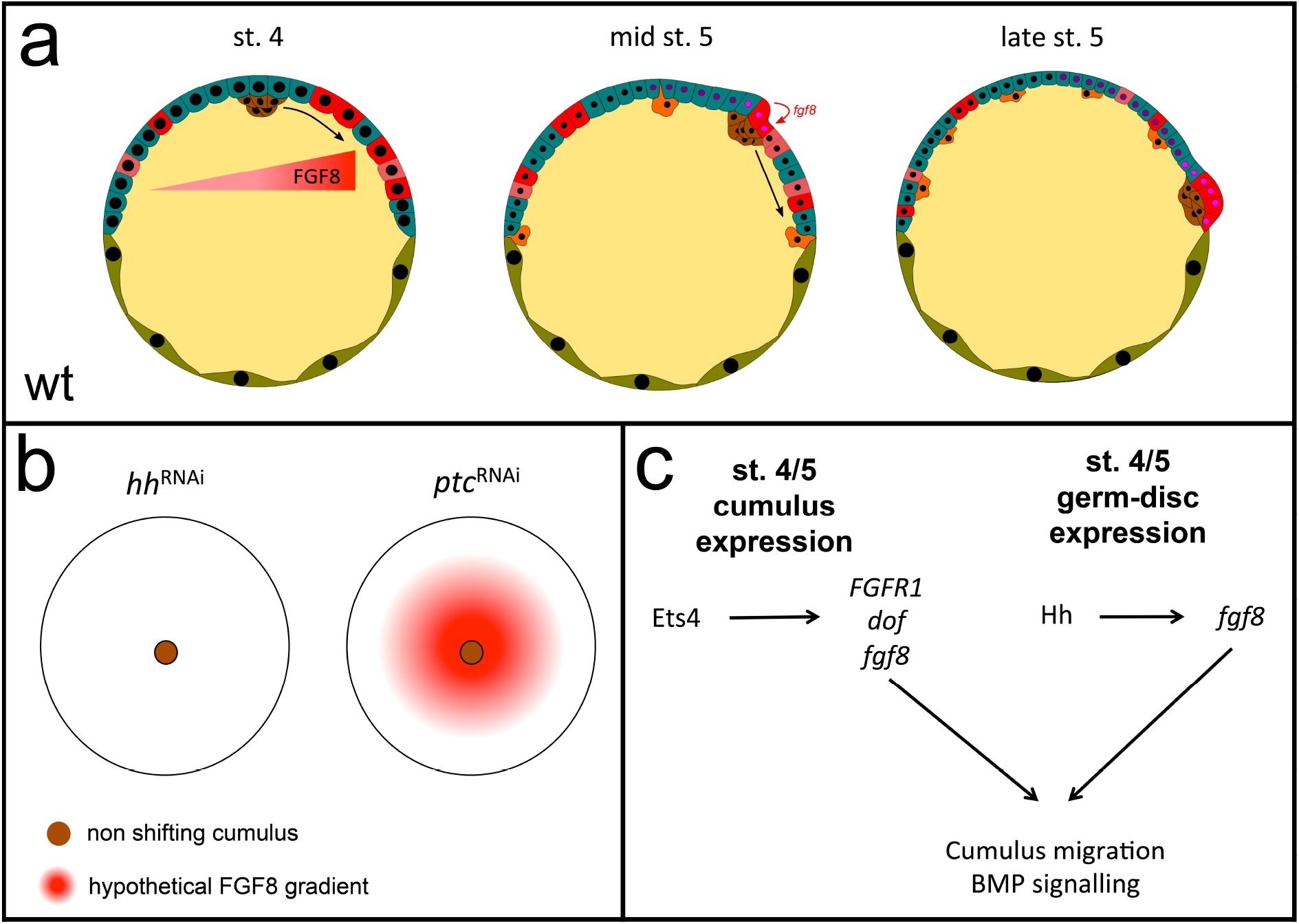
Hypothetical diagram of cumulus migration and the regulation of the FGF signalling pathway in *Parasteatoda*. (**a**) The diagram shows a cross-section of wild type germ-disc stage embryos. At early germ-disc stage (stage 4), stochastic expression of *fgf8* might result in a stochastic distribution of Fgf8 protein shortly before cumulus cells start to migrate. Cumulus cells are able to interpret the strength of this Fgf8 gradient (red graded triangle) and start to migrate towards high levels of Fgf8 (black curved arrow at st. 4). Please note: In this hypothetical model a cumulus attracting long-range FGF protein gradient is only required shortly before the cumulus starts to migrate (at st. 4). As soon as cumulus migration is initiated, the bulging of the overlaying ectoderm or another unknown mechanism might enhance *fgf8* expression (red curved arrow at mid st. 5) at the anterior border of the migrating cumulus. This process might further attract and could explain the straight movement (black arrow) of the cumulus cells towards the rim of the germ-disc. At the end of stage 5 the cumulus halts its movement at high levels of *fgf8* expression. Colour code: germ-disc cells – blue; extra-embryonic cells – green; yolk – yellow; *fgf8* expressing germ-disc cells – light and dark red; gastrulating cells – orange; cumulus cells – brown. Dark and light magenta colouring indicates former and present pMad positive nuclei of the ectoderm (the BMP signalling pathway gets activated by the Dpp secreting cumulus cells). (**b**) Top view on the germ-disc of stage 5 RNAi embryos. The brown circle indicates the cumulus that is not shifting in *hh* and *ptc* RNAi embryos. In the *ptc* RNAi embryo, hypothetical Fgf8 protein levels are indicated in red shading. In the wild-type, the cumulus is shifting straight towards the rim of the disc and stops at high levels of Fgf8 (compare to **a**). The cumulus of strongly affected *hh* and *ptc* RNAi embryos does not shift because of the missing (in *hh* RNAi, see Fig. 3l) or the ectopic, central (in *ptc* RNAi, see Fig. 3n) expression of *fgf8*. (**c**) Simplified gene regulatory network (GRN) that is responsible for the regulation of components of the FGF signalling pathway in the cumulus and in the ectoderm of the germ-disc.

## Discussion

During embryogenesis of most animals, BMP signalling is involved in setting up the dorsoventral (DV) body axis (reviewed in e.g. [42]). In spiders, the migrating cumulus is the source of this DV axis inducing BMP signal. In this arthropod system, the controlled localization of the BMP signal (via local secretion of Dpp from cumulus cells) is necessary and sufficient to induce dorsoventral body axis formation [16,18,19,21]. A failure in cumulus migration, the loss of the cumulus or the loss of the BMP signal, results in severe DV body axis patterning defects (this study, [16,18,19,22]).

Here we show, that FGF signalling is involved in cumulus migration in the spider *P. tepidariorum*. We provide evidence that FGF signalling is regulated by the transcription factor Ets4 and via the Hh signalling pathway. Finally, we show that FGF signalling is also involved in BMP signal transduction and we provide a hypothetical model in which cumulus migration might be triggered by the asymmetric localization of FGFs.

### FGF signalling is involved in cumulus migration in spiders

The observation that the knockdown of the FGFR receptors (this study, [22]) are not leading to any detectable cumulus migration phenotype and that the knockdown of the ligand *Pt-fgf8* and the downstream factor *Pt-dof* is only leading to a phenotype in a small portion of embryos indicate that this signalling pathway is not very sensitive to RNAi, at least in *P. tepidariorum*. It is very likely that small amounts of FGFR ligand as well as remnants (or maternally provided) FGF receptors are already sufficient to initiate normal cumulus migration and signalling. Nevertheless, the fact that we observed a similar FGF signalling phenotype for the ligand (*Pt-fgf8*) as well as for the downstream component (*Pt-dof*) reveals that FGF signalling has a role in cumulus migration in the spider.

Our expression analysis clearly shows that in evolutionary younger spiders like *P. tepidariorum* as well as in the basally branching spider, like the tarantula *A. geniculata*, several components of the FGF signalling pathway are expressed either in the germ-disc (*Pt-fgf8, Sg-fgf8, Ag-fgf8* and *Ag-fgf17*) or in the cumulus itself (*Pt-dof, Ag-dof, Pt-FGFR1, Ag-FGFR1*). This analysis makes it very likely, that FGF signalling is also involved in cumulus migration in basally branching spiders. Expression analysis of FGF components in other chelicerate species (e.g. Opiliones, scorpions, whip spiders, etc.) would be required to show if cumulus migration via FGF signalling is a synapomorphic character of chelicerates.

Our phylogenetic and expression analysis of FGF ligands indicates that there might be variation in the recruitment of different fibroblast growth factors during the embryogenesis of different spider species. The genome of *P. tepidariorum* is one of the best-assembled spider genomes [26]. In addition, multiple transcriptome assemblies are available for this species [40]. For these reasons, it is very likely that *fgf17* is indeed missing from the *P. tepidariorum* genome. As we could find *fgf17* in *Steatoda grossa* (a closely related cobweb spider), this potential loss of *fgf17* might be specific to *Parasteatoda* and not to cobweb spiders in general. However, as we could not detect any expression of *Sg-fgf17* during germ-disc stages, *Sg-fgf17* might not have a function that is related to cumulus migration. This is in contrast to the expression of *Ag-fgf17* in the tarantula that showed a very similar pattern to *Ag-fgf8*. These results indicate that both, Fgf8 and Fgf17, might have a role during cumulus migration in basally branching spiders like *A. geniculata* but that this function is restricted to Fgf8 in higher spiders. In turn, this might have facilitated the loss of *fgf17* in *Parasteatoda*.

### FGF signalling is controlled via Hh signalling and the transcription factor Ets4

Our results on the regulation of *Pt-fgf8* expression within the germ-disc are in agreement with the results of a recent publication, which shows the genome-wide identification of the downstream targets of the Hh signalling pathway [23]. Similar to the results of our study, also in the study by Akiyama-Oda and Oda, *Pt-fgf8* was no longer detectable in the cells of the germ-disc, when *Pt-hh* was reduced via parental RNAi. Furthermore, like in our study, *Pt-fgf8* was ectopically activated in the central region of the germ-disc upon down regulation of the negative regulator of the Hh signalling pathway *Pt-ptc* (Fig. 3k-n).

Previously, we identified the transcription factor Ets4 as an important player for dorsoventral axis formation in *P. tepidariorum* [19]. On the one hand, down-regulation of *Pt-Ets4* did lead to a loss of the integrity of the cumulus cells. As a consequence the cumulus got lost and DV body axis formation was not initiated. On the other hand, ectopic expression of Pt-Ets4 did lead to ectopic cell delamination and to an independent initiation of cell-migration of single cells. These results indicated that Pt-Ets4 is an important player within the cumulus but is not sufficient on its own to induce the formation of an ectopic cumulus. It is very likely that the combination of several molecular factors is required to set up a fully functional cumulus. To get a better idea of how Pt-Ets4 is regulating different processes within the cells of the cumulus, we performed an RNA-seq experiment and compared the expression profile of *Pt-Ets4* RNAi cumuli with those of cumuli of control embryos. This analysis clearly showed that the early activation of several components of the FGF signalling pathway within the developing cumulus cells seems to be under the control of Pt-Ets4 (Table 1). As FGF signalling is clearly involved in cell migration, this analysis indicates a link between Pt-Ets4 and cumulus cell migration.

### A hypothetical model for cumulus cell migration in spiders

During early spider embryogenesis the cumulus shifts from the centre to the periphery of the radially symmetric germ-disc. It is still unclear if this is a completely stochastic process or if the cumulus is directed towards the rim of the disc via an unknown mechanism. Life imaging of cumulus migration in different spider species indicates that the movement of the cumulus towards the rim of the disc is relatively straight, once the cumulus has started to migrate (see Movie S1 and 2, e.g. [22,24,43,44]). This holds even true for very huge tarantula embryos [18]. It was noted that in around 25% of wild type *P. tepidariorum* embryos, the cumulus is not perfectly centred within the germ-disc but is slightly asymmetrically positioned. However, even in these embryos the cumulus movement seems to not correlate with the positioning of the cumulus within the germ-disc and cumulus cells do not necessarily take the shortest distance to reach the rim of the disc [13]. This observation indicates that an unknown mechanism might be responsible to give a direction to the shifting cumulus cells.

Although the expression of *Pt-fgf8* is highly variable in germ-disc stage *P. tepidariorum* embryos, we would like to hypothesise that the stochastic and often very asymmetric expression of *Pt-fgf8* during stage 4 embryos (before cumulus migration, e.g. Fig. 2 d, e) might set up a short-term concentration gradient of Fgf8 protein across the early stage 4 germ-disc (Fig. 4a). This FGF gradient might be interpreted by the cumulus cells and could be responsible for the initial directional movement of the cumulus cells. Depending on the strength of this asymmetric or more uniform *Pt-fgf8* expression the hypothetical Fgf8 protein gradient could either be relatively steep or be more flat.

Studies in other animal model systems were able to show that FGFs can act as a chemo-attractant (e.g. [7,8], reviewed in [6]). Our observation that *P. tepidariorum* cumulus cells, at late stage 5, always stop their migration at high levels of *Pt-fgf8* expression (Fig. 2i, j, k, Fig. S8 and S9) indicates that also in the spider Fgf8 might act as a chemo-attractant.

This hypothesis is also supported by the differences in *Pt-fgf8* expression that can be found when the expression of different components of the Hh signalling pathway are reduced via RNAi. As indicated in Fig. 4b, these differences in *Pt-fgf8* expression and the resulting putative differences in the Fgf8 protein gradients, might explain the observed alterations in cumulus migration (this study, [22,23]). Briefly, in strongly affected *Pt-hh* as well as in *Pt-ptc* RNAi embryos cumulus migration was prevented [22]. This loss of cumulus migration could be explained by the loss (in *Pt-hh* RNAi) or the ectopic central activation (in *Pt-ptc* RNAi) of *Pt-fgf8* expression. In the case of the loss of *Pt-fgf8* expression in *Pt-hh* RNAi embryos, cumulus attraction via *Pt-fgf8* might be missing. As a result, cumulus migration is not initiated. In contrast, the ectopic central *Pt-fgf8* activation in *Pt-ptc* RNAi embryos might strongly attract and keep the cumulus cells in the middle of the germ-disc (Fig. 4b). Our expression analysis indicates that the clearance of *Pt-fgf8* transcripts from the centre of the stage 5 germ-disc, is an important aspect of the dynamic *Pt-fgf8* expression. This observation further supports the idea that the central ectopic expression of *Pt-fgf8* in *Pt-ptc* knockdown embryos prevents the cumulus from shifting.

In *P. tepidariorum*, Hh signalling is heavily involved in patterning the anteroposterior axis of the germ-disc [22,23]. This leads to the suggestion that Hh signalling might set up a positional value gradient across the germ-disc and that the cumulus cells move down along this emerging positional value gradient [22]. In this study it was also noted that Hh protein in vertebrate embryos might be able to travel a distance of around 300 µm and that this distance is very similar to the radius of *P. tepidariorum* germ-disc stage embryos [22,45]. However, the size of spider eggs and embryos is very diverse. In *A. geniculata* the radius of the germ-disc is already around 1 mm [18] and the embryos of this species are not the biggest amongst Mygalomorphae. For this reason we propose a different mechanism in which the cumulus on its own is driving the activation of a chemo-attractant in a posterior (centre of the germ-disc) to anterior (rim of the germ-disc) direction. This model is based on the observation that we could detect strong *Pt-fgf8* expression at the anterior border of the bulged germ-disc ectoderm (e.g. Fig. 2 i, several embryos in Fig. S8). The bulging is a consequence of the moving cumulus, which shifts underneath the germ-disc and is mechanically pushing the ectoderm upwards (see Fig. 4a, [22]). In our hypothetical model this cumulus driven bulging induces mechanical stress in the overlaying ectoderm, which in turn could trigger the further activation of *fgf8* expression in front of the migrating cumulus. It was shown that mechanically induced stretching or compression of cells is able to induce receptor activation and gene expression, including fibroblast growth factor receptors and ligands (e.g. [46,47]).

As our model is a kind of self-enhancing system and does not depend on a stable and long lasting long-range morphogen gradient, it might be functional in any spider embryo, regardless of the size of the germ-disc.

Alternative mechanisms could also lead to the local enhancement of FGFs within certain regions of the germ-disc, and this might then lead to the local attraction of cumulus cells. In zebrafish lateral line formation, it was shown that apical constriction leads to the formation of microlumen, which are in the centre of rosette-like structures. These structures are responsible to trap secreted FGFs and are able to increase signalling responses and the coordination of migratory cell behaviour (reviewed in [48], original article [49]). Future studies are required to proof if our model is valuable.

### Future directions

Our results indicate that *Pt-fgf8* is also required to activate the BMP signalling pathway in ectodermal germ-disc cells overlaying the cells of the cumulus. In wild-type, this activation is crucial to induce the dorsal field at the rim of the disc. This in turn is required to break the radially symmetry of the germ-disc (e.g. [15,16]).

The performed analysis indicates that the involvement of Pt-Fgf8 in activating the BMP signalling pathway is probably not on the level of ligand (*Pt-dpp* expression was not affected) activation but might be downstream of receptor activation. Future studies, like a genome wide search of FGFR target genes might help to understand, if FGFs are really required to activate the BMP signalling pathway or if other mechanism are causing the observed phenotype. In *Drosophila* it was shown that FGF signalling is involved in cytoneme formation (e.g. [50]). Also in *P. tepidariorum* germ-disc embryos cytoplasmic cytoneme like structures that were projecting from the cumulus towards the germ-disc epithelium could be identified [13]. Cytonemes are known to be important players in the BMP signalling pathway [51–53]. For this reason, missing cytonemes might be the reason that strongly affected *fgf8* knockdown embryos are lacking an active BMP signalling pathway.

Unfortunately, we failed to regionally down-regulate (via embryonic RNAi) or over-activate *Pt-fgf8* expression (via embryonic capped mRNA injection) in *P. tepidariorum*. In the future, also cell culture experiments could be useful to analyse if Fgf8 is indeed able to attract cumulus cells.

Recently, BMP soaked agarose beads were successfully transplanted to germ-disc stage tarantula embryos [18]. These BMP beads were able to induce a secondary axis in *A. geniculata*. Similar experiments should also be performed with Fgf8/17 soaked beads.

## Supporting information

Wang et al_Supplementary information

Wang et al_Movie_S1

Wang et al_Movie_S2

## Acknowledgements

We thank Siegfried Roth for commenting on the manuscript.

## Disclaimer

The funders had no role in study design, data collection and analysis, decision to publish or preparation of the manuscript.

## Authors’ contributions

M.P. designed the study, performed RNAi experiments and the phylogenetic analysis and wrote the paper. R.W. performed the RNA sequencing experiment, performed parts of the molecular and bioinformatics analysis and commented on the manuscript. P.S. performed the initial bioinformatics analysis of the RNA seq. data and commented on the manuscript. L.K. performed *in situ* analysis in *A. geniculata* and commented on the manuscript. All authors gave final approval for publication.

