## Supplementary material for "FGF signalling is involved in cumulus migration in the common house spider *Parasteatoda tepidariorum*": Wang et al_Supplementary information

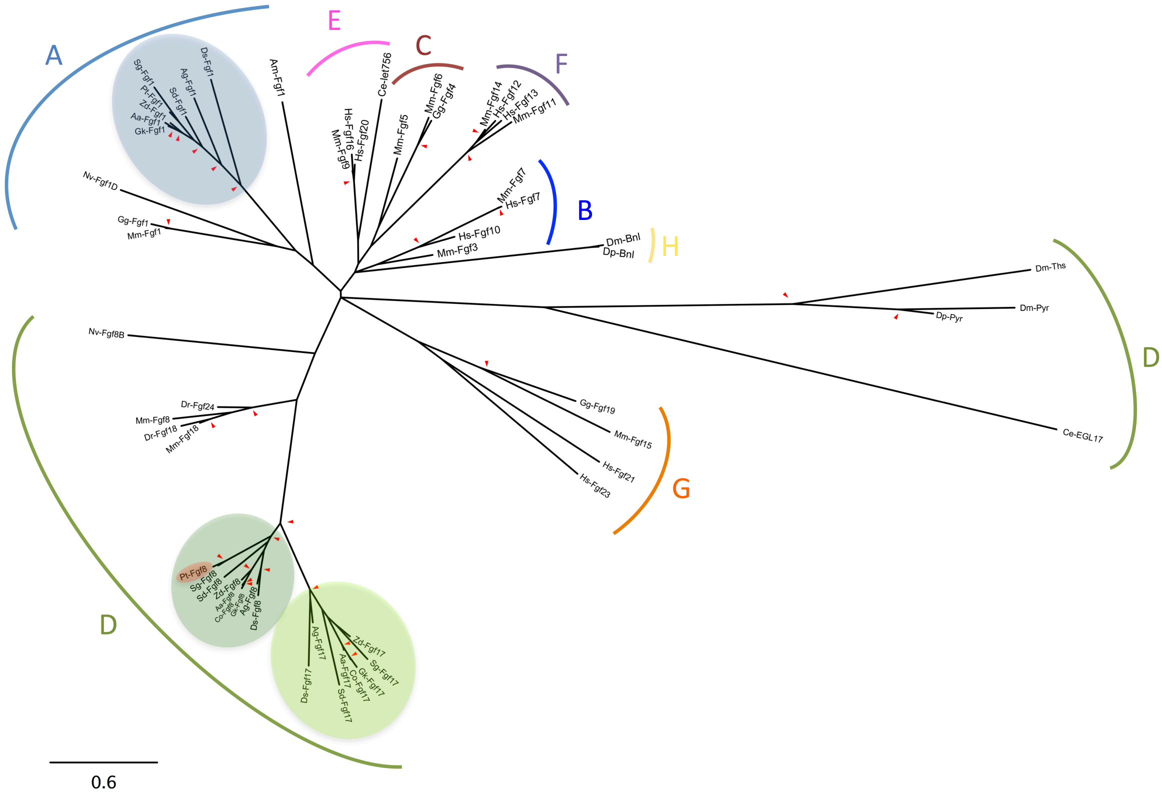


**Figure S1. Fgf Phylogeny.** The eight FGF families (A-H) were labelled according to Popovici et al. 2005. Red arrowheads mark nodes with bootstrap support greater than 800. Scale is substitutions per site. Spider Fgfs are highlighted in green and blue. Pt-Fgf8 is highlighted in red.

Species name abbreviation: Aa: *Argiope aemula*; Ag: *Acanthoscurria geniculata*; Am: *Apis mellifera*; Ce: *Caenorhabditis elegans*; Co: *Cyclosa octotuberculata*; Dm: *Drosophila melanogaster*; Dp: *Drosophila pseudoobscura*; Dr: *Danio rerio*; Ds: *Dysdera silvatica*; Gg: *Gallus gallus*; Gk: *Gasteracantha kuhli*; Hs: *Homo sapiens*; Mm: *Mus musculus*; Nv: *Nematostella vectiensis*; Pt: *Parasteatoda tepidariorum*; Sg: *Steatoda grossa*; Sd: *Stegodyphus dumicola*; Zd: *Zygiella dispar*.

Accession numbers: Dp-Pyr XP_001361810; Dp-Bnl XP_001360050; Dm-Bnl AAC47427; Dm-Ths AY553965; Dm-Pyr AY553964, Ce-EGL17 AAD00574; Mm-Fgf18 NP_032031;

Mm-Fgf 5 AAH71227; Dr-Fgf18 NP_001013282; Mm-Fgf 1 AAD00574; Dr-Fgf24 NP_878291; Gg-Fgf 1 NP_990511; Hs-Fgf10 O15520; Hs-Fgf7 NP_002000; Ce-let-756 Q11184; Mm-Fgf8 NP_034335; Nv-Fgf8B ABN70837; Am-Fgf1 ; Mm-Fgf1 AAH37601.1; Pt-Fgf1 XP_015927758.1; Pt-Fgf8 XP_015925960.1; Pt-Fgf1 XP_015927758.1; Zd-Fgf1 comp23447; Sg-Fgf1 see supplementary sequences; Gk-Fgf1 comp38018; Aa-Fgf1 comp11974_seq0; Sd-Fgf1 XP_035231158.1 ; Ag-Fgf1 TRINITY_74449; Ds-Fgf1 TRINITY_GG_25065; Pt-Fgf8 XP_015925960.1; Zd-Fgf8 comp16798; Sg-Fgf8 see supplementary sequences; Gk-Fgf8 comp16539; Co-Fgf8 comp6270_seq1; Aa-Fgf8 comp12173_seq0; Sd-Fgf8 XP_035208424.1 ; Ag-Fgf8 TRINITY_DN82690; Ds-Fgf8 TRINITY_GG_36900_c0_g1_i1; Zd-Fgf17 comp12612; Sg-Fgf17 see supplementary sequences; Gk-Fgf17 comp16435; Co-Fgf17 comp2940_seq0; Aa-Fgf17 comp5191_seq0; Sd-Fgf17 XP_035218510.1; Ag-Fgf17 TRINITY_DN82175; Ds-Fgf17 TRINITY_GG_24745_c0_g1_i1; Dr-Fgf24 NP_878291; Mm-Fgf8 NP_034335; Dr-Fgf18 NP_001013282; Mm-Fgf18 NP_032031; Nv-Fgf8B ABN70837; Mm-Fgf1 AAH37601.1; Gg-Fgf1 NP_990511; Nv-Fgf1D ABN70834.1; Am-Fgf1 XP_006571081.1; Mm-Fgf9 NP_038546.2; Hs-Fgf16 NP_003859.1; Hs-Fgf20 NP_062825.1; Ce-let-756 Q11184; Mm-Fgf5 AAH71227; Mm-Fgf6 CAA35925.1; Gg-Fgf4 AAA58706.1; Mm-Fgf14 NP_034331.2 , Hs-Fgf12 NP_066360.1; Hs-Fgf13 NP_004105.1; Mm-Fgf11 NP_001349553.1; Mm-Fgf7 EDL28153.1; Hs-Fgf10 O15520; Hs-Fgf7 NP_002000; Mm-Fgf3 AAI17062.1; Dm-Bnl AAC47427; Dp-Bnl XP_001360050; Dm-Ths AY553965; Dm-Pyr AY553964, Dp-Pyr XP_001361810; Ce-EGL17 AAD00574; Gg-Fgf19 NP_990005.2; Mm-Fgf15 AAO13811.1 ; Hs-Fgf21 AAQ89444.1; Hs-Fgf23 AAG09917.1. Several spider sequences can be found in the suppl. sequences (see below).

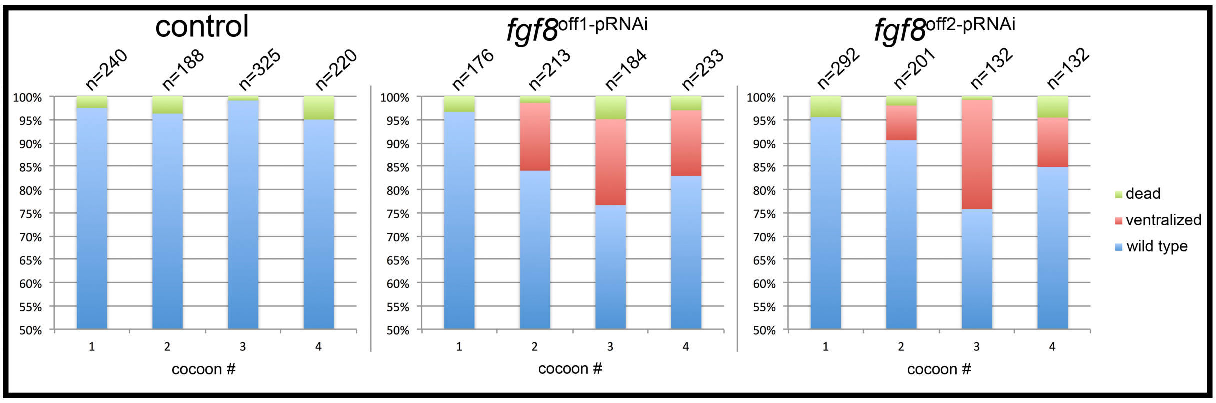


**Figure S2. Efficiency of the *Pt-fgf8* knockdown.** To check for "off-target" effects, four spider females have been injected with dsRNA of two non-overlapping *Pt-fgf8* fragments (off1 and off2), each. Embryos were monitored under oil. Knockdown efficiency is around 25% in cocoon #3.

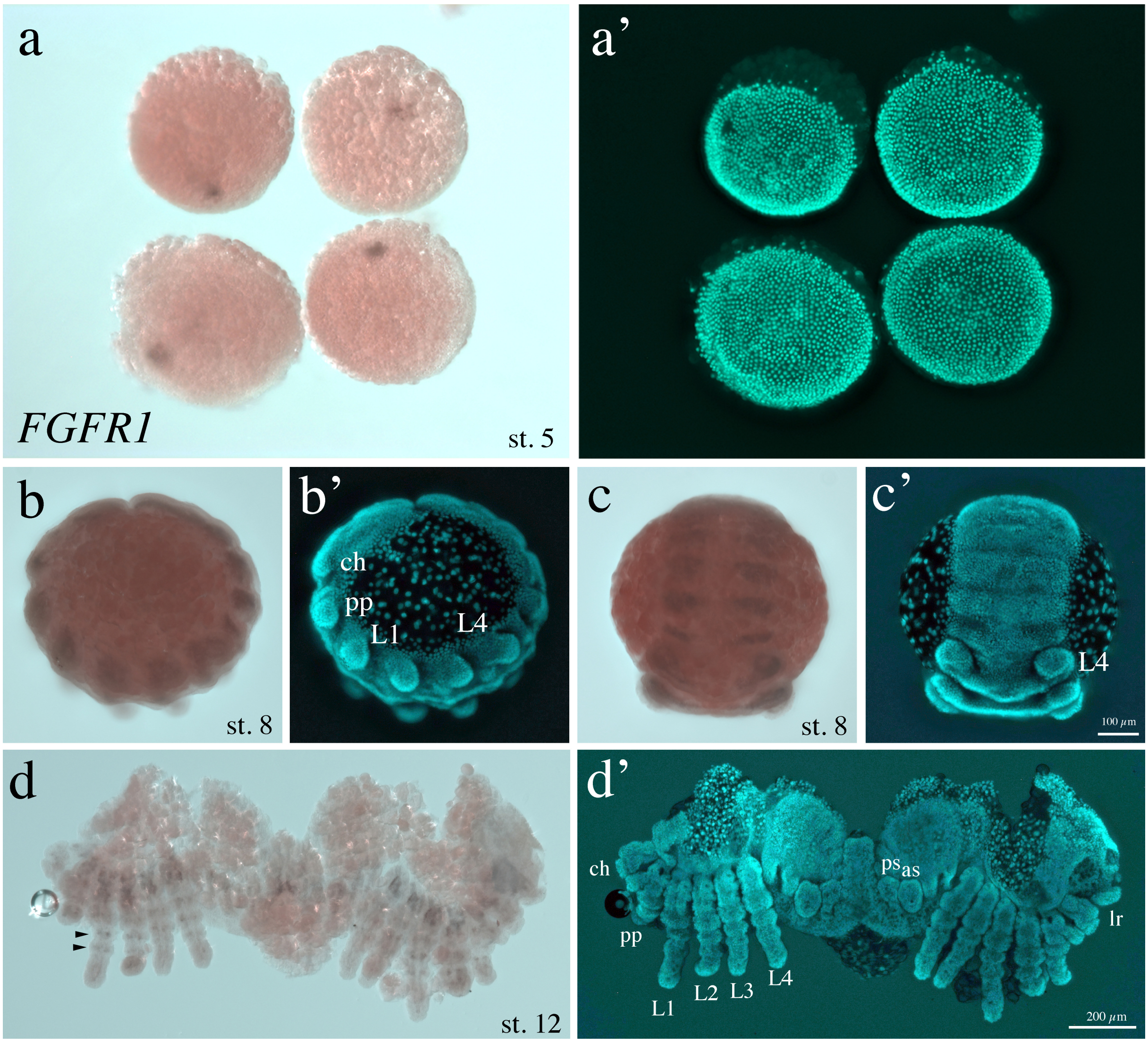


**Figure S3. Expression of *Pt-FGFR1* (st. 5 - 12).** *Pt-FGFR1* is expressed in the cumulus of stage 5 embryos (**a**) and in segmental mesodermal blocks at stage 8 of development (**b**-**c**). Arrowheads in **d** indicate faint mesodermal expression rings of *Pt-FGFR1* in the appendages of stage 12 embryos. Abbreviations: lr – labrum, ch – chelicera, pp – pedipalp, L1-L4 – walking legs 1-4, as – anterior spinneret, ps – posterior spinneret

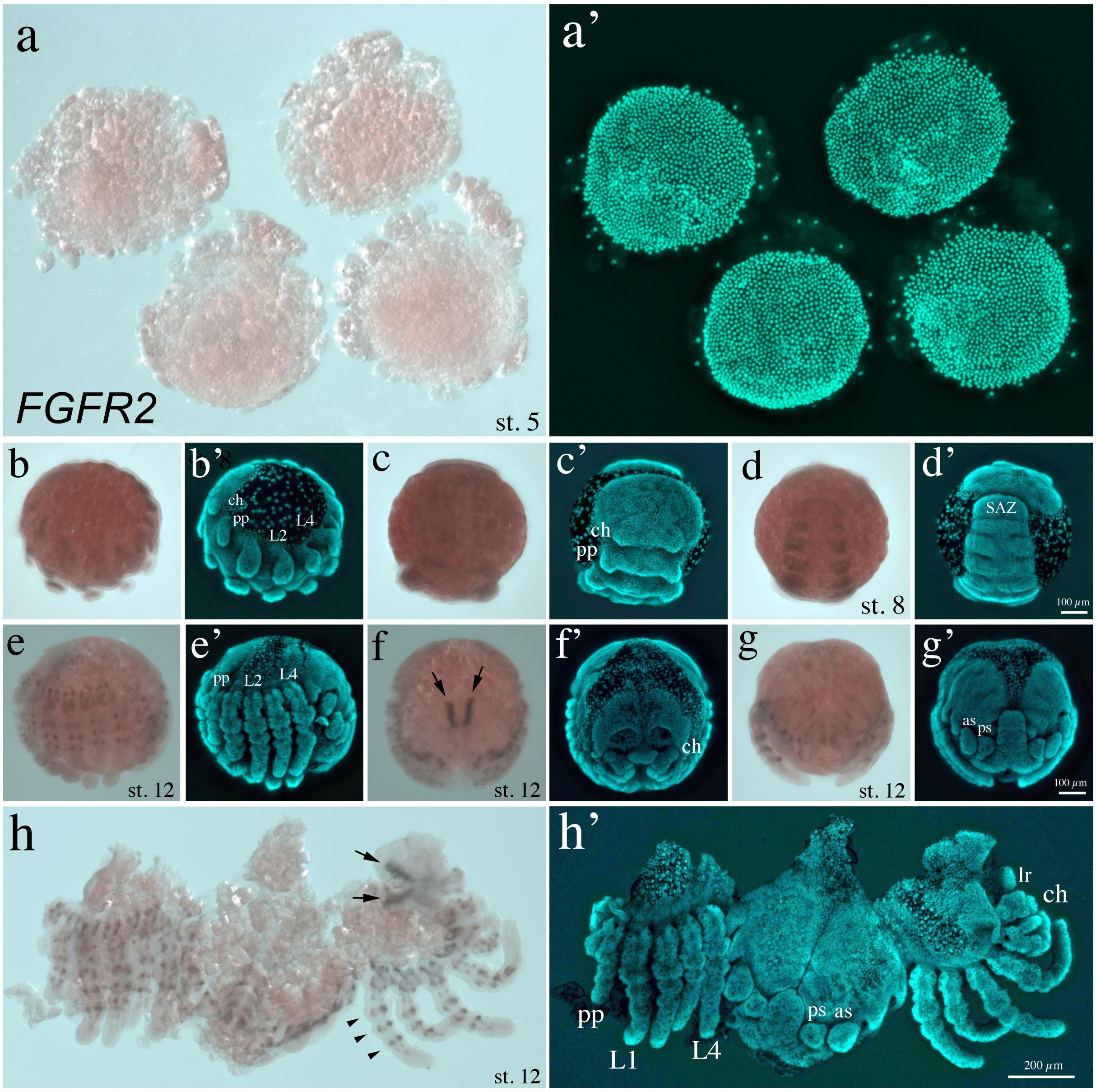


**Figure S4. Expression of *Pt-FGFR2* (st. 5 - 12).** *Pt-FGFR2* is not expressed in the cumulus of stage 5 embryos (**a**). Similar to *Pt-FGFR1* also *Pt-FGFR2* is expressed in segmental mesodermal blocks at stage 8 of development (**b-d**). *Pt-FGFR2* is expressed in two anterior domains within the developing nervous system (arrows in **f** and **h**). Arrowheads in **h** indicate strong mesodermal expression rings of *Pt-FGFR1* in the appendages of stage 12 embryos. Abbreviations: lr – labrum, ch – chelicera, pp – pedipalp, L1-L4 – walking legs 1-4, as – anterior spinneret, ps – posterior spinneret, SAZ – segment addition zone

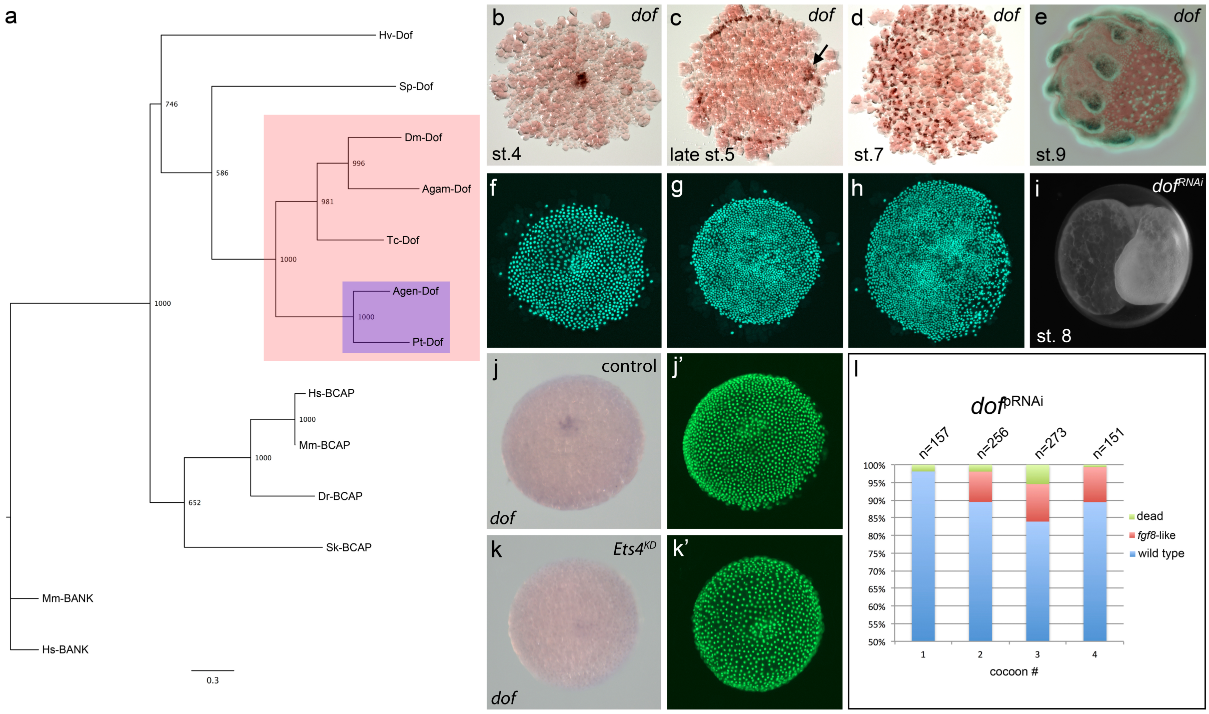


**Figure S5. Analysis of *Pt-dof*.** (**a**) Dof phylogeny. *Acanthoscurria* and *Parasteatoda* Dof (highlighted in blue) are branching with other arthropod (highlighted in red) Dof proteins. Species name abbreviation: Agam: *Anopheles gambiae*; Agen: *Acanthoscurria geniculata*; Dm: *Drosophila melanogaster*; Dr: *Danio rerio*; Hs: *Homo sapiens*; Hv: *Hydra vulgaris* Mm: *Mus musculus*; Pt: *Parasteatoda tepidariorum*; Sk: *Saccoglossus kowalevskii*; Sp: *Strongylocentrotus purpuratus*; Tc: *Tribolium castaneum*.

Accession numbers: Agam-Dof CAD27761.1; Agen-Dof TRINITY_DN82175; Dm-Dof CAA09298.1; Dr-BCAP XP_005156973.1; Hs-BANK BAB79255.1; Hs-BCAP NP_689522.2; Hv-Dof XP_012553666.1; Mm-BANK NP_001028522.2; Mm-BCAP NP_113553.1; Pt-Dof aug3.g4286.t1; Sk-BCAP XP_006814050.1; Sp-Dof XP_011666053.1; Tc-Dof XP_969604.2.

(**b-e**) Expression of *Pt-dof* in stage 4-9 embryos. *Pt-dof* is expressed in the primary thickening (**b**) and in the migrating cumulus (arrow in **c**). At late stage 5 a ring of *Pt-dof* (probably gastrulating, mesodermal cells) expression is detectable in the anterior (outer rim) of the germ-disc. *Pt-dof* expression in single cells (probably mesodermal cells; at st. 7) and in the mesoderm of the appendages (**e**). (**f-h**) Fluorescence nuclear Sytox green staining of the embryos shown in **b-d**. Bright field/Sytox green overlay (**e**). (**i**) The knockdown of *Pt-dof* is very similar to the knockdown of *Pt-fgf8*. (**j-k**) *Pt-dof* seems to be slightly down regulated in *Pt-Ets4* pRNAi embryos. (**l**) Similar to the knockdown of *Pt-fgf8*, also the knockdown of the downstream factor *Pt-dof* results in a low number of knockdown embryos that show the typical tube like (*fgf8*-like) phenotype.

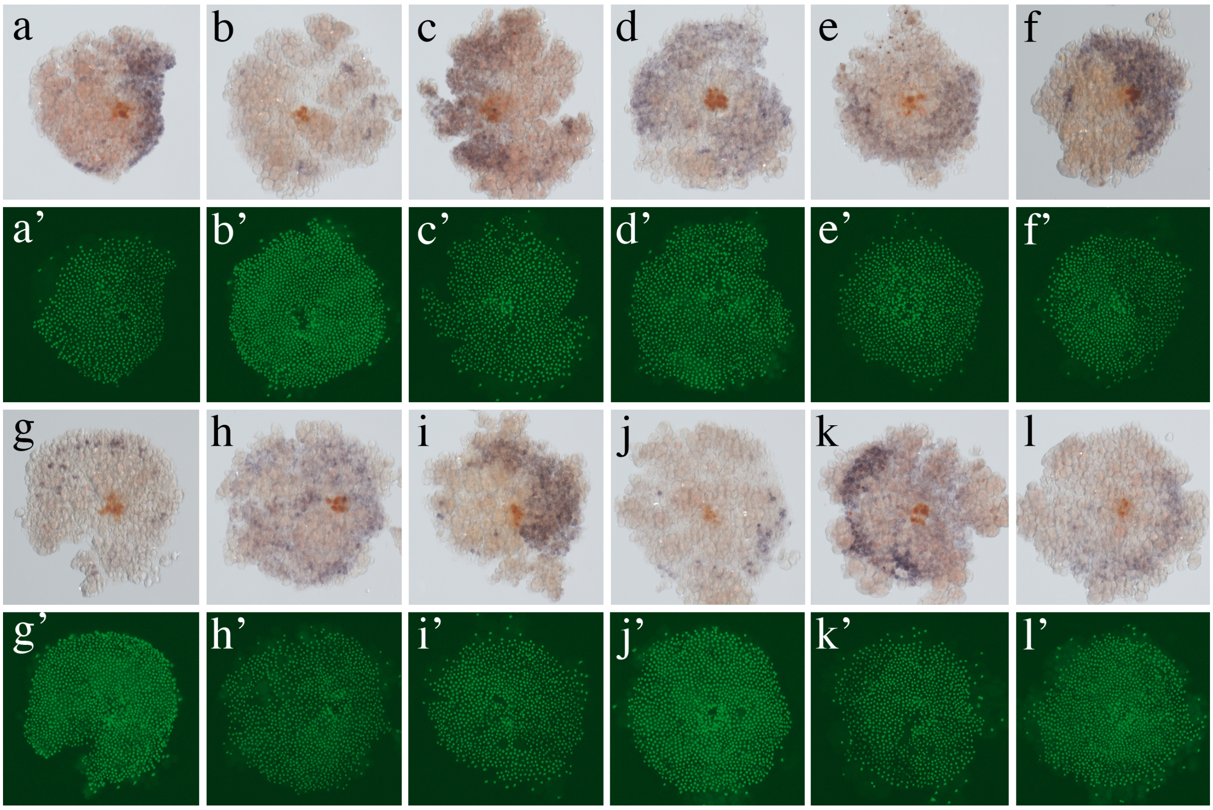

**Figure S6. Examples of *Pt-fgf8* expression at late stage 4.** Flat mounted late stage 4 embryos co-stained with the nuclear dye Sytox green.

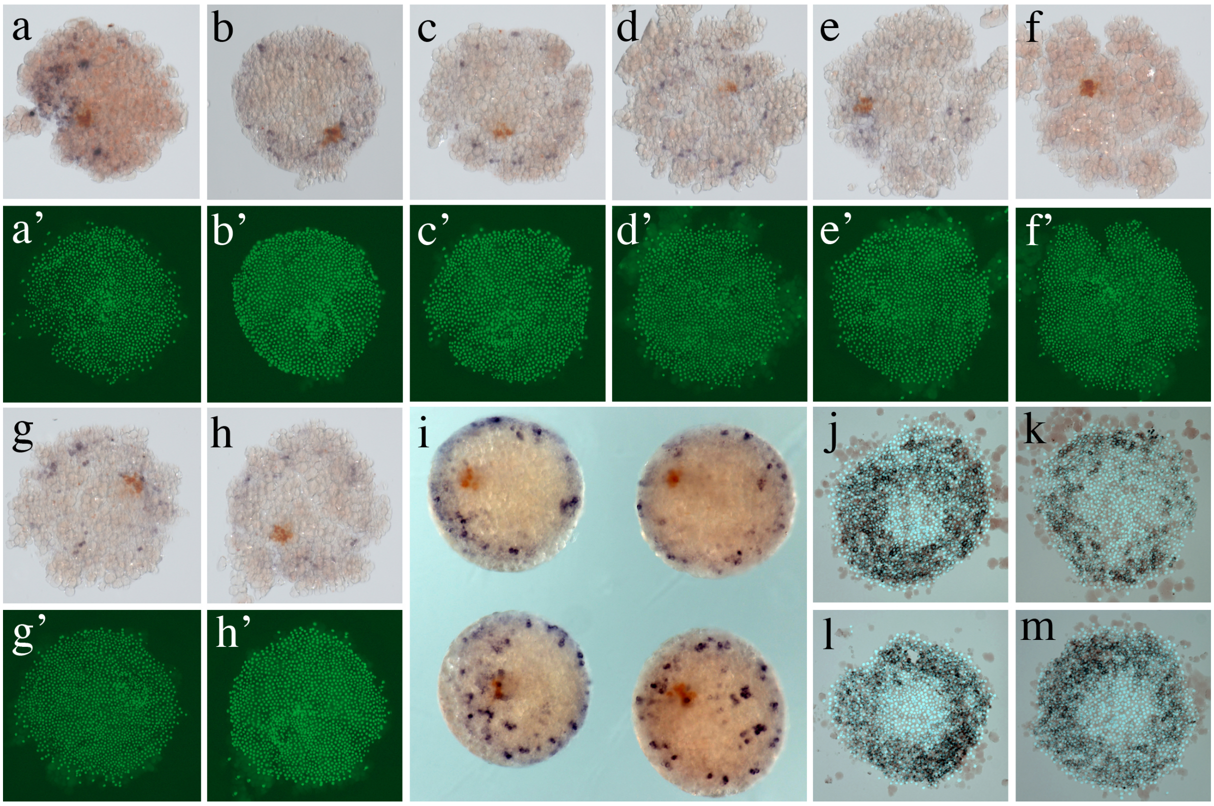

**Figure S7. Examples of *Pt-fgf8* expression at mid stage 5.** (**a-h**) Flat mounted mid stage 5 embryos co-stained with the nuclear dye Sytox green. Double *in situ* hybridisation (*Pt-fgf8* blue; cumulus marker *Pt-Ets4* in orange). (**i**) Whole mount embryos stained for *Pt-fgf8* (blue) and the cumulus marker *Pt-Ets4* (orange). (**j-m**) More examples (embryos of a cocoon from a different spider female) of *Pt-fgf8* expression at mid stage 5. Flat mounted embryos stained for *Pt-fgf8*. Embryos show a ring like expression of *Pt-fgf8*. Bright field/Sytox green overlay.

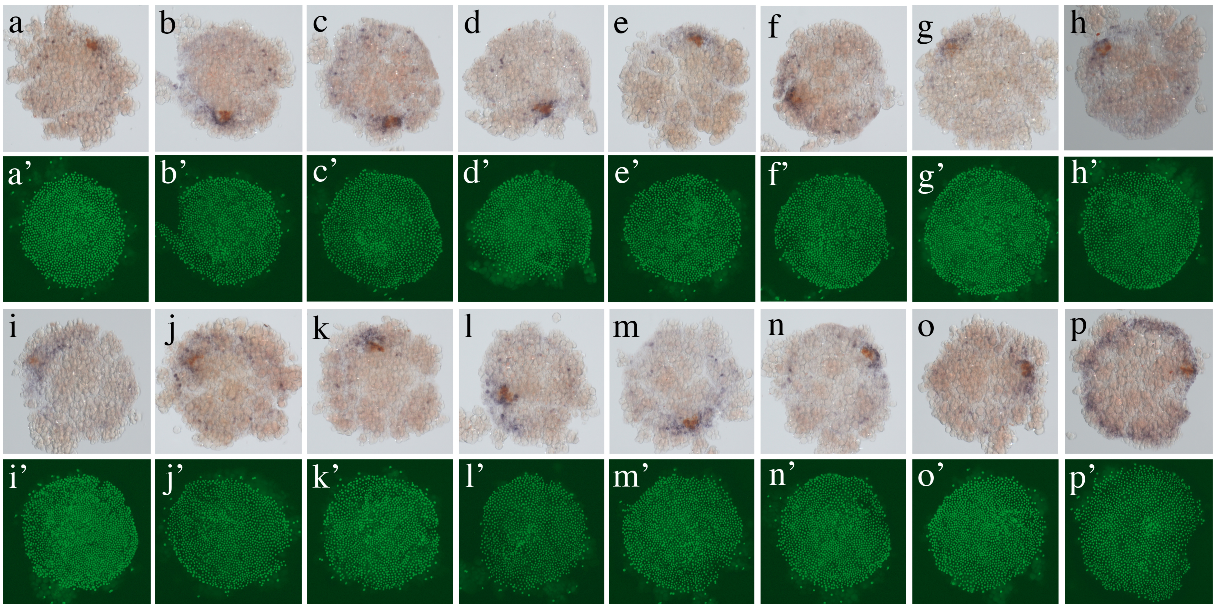


**Figure S8. Examples of *Pt-fgf8* expression at late stage 5.** Flat mounted late stage 5 embryos co-stained with the nuclear dye Sytox green. Double *in situ* hybridisation (*Pt-fgf8* blue; cumulus marker *Pt-Ets4* in orange).

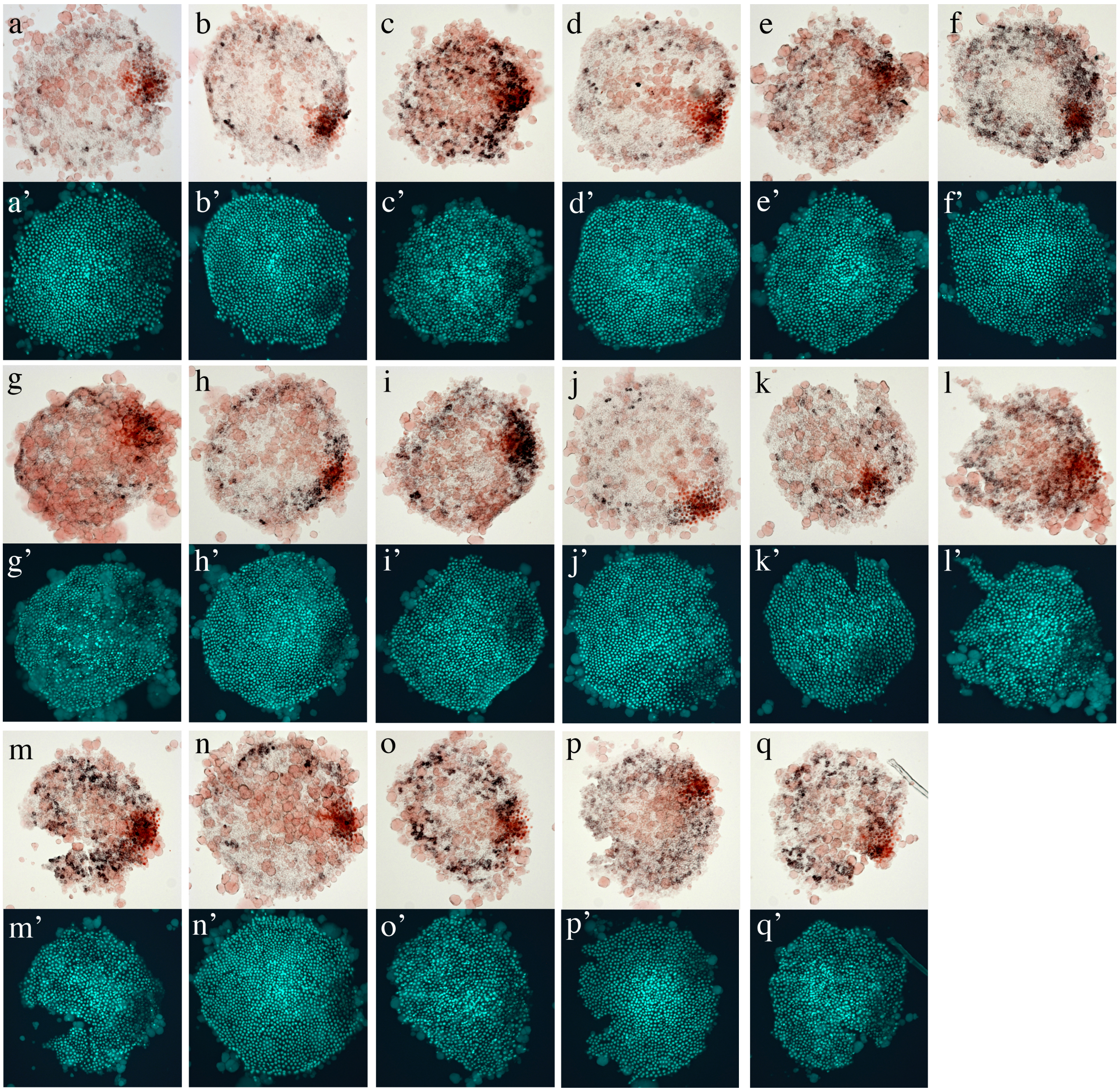


**Figure S9. Examples of *Pt-fgf8* expression at late stage 5 (pMad and *Pt-fgf8* double staining).** Flat mounted late stage 5 embryos co-stained with the nuclear dye Sytox green. Double staining (*Pt-fgf8* black; pMad antibody staining in orange). The pMad antibody staining marks the position of the cumulus and shows that the cumulus reaches the rim of the disc at high levels of *Pt-fgf8* expression.

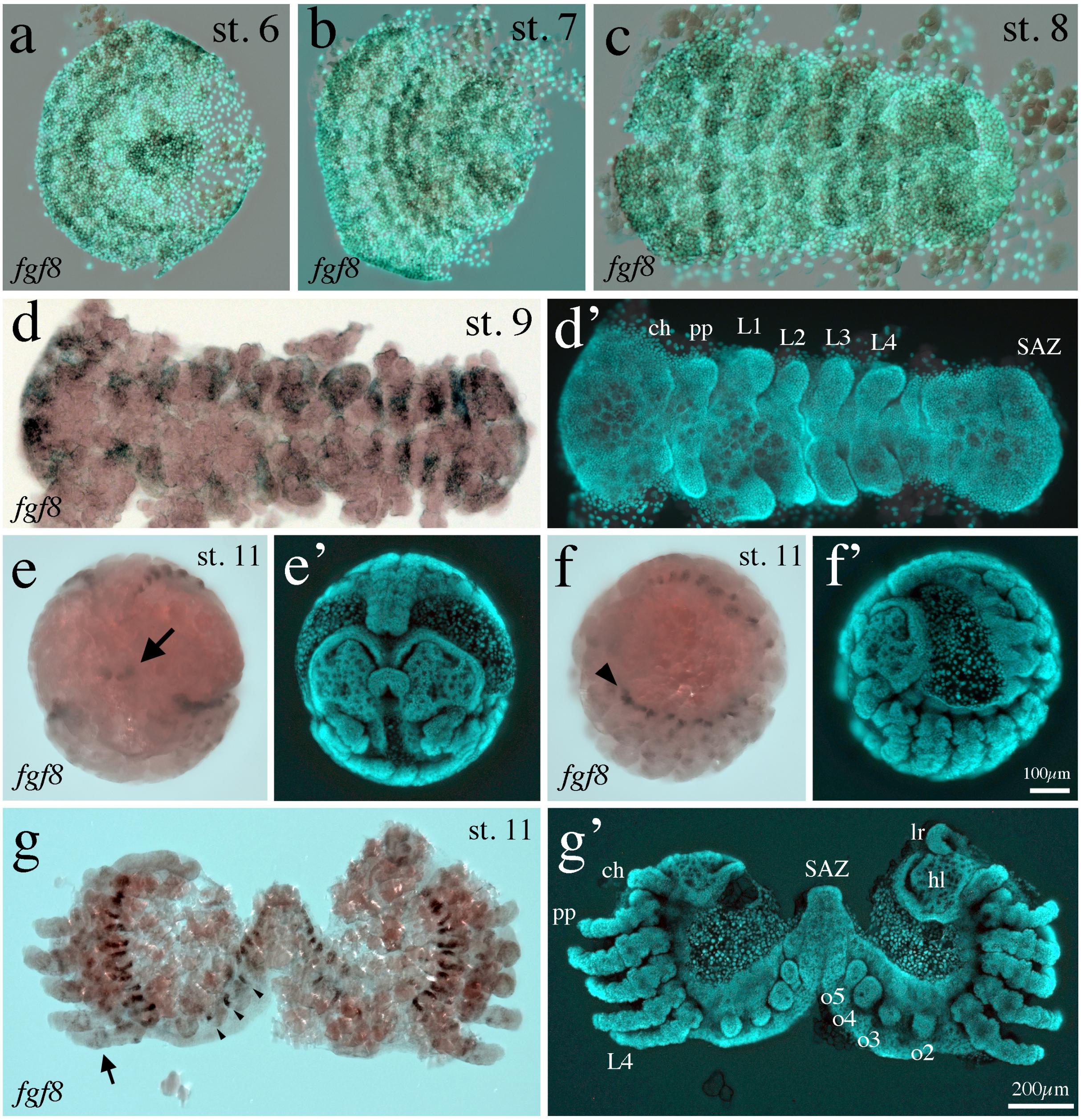


**Figure S10. Expression of *Pt-fgf8* (st. 6 - 11).** (**a-d**) Flat mounted st. 6-8 embryos showing segmental expression of *Pt-fgf8* (bright field/Sytox green overlay). At stage 8 *Pt-fgf8* is strongly expressed in the developing appendages, the head and the segments of the opisthosoma. At stage 11 *Pt-fgf8* is expressed in the region of the stomodeum (arrow in **e**), along the dorsal ridge (arrow head in **f**), in the buds of the opisthosoma (arrow heads in **g**) and in faint rings of the prosomal appendages (arrow in **g**). Whole mounted (**e** and **f**) embryos in an anteroposterior (**e**) and a lateral (**f**) view. The flat mounted embryo was halved along the ventral midline. Abbreviations: lr – labrum, hl-head lobe, ch – chelicera, pp – pedipalp, L1-L4 – walking legs 1-4, o2-o5 – opisthosomal segments 2-5, SAZ – segment addition zone.

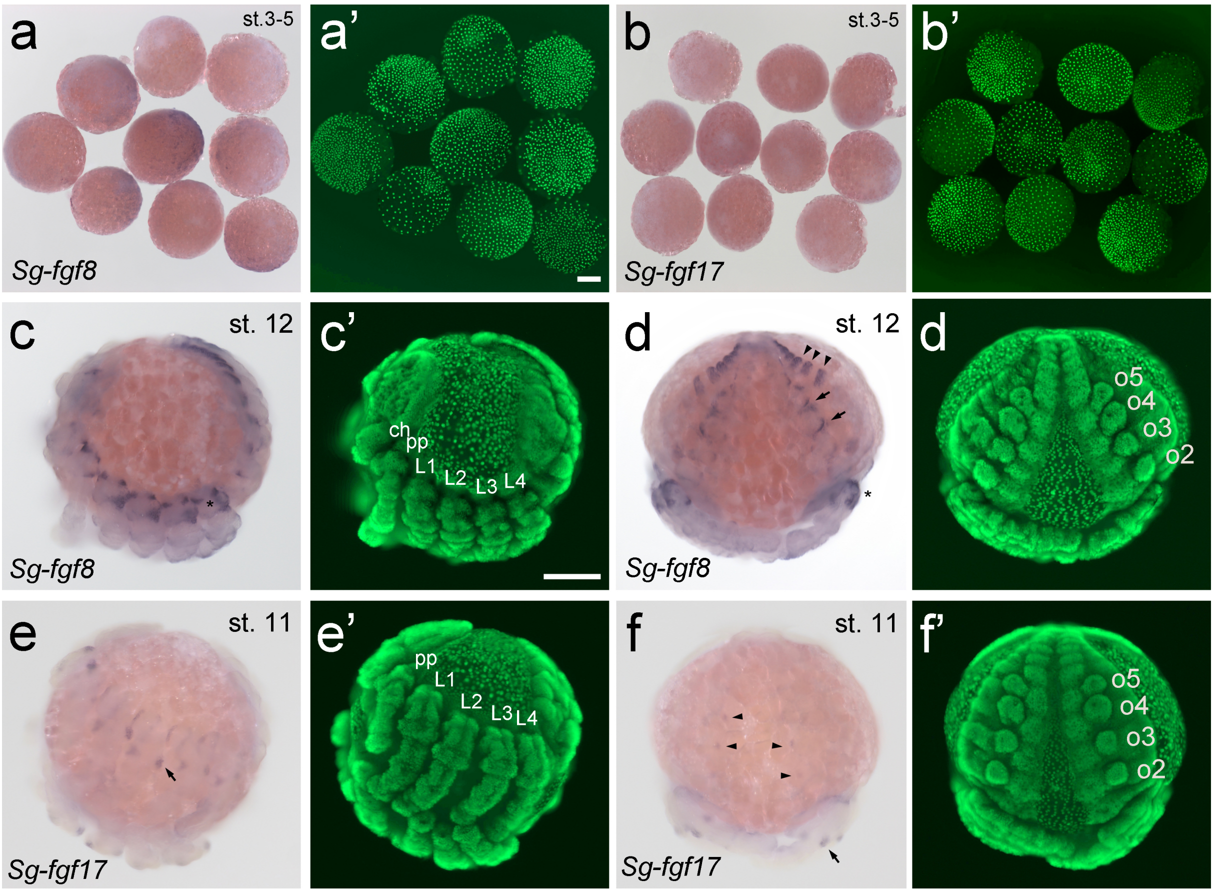


**Figure S11. Expression of *fgf8* and *fgf17* in *Steatoda grossa.*** (**a**) Like *Pt-fgf8* also *Sg-fgf8* shows variable expression at germ-disc stages. Scale bar is 200µm. (**b**) In contrast, transcripts of *Sg-fgf17* are not detectable at germ-disc stages. (**c-d**) At stage 12, *Sg-fgf8* is expressed very similar to *Pt-fgf8* at late embryonic stages (compare to Fig. S10g). *Sg-fgf8* is expressed at the base of the prosomal (asterisk in **c** and **d**) and opisthosomal (arrows in **d**) appendages and in dorsal segmental stripes at the posterior region of the opisthosoma (arrow heads in **d**). (**e-f**) *Sg-fgf17* is expressed in dorsal stripes of each segment and in proximal spots in the appendages (arrows in **e** and **f**). In addition, there are some tiny expression spots in the in the region of the ventral neural tissue (arrow heads in **f**). Abbreviations: ch – chelicera, pp – pedipalp, L1-L4 – walking legs 1-4, o2-o5 – opisthosomal segments 2-5.

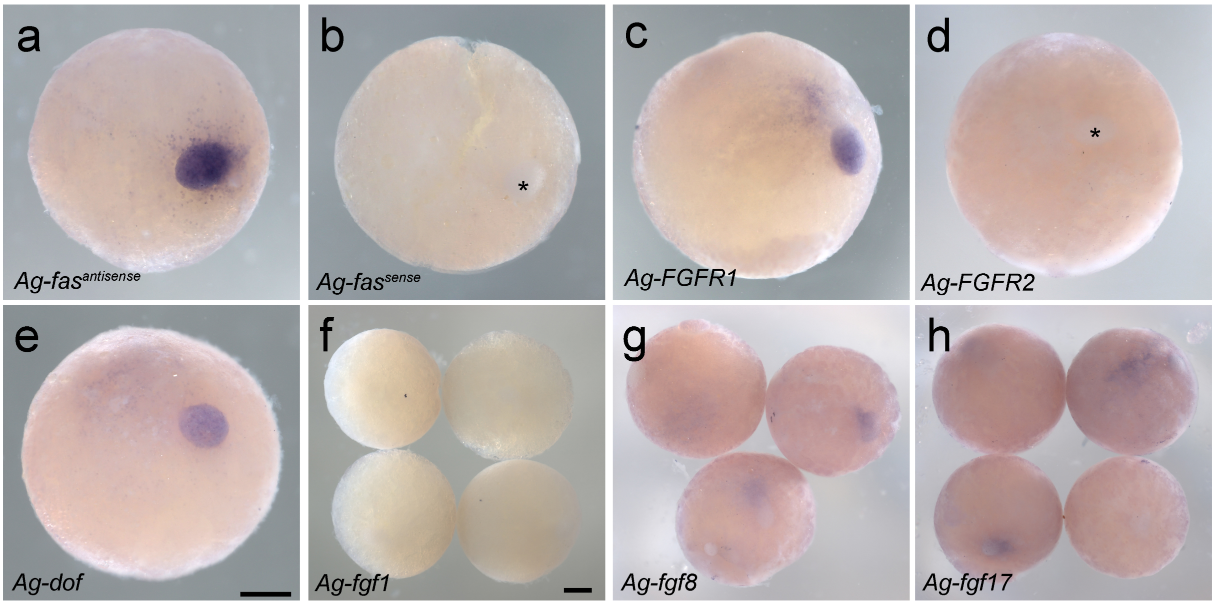


**Figure S12. Expression of Fgf components in *A. geniculata* embryos (st. 5).** (**a**) As in *Parasteatoda*, *Ag-fascin* is a strong marker for the cells of the cumulus (as a negative control (**b**), the sense probe of *Ag-fascin* was used). (**c-e**) Similar to *P. tepidariorum*, *FGFR1* (but not *FGFR2*) and *dof* are expressed in the cumulus cells of *A. geniculata*. (**f**) *Ag-fgf1* shows no expression at stage 5. (**g-h**) variable expression (often enhanced in the region of the cumulus) of *Ag-fgf8* and *Ag-fgf17* cumulus migration stages. The asterisk in **b** and **d** indicates the position of the cumulus. Scale bar is 500 μm.

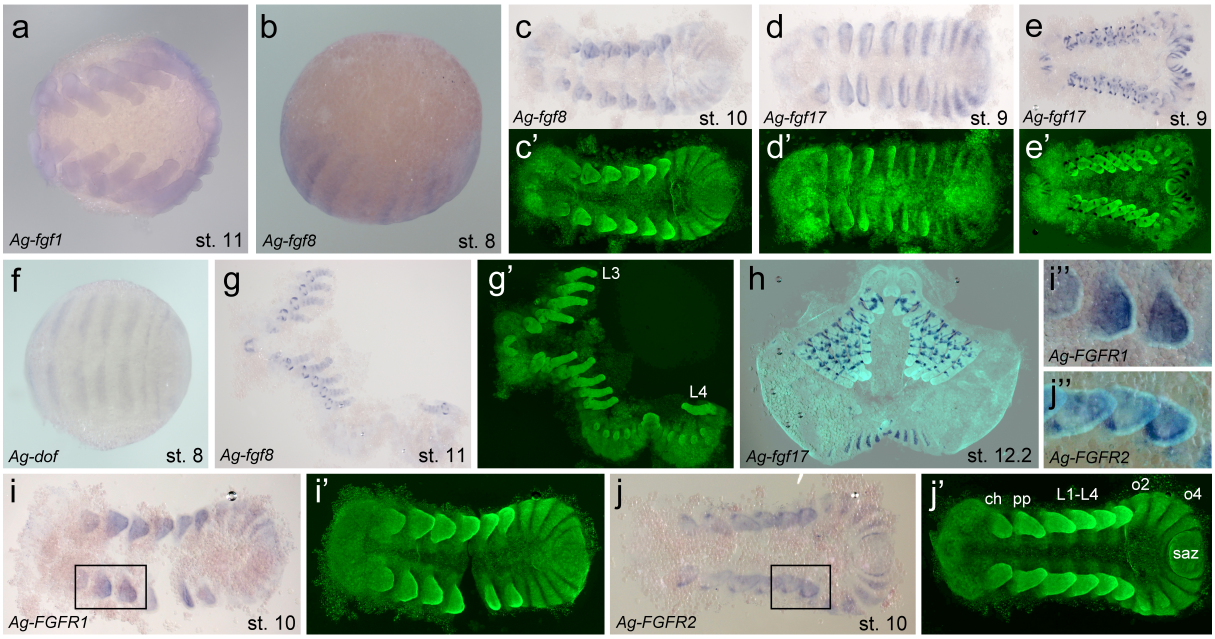


**Figure S13. Expression of Fgf components in *A. geniculata* embryos (st. 8-12).** (**a**) Like *Pt-fgf1* (compare to Fig. S14) also *Ag-fgf1* shows no, or weak ubiquitous staining throughout embryogenesis (e.g. stage 11). (**b**) *Ag-fgf8* is segmentally expressed at stage 8. (**c**, **g**) At later developmental stages, *Ag-fgf8* is expressed weakly in the segments of the opisthosoma. In addition, several rings of *Ag-fgf8* are detectable in the appendages. *Ag-fgf8* expression is also detectable in the labrum of late embryonic stages. (**d**, **e**, **h**) *Ag-fgf17* shows very strong segmental and ring like expression in the appendages of the prosoma. (**i**, **j**) At stage 10, *Ag-FGFR1* and *Ag-FGFR2* are expressed in the opisthosomal segments and in the mesoderm of the prosomal appendages. Within the mesoderm of the appendages, *Ag-FGFR2* seems to be enhanced along the dorsal cells (see **j"**). Boxed region in **i** and **j** is magnified in **i"** and **j"**. Abbreviations: ch – chelicera, pp – pedipalp, L1-L4 – walking legs 1-4, o2, o4 – opisthosomal segments 2 and 4, saz – segment addition zone. Staging according to Pechmann 2020.

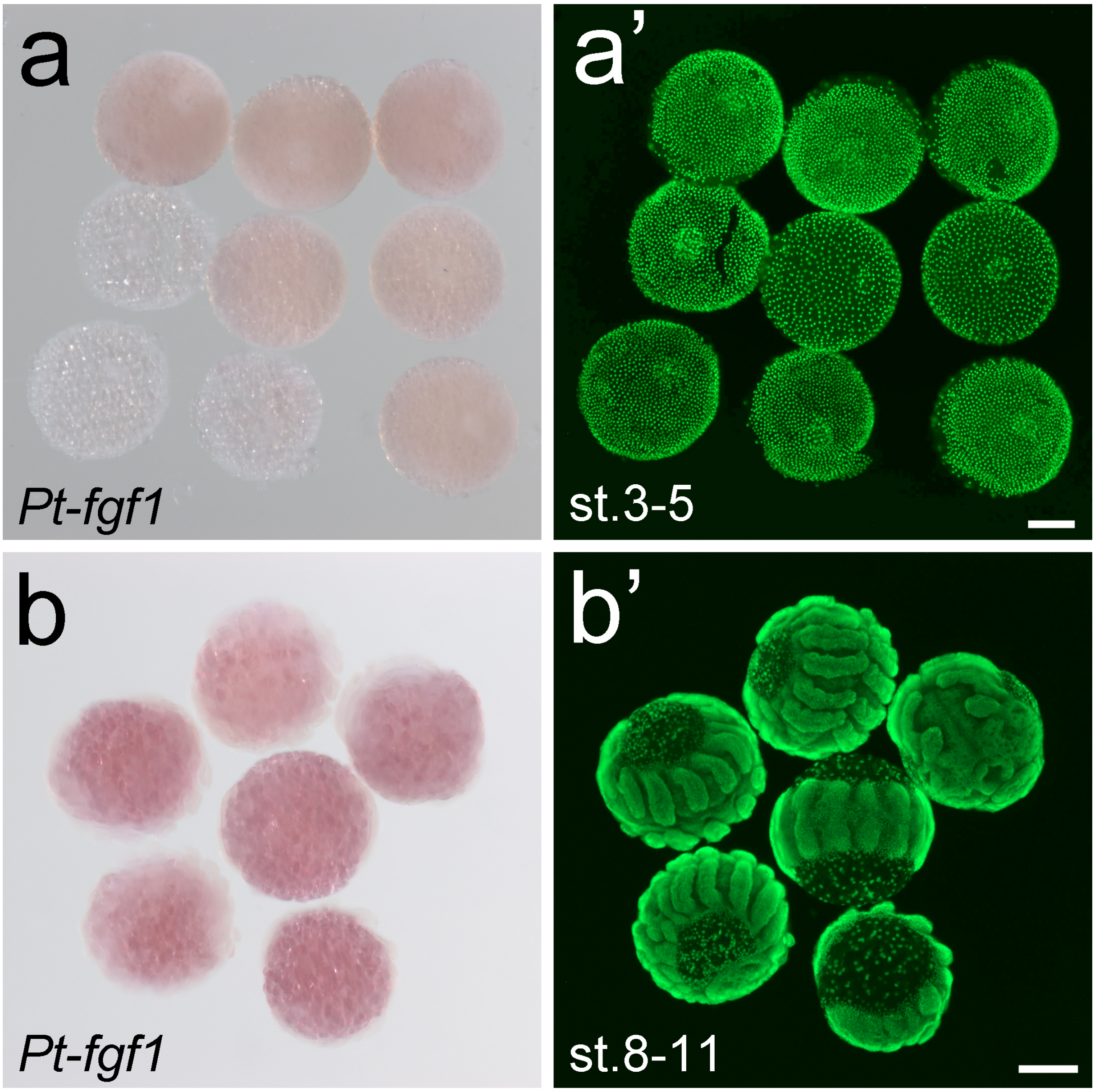


**Figure S14. Expression of *Pt-fgf1*.** *Pt-fgf1* shows no expression at any analysed embryonic stage. Scale bar is 200µm.

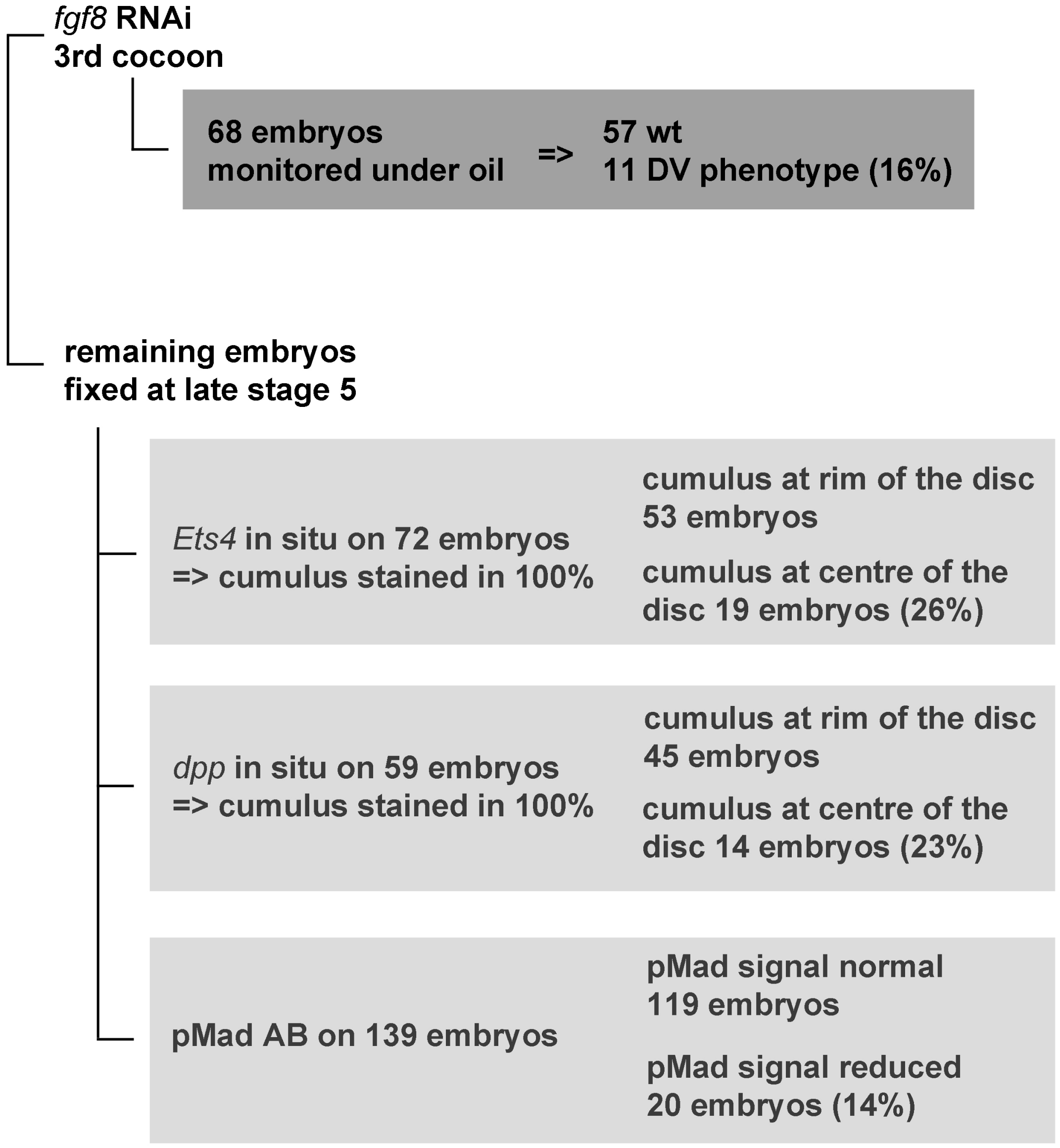


**Figure S15. Strategy to analyse the influence of *Pt-fgf8* on the BMP signalling pathway.** Embryos of a severely affected 3rd cocoon were used for this analysis. The embryos of the cocoon that were used for this diagram were also used for the experiments shown in Fig. 3 g-j. The analysed cocoon contained around 350 eggs. 68 embryos of this *Pt-fgf8* pRNAi cocoon were monitored under oil. 16% of these embryos showed the typical *Pt-fgf8* knockdown phenotype. The remaining embryos were fixed at late stage 5 and these embryos were analysed for the expression of *Pt-Ets4* (72 embryos), *Pt-dpp* (59 embryos), and for the activation of the BMP signalling pathway (139 embryos; via pMad antibody staining).

**Movie S1**
Comparison between a control and a *Pt-fgf8* pRNAi embryo. Movie starts at early stage 5 of development. Cumulus migration was not observable in the *Pt-fgf8* pRNAi embryo and the radially symmetric germ-disc of the *Pt-fgf8* pRNAi embryo was not transformed to an axially symmetric germ-band during further development (compare to the control). Instead, the cells of the *Pt-fgf8* knockdown embryo did overgrow the yolk completely, resulting in a ventralized tube. Compare to Fig. 1.

**Movie S2**
Time lapse imaging of the embryogenesis of *Steatoda grossa*. A regular blastoderm was formed at around 19h. Germ-disc formation was observed between 20h and 40h. Cumulus migration was observable at 40h-55h. Germ-band was formed until 150h. The movie ends at a stage that is comparable to *Parasteatoda* stage 9 (beginning of appendage development).

### Supplemental Sequences

Depicted are fibroblast growth factor sequences from various spiders. Protein sequences were used for the phylogeny shown in Fig. S1.
*Steatoda grossa fgf-8* and *fgf-17* sequences were cloned and sequenced during this study (see material and method section).
*Acanthoscurria geniculata fgf* gene sequences were found in the published transcriptome (Pechmann 2020).
Transcriptomes published by Kono et al. and Sánchez-Herrero et al. (Kono et al. 2019; Sánchez-Herrero et al. 2019), were searched for *Cyclosa octotubercula*, *Argiope aemula*, *Gasteracantha kuhli*, *Zygiella dispar* and *Dysdera silvatica fgf1*, fgf8, and *fgf17* sequences.

**Steatoda grossa**

```
>Sg-Fgf1
GGACAGGGGCATTCCCGGACACGTGGGATCAGAAAAGCAGCTACACAGCGGTACAGGCTACAATCTAGTCATCGATAGTGAAAACGGCGTGTAC
GGAACGAGAGAACCATTCAGTGAAGAAGCTATTCTACAGTTTTACTCCAGAAGAGGAGGAGAAGTTCAAATAAGAGGCAAGAAGTCTCAATTAT
ACTTGGCAATGAATGAAAAAGGAAGAATATATGCAGAGGCTGATCAAAACTCTGAAAAAACCTGGTTTAAAGAAACTTACGAGGGCGGATACAA
TCGTTATATATGGCTTGGAAATGACCAAGAGTGGTTTCTGGGCATCAAGCGGTCTGGAAAAATGAAGAAAGGCTCTAGAACGAATCAT
>Sg-Fgf1
DRGIPGHVGSEKQLHSGTGYNLVIDSENGVYGTREPFSEEAILQFYSRRGGEVQIRGKKSQLYLAMNEKGRIYAEADQNSEKTWFKETYEGGYN
RYIWLGNDQEWFLGIKRSGKMKKGSRTNH


>Sg-Fgf8
CAGGTTACCTACAGATTGCCTCCCTAGCGTACCTGTGTAGTTTTTTCTTCAAAATGGTATCTTGTCAAAGTCTGTCCGTCAGTTTGGAGAAGGA
AGCTTCGGCTCTTTTACAGAATTCTCACGACGTTGTGCGGACGTACAAGCTCTACAACCAGTGCAGTGGAAGGCACGTGCAGATAGTCGACATG
GCAGTCAACGCCAGGGGACCCTCCGACAGTATACACGCCAATATAAGTTTCAAATCAGTTGTTCATGTTCGACACAGTCACGCCATCAATATAA
AAGCCCAAAACTCCGAACACTACCTCTGCTTTAACAAGAAAGGAAAACTTGTAGTCAAATTCAATGGGAAGAAAAAAGCACCACTCATGCGGCG
CAAGGTGTGTCTCTTCAAAGAGGGCTTTAGCGATGACCATTACACAGTCATACAGTCACTCCATGACTCGTCGTGGTATGTAGGCTTCAACAGG
AAAGGAAAGCCTCTCAAGGGAAGTCTCCACTCTGTCAAAAAACACCAGAAGTGTTTCCATTTCATCAAGAGGGAGCACACGTAC
>Sg-Fgf8
GYLQIASLAYLCSFFFKMVSCQSLSVSLEKEASALLQNSHDVVRTYKLYNQCSGRHVQIVDMAVNARGPSDSIHANISFKSVVHVRHSHAINIK
AQNSEHYLCFNKKGKLVVKFNGKKKAPLMRRKVCLFKEGFSDDHYTVIQSLHDSSWYVGFNRKGKPLKGSLHSVKKHQKCFHFIKREHTY


>Sg-Fgf17
GTTCATCTGCACTTCTGTCATGGTATCTTGCCGAATCATTTCTGAGGCATTAATTGAAAGCTATGCTCATATATACTTACTCGACAACAATATT
AATAATTTTACGCCGTACAAAAGGCATTACAAGTTCTATAACCACTGCAGCGAAAAGCAGATTCAAGTTATTGGAAGTTCTATAACAGCTTTGG
GTACACCCCAGAGTCCAAATTCTAATCTAATGCTACAGTCAGCTCATTATGATAATCATTTTGGGGTCAACATCATGGGTGAAAGAAGTGGTCG
CTACATTTGCTTCAACAGGAAAAGCAAGCTGATTACAAGGTTCTCTGGAGCTAGCCCTCGCTGTATATTTACAGAAGAAACTAGTTTGGACAAT
ATGTACTCAGTTTTACGATCAGTTTTTAATCCTGAATGGCGCATTGGTTTCAATAAAAGAGGCAAACCCATTTCTGGATCAAATTACAAAAAAA
TACATCGCAGCCGGTGTTACTATTTTACAAAGCGGGGACATAATTATTTAAACCATCTTTATAAGAATGACCAACCAGGACCAAAGATTGGTAA
TCCTAATAAACTATCTCACCATCTCTGGAATGACACACGG
>Sg-Fgf17
```

FICTSVMVSCRIISEALIESYAHIYLLDNNINNFTPYKRHYKFYNHCSEKQIQVIGSSITALGTPQSPNSNLMLQSAHYDNHFGVNIMGERSGR
YICFNRKSKLITRFSGASPRCIFTEETSLDNMYSVLRSVFNPEWRIGFNKRGKPISGSNYKKIHRSRCYYFTKRGHNYLNHLYKNDQPGPKIGN
PNKLSHHLWNDTR

**Cyclosa_octotubercula**

>comp6270_seq1_(Co-Fgf8)
GTCGGGGTGGTTCACTATTTCTTTTGATGACAAAAAAAAAAAAATTGCACAGAACTGAATTAATATAATAAAATCGCATTCTTTCGTAATTCTGG
AAAGACAGCAATTGAAATGGAATTTTGGCAGAGCTGTTTTATTTGAAGAAATATGCGTGGGGAAGGTAACTGGATTGCCTAACTGAAAGCATTA
ACAATATTTTCTGGTTTCTTCAAATTGCCTCCCTAGCGTACCTGTGTAGTATATTCTTCAAAATGGCATCGTGTCAGAGTTTTTCCGTCGATTT
GGAGAGGGGAGCTCCGTCGATCCTGGAAAATTCCCACGACGTACAAGCTCTCCGCAGGACGTACAAGCTCTATAATCAGTGCAGTGGAAACCAC
ATACAGATCATCGGGAAAACAGTCAACGCCAAGGGTAGTCCCGACAGCCCTAATGCCAATATGAGCTTCACCAATGCCCGTTCAAGGCATCATT
CCAACGCCATTAACATTATGGGACAAAAATCTGGTCGTTACCTATGCTTTAATAAAAAAGGAAAACTAATAGTCAGGTTCAATGGAAGGAAGAA
GCTGTGTCTTTTTCGGGAAGGTATCAGCGAAGACCACTACACAGTGCTGCAGTCATTATACGATCCCGAATGGTATATTGGCTTCAATAGGCGA
GGTAAACCTCTGAAGGGAAGTCTGTATTCCAAAGCAAAAATGCAGCGTTGTTTTCACTTCGTCAAAAGAGATTATTCCTACGGCATGGATGAAT
ACACACCGCCTGGGCCTAAGATCAAAATCTCTTGGAAACTAAAAGACTTATTGACTGGCAATGGCCAACATCGATCGCGACCGCAAAGAAAAGT
GCCAAAAACGCCAACGAGGTGAAACTCCTGTGGAATTCCCGTGCTTCAGATATTAGATTCGAATTTTTTTTGGTTGTTGCTTCTTTTTCCTCTA
ACTCAAAAAAAAATTTCGAAATAGATGAAGATGCCAAGGCAGGCCTTGCGCGTTTTCGCTTCGATGTTTGCCAGTGTGTGGCTGATCACGCTTC
CAATCTGTTTTGAGGACATTAGCTGGGGATCGGCTTTGTGTTATTGCAAATGAAGTGCCAATTTTGTGTTCTTAGGAACCCCCTAAGAGTATTC
AACCAATATCTCTTCGGATTTTCCTCGATGTTCAAATGCAACTAGGTATTCTCAAAAGTTCTTATCTTAGAATAGGACTTTCATGTTATACGTA
GTATTTTCCTGAACTAAATGTTATTTATCCTTAAGAGTAGAAATTTATTTATTGCAATGTTAGTACCTTTTAGGAATTGTTACAATTAACTGTG
TTCAGAAAGTTGCTCCGAGTGAAACTAGATTCTAAATATGATTTTTCTTTTCTGGTAACCCGATGACTAGAAATATGGTCCATCCTTAACTTGT
CGCCTATACCATCTCGCACTAGAATAAAATATACAAGAAATTATCTCGTCAATGCATGCTGAAGTTTTCTCCAACCCTGCAAAGTGCTAGTGAT
CCAAAAAAAAGGATTTAAAGCCGATATTCTCCTGAAAGGATGATATCTTAAAAGAAAAATCAGACCGTACTCAATCCAAACAAAAAAAATCGGT
GTGAAATGGTTGTAGTCTGACTGTAGACAACCTGAGTCATGTATGTGAAAACATCGTCTCCCGCTTCTGTACAGAACATTTTTCGTTTTGTGTG
CTGTTCGATTTATGTGCCATCGGTCAAGATTCTTTCCTGATGGCATTTTGTTTTCATATTTTTAGATGACTTATGTTTGTTTTGAATCATTCCA
AACTGTGGCCAGTCCAGAATTTTGTAAAAATAAAATAATAAGTGCGCTTGACACCAAGCCTGAATTCGCAAGCAAATGTTACATTTCCAAGTTG
CATTGTTAATTGTTTTGTGTATAAATCTGTATATAGTTATTCTAAGTTAATGAACTTGATGTTTCATGATTTTAATAGAAGTAACAATAGTAGT
CTTTTGCTTACCAGTAGTGCAGATCTTCTGCCAGACATTTATTTCTTTCCTAATAGTATATAGAGGCATGTGTGGCAGAATTAAGTTGTTTTG
CTCCTGATCCCGAGTTGAATTACTTTCTTCCTTTTGCTTTCCTCGTTCCGAAAGAGGAATTTGGAGGCTGGAAATAACTTAAATATGAGGTGAA
ACGTGGGGAGACAGAAAAAAAAACAACTTAACTATGAAGGGAAATGTGGTGAAGCAGCTGGGAACAACTCAACTCTCTTGGTTAGAGACTTATGT
CTCATTCTAAGTTTCAGGGATTTAGGATATACAAGCAAAGAAAAATAATCCTCTTGTCAGAGGACCTGTACCATCCGTAAGCAAAAGCTTAACT
TATCTTTTTAGAACATTTTTGTATAGCTTAATCCGTTAGGAAGTTTTAACATTGATTGGATACTTATGACAGTTTAATAATTTATTTAACATTA
TGTATATTTAACATCATTGTTTTGAGTATTTGAAAGTCAACAAAGGACTTAGGAATGTTCTACTGCACAAAGTTTTAAAAAGAATTTTGAAGGG
GGAAAAAAAGCTGATATCATAGTAATTAGTCAAAATTAGATTACTTGATAATTCATGTCTACTTAATTTTTCTTGATTCATTTGAAATGACTAT
AATTTTATGATAATCAAACTATGTTTTAATACTTATCGATTCTCACTGACAACTTTTTAAAAATATCTTATTTCATAGAGTAAGAACAAAATCT
GAAAGTAAAAACAAGGATTTAAAAAACTCAACTGGATTGCTTAGCAATTGTTTTTAAAATTATTTATATTTATTAAGATTGATTTAAAAAAAAG
CTTTAATTAAGAGAGAGCTGTTTTTATTTCTTGCTGATTGAAAAGTTCTTACTCTTCGGATTAAGATATAATTTTTAACCATTTTGTTAATTTG
TTTCCTTAAAATTATTTTTATTAGTTTCTACTTTAACAAGTAATTTTTTTGCTATATATTTTATTTAAAATAAAATATCTGTATCTTGGTTGTT
TTCTCCTTTGATTTATTTCTAAAACGGAAATAAATGAAAGGAGGAATAATCCAAATCAAAAATTTTGTCTTTTGGCTAGGAAGAACGTGTAAAC
TTTACTTTCTTTATTGTTGATGTAAAAAGAGAGATATCTTAAAGTTTTGCATTTCTTCCATACATGAAATATTAAAACTATTTAAATCCTAAAA
TGCTATGAAGAGCACAGAAATCTCAACTATTATGTTTATTGTTTTTGTAATTGGATGATTTTTCAAAATCTCCAAATTGTTTTAGTAATTGTGC
AATTCTTTGCATTTTATACACTGCCATTTTGTATTTTAGCATCTTTCATTCCTGTTCATTCAAAAGACCTCTTTTCATTCTAAGTTGTAAATGT
TGTAAAAAAATAAATAGAAAAGTCTTTGAAACAGAAAAAAA
>Co-Fgf8
MASCQSFSVDLERGAPSILENSHDVQALRRTYKLYNQCSGNHIQIIGKTVNAKGSPDSPNANMSFTNARSRHHSNAINIMGQKSGRYLCFNKKG
KLIVRFNGRKKLCLFREGISEDHYTVLQSLYDPEWYIGFNRRGKPLKGSLYSKAKMQRCFHFVKRDYSYGMDEYTPPGPKIKISWKLKDLLTGN
GQHRSRPQRKVPKTPTR-

>comp2940_seq0_(Co-Fgf17)
AGACGAGTACGTGCCTATTGCTGTTTGAAAGCGCGAAAGAATTACGTGTGTTAGTAAGCTGATCATTTTTCCACTGCATTAAATCAAAAGGTTT
TTTTTTCAAATGTATTTCGATAATGAAACGGAAAATTTTAAACCGTGAGCCTGAATTAATTAATTTTGGTAAGTTAACGTCGTTTCGGAGCTAC
CCGTCAGAGACCTATTAATTACATCATGGAATAGCTCTTTCTTCGAATTAAATGGTAAAAGTAGTTAATAAAAGTCGTTTACAATTAAATGCTT
GTTCAGGCTTTAACAGGTTTTTCGGATTATTTCGAGATGAATCAGAGTTATATGCTTTGATATAATATATAAATTGCAGTCTCGCATTTGTTAG
GAAACAAAACGGTTGATTTTTGTGAGCATTTTCAGGTTGGTTACTGGCGTAAAATGTACAACGTTGCTGCGTTTAGTTAGAATAAGAAGGCGAC
ATTACGGCAAATGAAAATGAAACTGGAAAAAAGATTTCGTTCCGGCATATTACATAAGTCTCCTATATACCATTTACTACTGCTTTTGAGTGCC
TCTATGTTGATGGCTACCGTCTGCCTTGCTTTCTCTCCGGATGCCCTTATAGAGAGTGGTTCCAATTTCCTTTCTGACGATAATACCATAGCTC
ACTACAGCATTACGAGACGTTATAAGTTTTATAATCATTGCTCCTCTAAGCAAATTTTAATAGTTGGAAGTTCAGTGACTGCATCTGGTAGTCA
AGATAATCCTAACACAATTTTAATCCTGCAGTCGTCACACTACAAAAATCACAAGGGGATTCACATAATTGGTTCAAACTCTGGTCGATATCTG
TGCTTTAGCAGAAAGTCAAAACTAATCACAAAGTTCTCTGGAGCTAATCCTCGATGCATTTTCGAAGAGTTACTCAGTCCTGACTTCTATACAG
TCTTGAGATCGGTCTTCAACAAGGACTGGCTTGTGGGCTTTAACAGGAGAGGCAAGCCCCTATTGGGATCCAACCAAACCAAAGCACACAGGAG
TCGGTGTTATCACTTCACAAAGAGGGACCACTCGTATCTTGATGCCCTGCACAAAACCAGGCATTACGCCCCAAAGATCTCCAATCCCCACAAT
TTGATACATCTTCTCCATGAAAAGCGAACAAGACGACAGAAGCGACACGCCTTAAGATGAACAGTGTGTGCAGAGGGAACTGAGTGGCCCCCAC
TTAACAAAAAGGTTGTGACCTGTTTTTTTAATTTCAGTACTGGCTTGGAGTAAAAAAACGAACGACAGTTACGATTGTGCCACCAGTTTCCTA
ACCCACTTCCTCCAAAACTATAAATGTCGTTCAGTTTTCAAATGCCAGTCATTATGTGCTTCAAGTGTAACCCCGGTTTCATGATTTCATTTGT
TGCTTTTGTCATTTCTGGAACTGTCAATAGAATTGAAGTAGACTGTAACTGCCCATGATTTATGCTTTTTAAAGCCCTCTGTGTTTTCTGAAAC
ACTACGTTTTTAACTCTTATGTTTGTCTAAGTATCAATGGGCTAAAATCATGCTTTTCCTGTTTTTGTTAATATTCGTTTTATATTAGCTGAGG
TCATGACAGGTAGACTCAAAACTAGGTTCCTATTGTTACAAATTTCTCGAAGAATATTTTGCTAGTTTTATATACATTTTTGATAACATTTTAA
TAATAAATTCTTTAAATTAAACATTAATTCTCCGAAAATAATTGACAGTTCTATTATGAGAAATTCCTGATTTTAGACGATGAAACATTCCAAT
TTTAGTTTTGCTAGAGAACTTTAAAATGGATTATGATTTTAATGTATAATTGGTTCATCAGGGTAAACTGAATACTTTTACCGTATTTATGTAG
AATTGTTTTATTAATTGTGGTATGAATTGTTGTATGTACATATTTTTATTAGCTGTCTTGGCAATACTCAATGTTCCATTTTTAAATGAGCTAG
GTTTTAATTGTTACTGCAACATCCTAATGTGCTGTTTATTTTGATCAACGAAGAGCTTGAATCGAGACTTAAGTAAATTTCATTGAACTGTCAA
ATGAAAATCCCTATTTGTGGAACAAGAATCATTGAATTGCTTTTAAGTAAATGTGGTTTTATTAAGTTATGGTGCGCTGTTATGTATAACTTGC
GACATTCCATCCAGATTATATGGGTTCGTACTGCTACATTCCAGCATTTTTAAATTTGGTTTTTAGTTTTTATCTTATTTTAACTTTTTATTGA
AAATTAAGTTATTTAATTCTGACTCCTCAAACTACTTTTATTTATAAACTGAATTTTTTCTATGATATAGAACTTAGAGAAGAATATAGGCTAA
AGAATTTTTAAAAAACGAACTAAAGTATTTAGTAAATATGCTAGAAGATTTTTAAAAAACATTTCCAATGTACTTTGAAATTTTACATGGTTTT
GCGTTGAAGTTTACAATTCTTTAAGTTTCATGTTTACAGTTAAGTTTTAGTTTATTCACAGAATATAACAACTTAAAACTACAATCTTGTTTTG
GATATGTTTTGCGCAACAAAATTAAAGCTTACAAGGCAAGTGCTTAATTTGCCTGACCACTAAGTTAAAAAAAAAATCATTTTTTTTTATTTCA

CAGCACGACACTTGCATCCTTACATTTACAAGAATTTCATTTTTGAATTTTATAGTGTTTGACGGAAGGTTTAAACTTTTAAAATGCATTTTTA
CATTGAACTCTCTCATATAGAGTACGTGCCGCAAATATTGCATAGATTCATTTTAATCATAAATCGCGCCATATTTGGAAATAAGAAATTTAGA
TGAATACAGGGAAGATTTTCGAGTATTGAAGGGAAGCTTTCAGAGTATTCTGAGTTTGAATAAAATCACTTCATCTGACTTAGTTTGAAATACA
GACTTCAAACGCTGGGTAATCGAATATGTCTTTTTTTAAAGAAAATTAGCCATCACTTTTGAATGGGATGTATTTATTCATAGTTCACTGAGGA
AAAAATGCATTTATATACGAGAACCCAGCAAACATATATAAATTCTTAATACATTTGTTAACTAAAATTTATTCCTTCTTTTAAGAAAGTTTTG
CTATCTAATATTTTTATGCTTCGTTCTGAATTTAATCGTTTCTAGGATTCGTATTTTTTTTAATAAATGAGATTGTAGAAAATGCAATAATTTC
GCAACATGCCTGTTGTAAAATAATTTCTGTACAGTTTTTCAGATATATTATTTCAAACTTCAAATAGGCGAAATAGTCTCTGATTATAAATGTG
CCAAAATTGATTCTATAAATATTTTTTCAATAAAGTCTATGTGTGCCAAAGTCACTCAAATCTACCTCTTTCATAATACAATGCAATTTTAAGC
TCTTGCTATAAAACAAAGTATATTCTTGGAAAAAGGTGCTACTGTTAAAAGTGAGAGATAATTGGTTTGTTTGAATGCAAAAAGTGTTACAACA
AATCAAGAACATAACCTGTTATGAATCTGCCCCATCTTGTCAAAAAAGTATCTGTATCGCATTAAGACATAATAAAGCTTTTTATGGGCATTGA
AAAAA
>Co-Fgf17
MKMKLEKRFRSGILHKSPIYHLLLLLSASMLMATVCLAFSPDALIESGSNFLSDDNTIAHYSITRRYKFYNHCSSKQILIVGSSVTASGSQDNP
NTILILQSSHYKNHKGIHIIGSNSGRYLCFSRKSKLITKFSGANPRCIFEELLSPDFYTVLRSVFNKDWLVGFNRRGKPLLGSNQTKAHRSRCY
HFTKRDHSYLDALHKTRHYAPKISNPHNLIHLLHEKRTRRQKRHALR-

**Argiope aemula**

>comp11974_seq0_(Aa-Fgf1)
CTAAAATACTCATCACTAAAAGCCTCATGTACAAAGTGAACGAAATGGGATCCATGCGGGCCAAAACTAACGAATGAGCTTTGGTTTAAAACGT
AACTTTCTAACAAAGAAACTCACATCGATACTGTACTGCATTTCTTTTTTTTAAAAAAATACAGAAAGCAGTTGAAATAAATAATGATAAAGAAA
ACCTGACTGATTATTTCTTCTATATAATTTTCCATCAGTTACTATTGCTATTAATAATGTTTCATAACAAAATAATTATCAAGCTATTTTCTGG
ATTAAAGTTGCATTTACATTAGGACATTTATTGGGCGCAATAGGAAGAACTTTAACTCCAATCGAAATCGAGAGACTAAACGAAAATCTGACAT
CAAGAATGGCTCGAGTTAGAATCGATTGTCCAAAACATCAAGTGGTCCTTATGGGTTCATGTGGCATTCTGTTGTGCCTAACATTTGTCAATGT
AAATTCCCTTTGAAATCTAGAATCAATGGCTTCACCATAACATTCCACATCGCCTTATTGCAAATACAACGCCCAAAATTAAGATTGTGGTAAA
TGCAGTTGTATTGGCAATGGGGCGATTAGTTAGGATTCGGCATACCAAGCAAAACTGACTTTCCTTTTCTATAAATATAAAAGTAAAATAATTT
GCGAATTTAAATGTCATTGATAGAGAGTCCGCGATTCCAAAGAATCTAAAACAGAGGACAAGCACCTCCGCTTATTACTTGATCTCCACTATTT
GCAGATGCTTCAATCTAACAAAACATCAAAACTGTTAAAGTATACACCAATGAAAGAGTTCTTGCAGCCGCTTAGTTATACGAAGGCTCTGCAG
ACCTCGGGACAAACTTCACAGCGTTTTGGTCTCGCCTTGTTCTGTGGCCACGCTTCATTTTTCCGGATTTTTTGATCCCCAAGAACCAATCCTT
TTGGTTGCCAAGCCAAGTGTAATAATTCCACCGATCCTCGTACTCCTCTTTGAACCAAGTATTTTCAGATTCAGGATCACTCTCAGCATAAACT
TTTCCTTTGTCGTTCATTGCTAGGTACATGTTAGTATTTTTCCCTTTGATTTGAACTTCTCCTGGTGCTCTTGAGTGAAAATGTAGAATAGCTT
CTTGGTTGTAAGGCTCCCTTGTGCCGTAGACATTCCCTTCGCTGTCAATGACTATGTTGTAGCCGGTGGCACTGTGCAATTGTTTCTCCTTGCC
AATGTGGCCCGGTATTCCGCGGTCCGTCTCCATGGAGAGGCCAGCATAGCGGGGACCCGTCTGCTCTTCTTCTGTGGCCATCTTGTTCGGAGGC
CCGAGAGCAATGGGATCCTGCTCTTTCGCACAGGTGGAATACCGAGCAGTTGTTTCATCAGCACTATTCCAACACCAAATGCACATCGCCGGTT
CGTCACGCATTATAGGTATCCACTGAATAAAAAAAGAAATTTCGTCCGTCACTGCGACTCGCTCAAGTTACACTGCCACTGAGCTACTAGGTGT
CATGAATTACTATTCAATTTTTTTCCGCCGCTCTTTTTTTTATATGTGTAGGAG
>Aa-Fgf1
MRDEPAMCIWCWNSADETTARYSTCAKEQDPIALGPPNKMATEEEQTGPRYAGLSMETDRGIPGHIGKEKQLHSATGYNIVIDSEGNVYGTREP
YNQEAILHFHSRAPGEVQIKGKNTNMYLAMNDKGKVYAESDPESENTWFKEEYEDRWNYYTWLGNQKDWFLGIKKSGKMKRGHRTRRDQNAVKF
VPRSAEPSYN-

>comp12173_seq0_(Aa-Fgf8)
GCAGGGCCTGGCTGGACAGCATTTAAGTGCTAGAAAACTGCAGCATGTATTGACGTGAGAATTTCTCGTATATTTTTTTTTTGTCTAGTGCGAC
ACATATTGAAGATAAGAAGAGGGTGTTTCGTAATTTTAAGGAAGGTTTCCAATTTCAGTCATCGATAAACAGAAAAGAAAAAACCACTTTATGA
ATCTTGTTTCACTCGGAACAACTTTCTGAACACAGTTAATTGTAACAATTCCTAAAAGGTACTAACATTGCAATAAATAAATTTCTACTCTGAG
GTTAAATAACATTTACTTCAGGAAAACACTACGTATAACATTAAAGTCCTATTCTAAGGTAAGAACTTCTGAGAATACCTAGTTGCATTTGAAC
ATCGAGGAAAATCCGAAGAGATATTAGTTTAATACTCTTAGGGGGTTCCTAAGAACACAAAATTGGCACTTCATTTGCAATAACACAAAGCCGA
TCCCCAGCTAATGTCCTCAAAACAGATTTTAAGCATGATCAGGCCACATACTGGCGAACATCGAAGCGTCAACGAACAGGAAAACGCACAAGGC
CTGCCCTGGCATCTTCATCTATTTCGAAAAATTTTAGTTAGAGGAAAAAGAAGCAACAACCAAAAAAATTCGAAATCTAATATCTGAAGCACGA
CATTTATGTCGGGAATTCCACAGAAGTTTCACCTCGTTGGCGTTTTTGGCACTTTTCTTTGAGGCCGAGATCGATGTTGGCCATTTCCAGTCAA
CAAGTCTTTTAGTTTCCAAGGGTTTTTGATCTTGGGGCCAGGTGGTGTGTATTCATCCATGCCATAGGAATAATCTCTTTTGACGAAGTGAAAA
CAACGCTGCATTTTCGCTTTGGAATACAGACTTCCTTTGAGAGGTTTACCTCTCCTATTAAAGCCAATAAACCATGTTGAATCGTATAATGACT
GTAACACTGTGTAGTGATCTTCGCCAAAGGTTTCTCGAAAAAGGCATAGCTTCTTCCTTCCATTGAACCTGACTATTAGTTTTCCTTTCTTATT
AAAGCATAGATAACGACCAGATTTTTGTCCCATAATATTAATGGCATTGAAATGATGCCTAGAACGGGCAGAGATGAAGCTCATATTGGCATTT
GGGCTGTCGGGACTACCCTTGGCGTTGACTGTTTTACCGATGATCTGTATGTGGTTTCCACTGCACTGATTGTAGAGCTTGTACGTCCTGCGGA
CGTCGTGGGAATTTTTCAGAATCGACGGAGCTCCCCCCTCCAAATCGACGGAAAAACTCTGACACGATGCCATTTTGAAGAATATACTACACAG
GTACGCTAGGGAGGCAATTTGAAGAAACCTGAATATTAACATGTGGCATCAACAAGTAACCAAGGAGACATGTGTCTCCTAATGGATCCGCCGA
CACGCATTGAAATTTTAAAGAGTCACTTCTCTTGAACAATCCAGCTCGCACCAAAAATGTATAGAAGACATCTAACAAAATCCACACACAAACA
CTTTTTTTTCAAAATGAAACTTTTCTCTTTCAAAAAAAAAAAATTATCCTTTTTAGTTATATTTATCCGGAAGAGAAGACTGAAAAAAAAATCTA
CATTATGCACAAGAATTTTCTTTTTTATGGATATTTCCAGCGCCGTCTGTCACACAAACACCAGCGCAGTTCGTACGCTCGCCACCGCTAGCGA
GAGGAATCAAGATGTGAAG
>Aa-Fgf8
MLIFRFLQIASLAYLCSIFFKMASCQSFSVDLEGGAPSILKNSHDVRRTYKLYNQCSGNHIQIIGKTVNAKGSPDSPNANMSFISARSRHHFNA
INIMGQKSGRYLCFNKKGKLIVRFNGRKKLCLFRETFGEDHYTVLQSLYDSTWFIGFNRRGKPLKGSLYSKAKMQRCFHFVKRDYSYGMDEYTP
PGPKIKNPWKLKDLLTGNGQHRSRPQRKVPKTPTR-

>comp5191_seq0_(Aa-Fgf17)
GGGGCGCTGGCTCATATTCCCTTCTTAATTTTGCTATGAAAAAATTCCACTAATAAATAAAATTAGTTTGGTACACTTTCAGGAAAGTTCAATG
TCCAATTTTAATTGAAAAAATTTAAAGTTCTATAACAGTTTTAAAAAATAAATTAAATTAAAAACCACATTCATTTCAGTTTAAAATAGCATAA
GAACCCTTGTTGGAATGTAAAAGTATAAGTCAATATCCTACGGATGGAATGTTAAGAGTTAAACGTAACAGGGCAGCATCAATTAATAAAATCA
AAATTTATTAGAAGCAATGGAATGATTCTTGCAGCACAGATAAAAGTTTCACGAGTTGACAACAAATTAGAATGTGATTAAGGCCCGATTCAAG
CTCTTTGTTGTTAAATATAATAGCTCCTCTAATATTTCAGAAACTATTACAATTTAATTCGTTAAAAAATGGAACAAACAGCATTGCGAAAATG
ACATAAAATATGTACATACAGAAATTCATACCACGGTTGATAAAACAATTCTACATAAATATGCTAAAAATATTCATTTTACCCTGATAAACCA
ATCATACATTAAAATAATAATCCATTGCAAATTTCTTTTGTAACCTTAAATCGGAATGTTTCATAGTCTAAAAACTGGCCTTTTGTACAAACAA
ATAGTCAAAAAAAATTTTTTTATGAAAAGTTATGCTTGATTTAACGAATTCATCATTAGAATCGTATCATCATAAAAAATGCTTATAAAAATAG
CAAGACAATACGATACTCCTCGAGAAATTTTTAACAATAGGAACCTAGTTTTGTGTCTTGACTTGTCATGCTCTCAGCTAATATAAAACGAATT
TTAACAGAAACAGGAAAAGCATGATCTAAGCCCAAGACATGCGGACATACACTGAGTTAAAAACAATGTTTTAAAAAACACTGAGCTCAAATAG

CATAAATCATTGATTGTTATAGTCTATTCTAGTGACAATTCCAATAATGAAAAAAGCAACAAATGAAATCATGAAACCGGGGTTATACTTAAAG
TGCACAATGACTGGCATTTGAAAACTTGACGTTTATAGTTTTGAAGAAAATGGGAAGAATAGGAACCGGTGGCATAATCGCAACTGTCGTTCGT
TTTTTGCTCCAAGCCAGTGACGAAATAGTAAACAGGTCATAACCTTTTTAATAGTAGGGAGCCACACAATTCCTTCTGTACATAAGTTCATCTT
CTCATTTCATTAAAGGCGTGCCGCTTTTGCCGTCTTGTTCTCTTTTCGTGAAGAAGATGCAGCAACTTGTGGGGATTGGAGATCTTTGGAGCGA
AATGCCGGGTTTTGTGCAAGGCATCAAGATACGAGTGATCCCTTTTCGTAAAATGGTAACAACGACTTCTGTGAGATTTGGTGTCAGATCCTAG
GATGGGCTTGCCTCTCCTGTTGAAACCCACGAGCCAGTCTTTATTGGAAACTGATCTCAAGACTGTATAATAGTCAGGACTGAATAATTCTTCG
AAGATGCAGCGAGGATTAGCTCCAGAAAACTTTGTGATTAGTTTACTTTTTTTACTAAAACACAGATATCGACCACTTGTTGCACCAATAATTT
GAATTCCGTGATGATTTTTGTAGTGCGCCGATTGTAGAATTAAATTTGTGTTAGGGCTATCTTGACTACCTGATGCGCTAACTGAATTTCCGAC
TATCAGAATTTGTTTAGAGCTGCAATGATTATAAAATTTATATTGTCTTCTAATGCTATATTGAGTTATGGTATTGTCAGACAGGAATTTGGAA
CCACTTTCTAAAAGGGCATCCGGAGAGAGAGCGAGACAGAAAGTAGCCATCAACATAGAGACACTCAAAAGCACTAGTAAACGGTATATAGGAG
ACTTATATAACGAGCCAAAATGATGCCTTTGCCCCTGTTTCATTTTCATTTGCCGGAATGTCACCTTCTTATTCTAACAAAATGCTGAAACGTG
CATATACGCCCGTAACTGACTTGATAATGCTTACTAAAATTCACTGTTACGTTTTCCTAACAAATGCG
>Aa-Fgf17
MKMKQGQRHHFGSLYKSPIYRLLVLLSVSMLMATFCLALSPDALLESGSKFLSDNTITQYSIRRQYKFYNHCSSKQILIVGNSVSASGSQDSPN
TNLILQSAHYKNHHGIQIIGATSGRYLCFSKKSKLITKFSGANPRCIFEELFSPDYYTVLRSVSNKDWLVGFNRRGKPILGSDTKSHRSRCYHF
TKRDHSYLDALHKTRHFAPKISNPHKLLHLLHEKRTRRQKRHAFNEMRR-

**Gasteracantha kuhli**

>comp38018_(Gk-Fgf1)
CTGCATTCTGATCCCGTCTTGTTCTAAGACCACGCTTCATTTTCCCTGATTTTTTTATTCCCAAGTCCCATTCCTTTTGATTGCCAAGCCAAGA
GTAATAATTCCATCCATCTTCATATTCTTCTTTGAACCAAGTATTTTCTGATTCAGGATCGCTCTCTGCATAAACTTTCCCTCTGTCATTCATT
GCTAGGTACATGTTTGATTTTTTTCCTTTGATTTGCACTTCTCCTGCCACCCTTGAATGAAATTGCAGAATAGCTTCTTGGTTGTAAGGCTCCC
TTGTCCCATATACGTTCCCTTCACTATCGATGACGATGTTATAACCAGTGGCACTATGCAACTGTTTCTCCCTACCGATGTGACCCGGTATGCC
GCGGTCCGTCTCCATGTTAGCAAAGCCGGAACCTTTCTGCTCCTCTTCTGTGGCCATCTTGTCGGGAGGCCCAAGAGCAATGGGATCCTGATCT
TGTGTACAGGTGGAATACCAAT
>Gk-Fgf1
WYSTCTQDQDPIALGPPDKMATEEEQKGSGFANMETDRGIPGHIGREKQLHSATGYNIVIDSEGNVYGTREPYNQEAILQFHSRVAGEVQIKGK
KSNMYLAMNDRGKVYAESDPESENTWFKEEYEDGWNYYSWLGNQKEWDLGIKKSGKMKRGLRTRRDQNA

>comp16539_(Gk-Fgf8)
CCCCGACTCCGTAGTTCTTCATTCTGTATTGCATCCCGCTCCGCTAGCAATAGTGAACGGACGTGTGACAACGCAGGAAATATCTATCAAAAAG
AAAAATTTTGTGCATAATGTAAGGTTTTTTTTTGCGGTCGTCTCTTCCGGATGAACACAGCTTAAAGGATATTTTTTGAAAGCGAAAAGAAAAT
TTGTTTGAGCAAGTTTTGTGTGGATTTTTGTAGATGTCTTTTATACATTTTTGGTGCGAGTTGGATTGTACAAGAGAAGTGACTCTTTAAAATT
TCGATGCGTGCCAGCGGATCCATTAGGAGACATATGTCTCCATGGTTACTTGTTGATGCCACATGTTAATATTCAGGTTTCTTCAAATTGCCTC
CCTAGCGTACCTGTGTAGTATATTCTTCAAAATGGCATCGTGTCAGAGTTTTTCCGTCGATTTGGAGAGGGGAGCTCCGTCGATCTTGAAAAAT
TCCCACGACGTACAAGCTCTCCGCAGGACGTACAAACTCTACAATCAGTGCAGTGGAAACCATATACAGATCATCGGTAAAACAGTCAACGCCA
AGGGTAGTCCCGACAGCCCTAATGCGAATATGAGCTTCACTAATGCCCGTTCTAGGCATCATTCCAATGCCATTAACATTATGGGACAAAAATC
TGGTCGTTATCTATGCTTTAATAAGAAAGGAAAACTAATAGTCAGGTTTAATGGAAGGAAAAAGTTATGTCTCTTTCGAGAAAATATCAGCGAA
GACCACTACACAGTGCTACAGTCATTATACGATTCAACATGGTATATTGGCTTCAATAGGAGGGGCAAACCTCTGAAAGGAAGTTTGTACTCCA
AGGCAAAAATGCAGCGGTGTTTTCACTTCGTCAAAAGAGATTATTCCTATGGCATGGATGAATACACACCACCTGGGCCTAAAATCCAAAATCC
TGGTAAATTAAAAGACTTGCTAACTGGAAATGGCCAGCGTCGATCTCGGTTGCAAAGAAAGGTGCCAAAAACGCCGACGAGGTGAAATATCCAA
GTAAAGATCGTGCTTCAAATGTTAGATTTCGAATTTTTTTGGTTGTTGCTTCGTTTTTCTCTAACTAAAATTTTTCGAAATAGACGAAGATGCC
AGGGCAGGCCTTGTGCGTTTTCTTCTCCGTTGACGCTTTGATGTTCGCCAGTGTGTGGCTGGCTGATCATGCTTACTATCTGTTTTGAGGACGT
TAGCTGGGGACCGGCTTTGTGTTATTGCAAACGAAGTGCCAATTTTGTGTTCTTAGGAACCCCTAAGAGTATTCAACCAATATCTCTTCGGACT
TTTCCTCGATGTTCAAATGCAATTAGGTATTCTCAGAAGTTCTTATCTTAGAATAGGACTTACATGTTATACGTAGTATTTTCCTGAAAGTAAA
TGTTATTTATCCTCAGAGTAGAAATTTATTTATTGCAATGTTAGTACCTTTTAGGAATTGTTACATTTAACTGTATTCAGAAAGTCGTTCCGAG
TGAAACTAGATTCATTATGTGATTTTTCTTTTCTGATATTCGATGACTAAAACTGGGAAACTTATTGAGAAATATGAC
>Gk-Fgf8
MLIFRFLQIASLAYLCSIFFKMASCQSFSVDLERGAPSILKNSHDVQALRRTYKLYNQCSGNHIQIIGKTVNAKGSPDSPNANMSFTNARSRHH
SNAININIMGQKSGRYLCFNKKGKLIVRFNGRKKLCLFRENISEDHYTVLQSLYDSTWYIGFNRRGKPLKGSLYSKAKMQRCFHFVKRDYSYGMDE
YTPPGPKIQNPGKLKDLLTGNGQRRSRLQRKVPKTPTR-

>comp16435_(Gk-Fgf17)
AACACAATTTTAATTCTGCAGTCGTCACACCAGTCTCTTTTGGTTGTCAATGCACAGAGCGTCGCTGTATCGCATTAGAATAGGAAGCCGACAT
TCTGGCAAATGATAAAAGAAATCTTGGGGAAAGATATCGTTTGTCATGTTGCATAAGACTGCTGCATACCGTTTTCTACTGATTTTTAGTATTT
CTATGGCTACTATATGTCTTGCTCTCTCTTCGGATGCCCTTTTAGAGAATGGGACCAAATTCTTCATTGATGACAATACGATATCTCGATACAG
TATTACAAGACGTTACAAATTTTATAACCATTGCTCCGCTAAACAAATGCTAATAGTCGGAAGTTCAGTAACGGCATCTGGTAGTCTGGATAAC
CCTAACACAATTTTAATTCTGCAGTCGTCACACTACAAAAATCACAAAGGGATTCATATAATAGGCTCGAACTCCGGGCGATATCTGTGCGGCA
GTAAAAAAAGCAAGCTAATAACAAAGTTTTCTGGTGCCAATCCTCGATGTGTTTTTGAAGAGTTACTCAGTCCTGATTTCTACACCGTGCTCAG
ATCTGTTTATAACAAGAGCTGGTTGGTAGGCTTTAACAGAAGAGGCAAGCCCATTCTGGGAGCAGACAGCACCAAAACACACAGAAGTCGGTGT
TATCATTTCACAAAGAGAGATCATTCATATCTTGATGCCTTGCACAAAACCAGGCATTACGCTCCAAAGATCTCCCATCCCCATAAGTTGATGC
ATCTTCTTCATGGGAAGCAGAAGTGGGTCACCTT
>Gk-Fgf17
MLHKTAAYRFLLIFSISMATICLALSSDALLENGTKFFIDDNTISRYSITRRYKFYNHCSAKQMLIVGSSVTASGSLDNPNTILILQSSHYKNH
KGIHIIGSNSGRYLCGSKKSKLITKFSGANPRCVFEELLSPDFYTVLRSVYNKSWLVGFNRRGKPILGADSTKTHRSRCYHFTKRDHSYLDALH
KTRHYAPKISHPHKLMHLLHGKQKWVT

**Zygiella dispar**

>comp23447_(Zd-Fgf1)
CTGAGTTCAAAAAAAAGAATAACCTAAATAAAAAAATGAAAAAAAAAAGGCAAAGAAACCCCACTTAATACCTGATTTCTAACATTTGCAGATG
CATTTTCTAACAGAACATCAATTTCAACATCCTTAAAGTAAACATCAATTAAACAGTCTCTGCAGCTTCTTAATTATAAGAAGGTTCTGCAGAC
CTGGGTACAAACTTCACAGCATTTTGGCCCCTTCTTGTCCTGTGGCCACGCTTCATTTTACCAGTTTTTTTTATCCCCAAATACCATTCCTTTT
GGTTACCCAGCCAAAAATAATAGTTCCAACCATCTTCGTATATTTCTTTGAACCAAGTATTTTCTGATTCAGCGTCATTCTCTGCATAAACTTT
CCCTTTGTCATTCATTGCTAGGTACAGGTTGGATTTTTTCCCTTTGATTTGAATTTCGCCTGCTGCTCTAGAATGGAACTGTAGAATTGCTTCG

TTGTTGTAGGGCTCCCTCGTGCCGTACACGTTGCCCTTTCCGTCGATGACGATGTTGTATCCAGTGCCGCTATGTAGCTGTTTCTCCTTGCCGA
TGTGACCCGGAATTCCCCGGTCCGGTTCCATGGCCCGGGCGTAGGCTGGACCTATTCGCTCCTCGCGGACGCACTTCCGTCCTCGGAGGCCAT
TTTGACGGGAGGCCCGAGAGCAGTGGGATCCTGTTCCTGCGTACAGGTGGAATACCGATCCATTGTGTCATCAACACTATTCCAGCACCAACTG
CACATCGCTG
>Zd-Fgf1
AMCSWCWNSVDDTMDRYSTCTQEQDPTALGPPVKMASEDGSASREERIGPAYARAMEPDRGIPGHIGKEKQLHSGTGYNIVIDGKGNVYGTREP
YNNEAILQFHSRAAGEIQIKGKKSNLYLAMNDKGKVYAENDAESENTWFKEIYEDGWNYYFWLGNQKEWYLGIKKTGKMKRGHRTRRGQNAVKF
VPRSAEPSYN-

>comp16798_(Zd-Fgf8)
AGCACAGCAGGCTCAGAGTCCTCACATCTCTTTTCTCCGAGTCTCGCTCAACTGACGGTCTGCTCGTTCACTCTCTCGCTCTTTTGTGACATCT
GAGTATTTTTTTTTCCTTCCCCTTTTTCGAAAAAACAAAGAGAACAAAAAGACAGCATTGTTGTGTCATTATGTAATTCATTCCCAACCTGTTC
GTGACGACTGCAAATGAATGTGACTTAAGTGGATACTATTTTCTTTTTCCCGGAAAAAAAAAAAATTAGCAAGTGCCAAGTGTGGATTTTGTAAA
TGTCTTTTACACTTTTGGTGCGAGCTGGATTGTAAAAGAGAAGCGACTCTTCAAAATTTCGATGCGTGCCAGCGGATCCATTAGGAGACATTTG
TCTCCATGGTTAACTGTTGATGCCACATGTTAATATTCAGGTTTCTTCATATTGCCTCCCTAGCGTACCTGTGTAGTATATTCGTCAAAATGGC
ATCGTGTAAGAGTTTTTCCATCGATTTGGAGAGGGGAGCTCCAGCGATCCTGAAAAATTCCCACGACGTCCGCAGGACGTACAAACTGTACAAT
CAATGCAGTGGAAATCACGTTCAGATCATCGGTAAGGTACTCAATGCCAAGGGTAGCCCCGACAGCCCAAACGCCAACATGAGCTTCATCTCTG
CCCGTTCAAGGCATCATCTCAATGCTATTAACATTATGGGACAGAAGTCTGGACGCTACCTATGCTTTAATAAAAAAGGAAAACTTATAGTCAG
GTTCAATGGCAGGAAGAAGCTGTGTCTCTTTCAAGAAGATATCAGCAAAGATCACTACACAGTGTTACAGTCAATATACGATCCAACGTGGTAC
GTCGGCTTCAACAGGCGGGGTAAACCACTGAGGGGAAACTTATACTCCAAATCGAGTGTACAGAATTGTTTTCACTTTGTCAAGAGGGATCATT
CCTACGGCATGGATGAGTACACGCCACCTGGGCCCAAGATCAAAAACCCTTGGAAACTAAAAGACTTGTTGACAGGACATGGTCACCGTCGATC
GCGGTCACAGAGAAAAGTGCCAAAAACGCCGTCGAGGTGAAACTGTCCAAGAATTTCCAATAAAGAGTCGTGCTTCAGATATTAGATTTCGAAT
TATTTGGTTGTTGCTTCTTTTTTCCTCGGACTGAAAATTTTCGAAAATTCCAAACGCCAGGGCTGGCCTTGTGCGTTTTCCTTTCCGTTGACGC
TTTGATGTTCGTCAATGTCTGGTTGACCATGCTTTAAGAAGCTATTTCGAGGACATTTGCTGGGGATCGGCTTTGTGTTATTGCAAATGAGGAA
GTGCCAATTCTGTGTTCTTAGGAACCCCTAAGAGTATTTACAAATATCTCTTCGGACTTTCCTCAATGTTCAAATGCAACTAGTATTCTCAAGG
TTCTAATCTTAGACTAGGATATTCATGTTATACGTAGTTTTTCCTGAAGTGAATGTTATTTATCCTAAGAGTAGAAATTTATTTATTACAACGT
TAGTCTCCTGTTAGGACTTGTTACAATTAACTGTGTTCAGAACATTGTTCAGAATAAAATTAGATTCACAACGAGTTCTTTTGTAAAACTGAGG
ACTTAATTGGAAAATTATCGTAGAAATATGATTCAGCTCTTCTCGACTTCATTATCATCTCAAAGTAGAATAAAATATGCAAAGACATCCTTTG
CTGATACCTGCTGAAAGTCTTCTGGCATTTAAATGTTCTTCAACCCAAACTAACTCCCATTGACGAGGGGAGGTGATTCGTCTTAAGAAAAGAT
GGACACTTCTCTCAGGGAAAGTCCAGTTACTCTAAGTCGGTGCAAAGTGTGGTTGCTGTCTGACTGTCGACAACCAGAGTCATGTGTATGTGAA
AACATCGTCTCCCATTATTTGTACAGAACATTTTTGTTTTGTTTATGTGTTGTTCTAATCATGTGCCATCGGTGAAGCTTCAATTCTAATGGTC
TTTTGTTTTCATACTTGTAAATGATTTATGTTTGTCTCAAATCATTCCAAACTGTGGCCAGTCCAGATTTTTATAAAATGTGACTCCAAGCCTG
AATACGCGAACAAACGGTGATGTTTCCAATTCTTGTTAGATGTTCAACGTCTAAATTTGTACATAGTTATGCTGTTTTGATGAACTTAGTCATT
TAATGATTTTAACAAAGTAATAATATTAATTTCGGATTTAAGAATACTTGTATAGTTTTAATCCATTTGAGTTTTTAACATTGATACGATACTT
ACAACTGTAAACTGTGTCGCTCATTTCCTCTATGCATGTTTGTTTAACGATTGAATCTACAGTATTTAAATAGTCTAGTTTAAAA
>Zd-Fgf8
MLIFRFLHIASLAYLCSIFVKMASCKSFSIDLERGAPAILKNSHDVRRTYKLYNQCSGNHVQIIGKVLNAKGSPDSPNANMSFISARSRHHLNA
INIMGQKSGRYLCFNKKGKLIVRFNGRKKLCLFQEDISKDHYTVLQSIYDPTWYVGFNRRGKPLRGNLYSKSSVQNCFHFVKRDHSYGMDEYTP
PGPKIKNPWKLKDLLTGHGHRRSRSQRKVPKTPSR-

>comp12612_(Zd-Fgf17)
TAAAACACTAGCCTAGACTATTTTTCTCGTATTTTGAATGGAAAAGAAATAAAAGCCAGACATACAAAGAATAGAAAGAAAAAAAAAAAAGCAGT
GTTACCCAAGGGTATTGGAAGCAGCGTTGTCTGGAAAAAGAAAGAATCAAAAACGGGAAAAAAAAACTCTCCGTCAAAATAGAGTTTTAATATTT
AATAGGCCATAAATCCTCAGTGGCCTCCTGTATACCTTACACTAGTGCTTTTGAGTACACTAATGATGGTATCTGGCAGAGATGTTGCAGATGT
CCTTATAAAAGAAAAGGATGGTGCCAAATTTTTGACTGACAATTCCATAGCAAGTTTCAGCCTCTACAAGAGACATTACAAATTTTTCAATCAT
TGCAGTGAAAAGCAAATTGAAATTATTGGAAGTTCAGTGACAGCATTAGGAGGCCCAGATAGCCCAAACTCAAATCTTATATTACAGTCAGCCC
ATTACAACAACCACTTCGGCGTCCACATAATCGGAGAAAAAAGCGGTCGCTTCTTGTGTTTCAATAAGAAAAGCAAGCTCATTACACGGTTCTC
GGGAGCGAGTCCACGCTGTATTTTTGAAGAACAGTTCAGTCCTGATTACTATACGGTATTGAAATCTGTTTTTAACCCGGACTGGTGCGTCGGA
TTCAACAGAAAAGGCAAAGCAATTCTTGGATCGGAAAAAGCCAATCACAGAAGCCGCTGTTTTTACTTTACAAAACGAGACCACTCATACCTTG
ATGGTCTGCACCAAACCAGGCATTATGGGCCAAAGATAGCCAATCCTCACAAGTTGATGCATCTTCTTCGCGAAAAGCGTGTGAGAAGAAGAAA
GCGGCACAACTTTTACGAGATGAGCAGGTGAACTGTACCAATGGAAGAAATTGTGTGGCCCTTCTGTATCAAGGTTGTGACCTGCTTACCATTA
CATACTTGACAAGGTCTCTTTATTTTTTTTGAATACGCGTTACTGAACGACAGTTACGGTGGTGCCACCAATTCCTTGCCTGCTTTTCTTCAAT
ACTATAAATGTCGTTCAATACTCAAATGCCAGTCACTGTGCAGTTTAAGTATAACCCCGGTTTCATAATTTCATCTGTTGCTTTTGTCATTGTT
AGAACTGTCACTAGAATTTAAGTGGATTGTAGCTATCCATGATTTACGCCTTTAAGCTCGTAGTGCATTTCGAAACTGCAGCTTTTACTGGATC
TGTGTCTAAGTATCAAGCTGCGACCATTTTTTTCTCCCTGTTTTCTAAAGTTAAGTTTACAGATACTTGTATGCGCTGACAGTATGGCGAGTCA
AGACCTGAAAATATGTTCCTATTGTTTGCATTTTCTTAAGGAATGTTTCATCTCTTGTTATTTTTACACATATATTTGTTTTGCATGACATAAT
TGCATCAATTAGAAGTCTGTTATTAACTTAGACGAAGAATCGTAATCTTCACATCGTATGCAATACATCTGTGAAAAGTAACAACTATTTTGAT
TGTTGATACATTTTTTTTCTTAATGAAAGTTCACCAGTGGAAAAAATTACTTTGAAATGTGAGTTTAGTGTATGATTGGTTTATCAGGGTAAAT
TGAATATTTTTTTTTTCACATTTTCTAAAGTTTGTGAACGAATAAATATTTATTGAGGATGATATAAATTTCTTATATGTACATAATTTATCTT
ACATTTTAACCATTGTTTCGTAATTTCCATTTTTAAAGAAGCAAATTTGTGAATATATATGGATTTTGCAAGATCTGGATGAAATGAATTTTAT
TGTATATGTATCAACAAAGAGCTTGAATTGATTTTAAACTATTTTTATTCGCATGTCATTCTCGAATCTTGATTTTTATTGAACGGCTTATAGT
GTAAAACTCTTGTATAATTCTCAAATACATTAAGGGCAATTTTTTTGTTATTCTGTTTACGTAATATTTAATGATATTTGTATAACAATGTGTG
AGCACACATTCCACTCATTAAAACTTTTTATTAACATACTCCAACATTCCAGGATAATATTCTATATGAGTATTTGATTTTTATATTTATTTAT
TGTTTTAAAATATATTTTTAAAATTATTTTTAATTTTAACAAAAAATAAATATTTATCTATATTCCTTTAAGGAAAATCTTCGAAATCTAGGTT
CATTAACGGTAAGAGCTTCATTAAAAAATACAATACACTTTTAGAATACTAAATTTCTGATGATTTCCCTATTCGTTTCATTCATTTTAATCAT
CATCACAATAGCATAATTTGTGTTCCACCTCTAGTCTATTACTACTTTTCGAAAATCAGACACAATTTTACTTCAATAAATTTTCATAAGACTA
CTATTTCAAGCAGTTTTTTGAATGAAACAAATTAAATCATCAGTCTTGGTGTTTTTTATTTTTTTTCCAACAGTTCCTAATTTGAAAATTTAAA
TTTGGTTCAGGTGAAAGGCATCAAAGTATATTACAAAATTGAAAAGTTATAAAATTTGTTTTCTTGGTTTATTTCCATTTTAAGGAACTTAGTA
ATTCATAAAGTCACGAAAAAATTAATTATGTTAATCATTTTGTAAGTATTAATTGCGTTCTTTAATCATAATACATCGTGCTTCTTTTGTGCAT
TTACCTTTTTTCTTGGTTACAACCATGATGGTCTGAAATGGATATATCACATTAAAAGCAAGAAAAGGTTTTAATTCCTTATTATTATAGTACC
GCAACATAAATGAAAATCGTTTCACATTTTATTAATAGTACGCTTTATTTATTAAAACAATTAGCATACTACTGTATTGAAATCCGAGTGCAAT
TTTCATCATAAGCAAATAAATGCATTTTAAAAAGTCTTACGACAAACTTAAAAACAAATTTATGAATCAATGCAATAAATTAAAACTTCTTCTA
AATACAATATTGACGTAATTTTAATGCAATGTAGTATTTTGTTGTACGAGAAACAAACGTTGCGTACCTAAATACTTATATACAGTCTGTTTTT
AAATATTTTACAAATTTGTGAAAAGTTATTTTTTCAATCTGTTTAGAAAAGTTGGATAATTATTTTTCCATGCTTTTATATAGAAGAAAATGCT
CTATTTTATGCATTTGTTATTGATTTTATCAAACGCTGTCTTTGTGAATCAAACTCTGTATGCATTTTCAGTAGACAAAGAGGAAAAACAAGAT
GTATTTTATGATATTTTTTAGCACAATAAGCGTTTTCATGGCGGTCTTATGGAGTGAAATTACTTTTAGCTTCAAAAAGTCAATGTATTTTTAT
AACTATGCGGTTGGTTTCATAAATGTGTGCCAATATTGATTTTATAAATGTTTCTAATGCCAGTCTACAATTTCAATTGTGTGCCAAAGTCCTG
TGAAGCTACCTCTTCCATTTAGAATTTAATACAATGCAATTTTCAGCTGCTAATAGCACAGTATATTCCTTCTGAGGTGCTAATTAAAGGACA

GATACTGATATATGACTTGTTTAATTATAAGAAGTGTTTCATGAAGAAACACAAAGACAGATTTCGTCTTAATAATCTATCTGATCTTGTCACA
AAGAAAGAATTTGTTTGAAAAATCTGACCTATCTTGTCACGAAAAAAGAATCTGTATTGCATAGGACCTAATAAAACTTTGTATCAG
>Zd-Fgf17
MMVSGRDVADVLIKEKDGAKFLTDNSIASFSLYKRHYKFFNHCSEKQIEIIGSSVTALGGPDSPNSNLILQSAHYNNHFGVHIIGEKSGRFLCF
NKKSKLITRFSGASPRCIFEEQFSPDYYTVLKSVFNPDWCVGFNRKGKAILGSEKANHRSRCFYFTKRDHSYLDGLHQTRHYGPKIANPHKLMH
LLREKRVRRRKRHNFYEMSR-

**Acanthoscurria geniculata**

>TRINITY_74449_c1_g1_i2_(Ag-Fgf1)
GGAAATGAAATTTTTTTTTCCCAATCATGTAATGACTACCCGCATGAAATCAACGTTCAGTTCTGTTCGAATATTCACAATTCTACAAGATAG
AAAAGAGTACTATACATTTCAAAACACCGTACTGGATGGTATGGCTGTATTAACAGTATTTTCCCCTCTACAGAAATGTCACATGAAACTTGAT
AAACACAGAATGATGAAAAAAGAAGTTTCCTATATGCATACAACTATGTGTAAGCTCTTTTGACACCAACGACTAATGAAACGAAGCATCCTGA
GATTTCTACACTTTCTGAGAATTTGTCATATCTACAATTAGAGAAGCAACTAATTTGCAATCTTCAACATCCGCTCACCGGGTCATTGAGAGTC
AAGTAACATCCATAAGCCACGTTCGCAACACTTATCTGGCGGGACAACGAAGTACGGGACAGTTGATTTTTTTTCTTTGGTCCTCTGAACACTG
ACGAAATGTGAATCGCTGCTGCTAGCAATGTCATAGCTACTGGGTTGGAGGGTATGCCGGTCTAGGAATGAACTTGACGGCTTCCTGTCCATAT
CCTGTCTTTTCACCTTCTTTGGGCTCGCCAGATTTTTTAAGGCCTAAATACCAGTTTTTGTCCCAGTGGACTTTGGAGAGGTAGTAATTCCACC
CGTCACCGTATTTCTCGACAAACACTATATTTGGTGATTTGGGATCAACCTCAGCATATACCCTGCCTCTTTCGTTCATTGCCACGTACATCTG
AGAAGATTTCCCTTTGATTCGAACTTCACCACGTGCCGGTGAATCTATTTCAAGAACAGCTTCTTCGCTATAAGGTTCCCTGGTTCCGTAAACT
TTACCTGAGGGATCGATGGCTAAATTGTAACCCGTGGCACTGTGGAGCTGCTTGATAGGCCCCAGGTGAGCAGGGAGGCCACTGTCATTCTTGC
CACCAAGGAAAGGTGGTTTGAAGTCGAAACCGTCGCCCTTGGTAGGATCAGGTTCCCCGGAAGTCCTTCCATAAACTCCGGCGCTGCCCGCTGA
ATAGTTGTCCTCGTTGCACGCCATGCTGAGCTC
>Ag-Fgf1
ELSMACNEDNYSAGSAGVYGRTSGEPDPTKGDGFDFKPPFLGGKNDSGLPAHLGPIKQLHSATGYNLAIDPSGKVYGTREPYSEEAVLEIDSPA
RGEVRIKGKSSQMYVAMNERGRVYAEVDPKSPNIVFVEKYGDGWNYYLSKVHWDKNWYLGLKKSGEPKEGEKTGYGQEAVKFIPRPAYPPTQ-

>DN82690_c5_g1_i1_(Ag-Fgf8)
CGCGTTATCACGGACGTTTCGTGAGTTCTGTTGTCTGTCTCGTATTTTTCTTTGAGGAATAGTTTCCGAGACTCGAGAGAGAG
AGAGTTCGAGTCATCATTAAAAAAACTCTTGGTTGCTCCAGAGAAAACTGGATATTTCATTCGCTTTCCTCGCTGAACATTCC
CTTGTGCGAATAATTTTTTCTACATTTTTACGTGCCTTCTTTCTCGGATGTTGGGGAACGTGACGTCCCCGTGATGACCGTCT
ATCGTAGTGACTGGGAATACCGTTGTGTGATCGAAGCCGCCGCCGGGTGTCAATCAATATCAGGTTCGTGCCGATGCTGCGCG
GGAAGCATCTTCGCCCGCCTAGATGATCGGAGACCCCTGTCTCCATGCTTCACCGTTGATGCCACATGTTAATATTCAGGTTA
CTTCAAGCCGCTTCCCTAGCGTACCTCTGTAGTTTACTCGTAGAAATGGTATCTTGTCAGAGTCTGTCTGCTGGTTTGGAGAA
AGGAGCTCCTGAAATTCTAAAGAATCACACTCCGTCGAAAGACATTAGGAGGACGTACAAGTTGTACAACCAGTGTAGCAGTA
AACATTTGCAAATAATAGGAAAGTCAGTCAATGCCAGGGGGCTCCCGAACAGTCCCAATGTGAATTTAAGTTTCGTCTCTGTG
CATACACGCCATAATCCCAACGCTATGAATATCGTTGGACAAAAATCAGGGAGGTTTTTGTGTTTCAACAAAAAAGGCAAACT
CATCGCCAGGTTTAACGGCCGCAAGAAACTTTGCCTTTTCAAAGAAGACTTCAGTGAGGATCACTACACGGTGCTCAATTCGC
TCTACAACACCAGCTGGTACGTTGGATTCAACCGAAAGGGTAAACCACTGAGAGGCAACCAGTATTCAAAGAGTAAACTACGG
GACTGTTTCCACTTCGTCAAGAGGGATCATTCGTACGTACAGCTGGAAGATTCTCGGAGACCCTGGCCTGGGCCTCCAATTAA
AGACCCAACCAAGCTGAAGGATTTGCTTACAGGGCGGAGAACACACATAGGTAGGACTGCAAAAAATAGGCCCCGTCCTCCAA
GGTGAAATGGTAAAGATCATAAAACGGGAATCTTTTATTTTGCCAGGCTGGACCTGGTGACAATCATGCGGTGCTACGTTCGA
ATATTGCGCCCCTTCTTCCAGCGTAGTAGTCTTTACTCCCTGCAGTCCTTTTGTGCGCTCTCTGTTCTTTGGAGCTCCTTCCG
TGACAAGTGCCAATGACAGCAGCTTCTGTATATCCGAACATCCGCACTTCTGTTTTCAACTGTTTTTGCGGTCTTGTACAGTG
TATTTAAATAGATATCATTGTCTCAAGTGTGTCATCTCAATGAACCAGTACTATGAAAACTCATTATCTAGCGTTGCAGATCA
TGTGTTTTCTGACCATTCCATTGCTGGTAGTTCTTGGTCACTGCACTCAGGTGAAGAAAGAGCTAACTGCACATGCAAAACTG
CATTCAAGAGCGTCAGCCGTCTCTCTGAAAGTGAGCTGTATTTTGACCGAATTTCTGTGAACTTACGGCGTGCAGAATATGCC
AATGAACAATGCATTTCATGTTTTTATTCTGTTTCCAGAATCTTCCATTAGTCAAGTTGTTGTTGTTGTTTATAGCAATCATA
ACATTCATGTGGTGAAGAAGTCGTCATCCACACGTTAACAGTTCTTGATTAATACCTTATTCACCTGCAATGATTTAAAATAC
GTATTCCATTATATTTTTTTTGTGGATTATGAGAATTTTCAATTTAAGTCTTAATTTTATAGAATATATCCAACAAGTAGTCG
CATGTACAATTTGCCCTGCAGTATAGACATTCAGAAAATTCTGGCAGGAAAGTTACGAATTAAACCTTATTTTTTTAAAACTA
AGTTTTCCGAAAGTTGAAAAAAAAGTAACTAAATCCATTAACATGATTTAACAAATCACAGTCGTTCGATGGATTACAATTGT
TGTACGTTTTCTGTCTCTGTCTTCAGTGCTTTAAAAAAATAGCCAAGGTGTCAGGCAGTAGTTCCGGAAGTAATGGGTAATCT
TTGCGTAGGCAGGTCATAATGAATAATAATGACATAACTGAGAAAATAACTAAAATAGGAAAATTTAAAAAAAATTGGCTGAA
ACTTAACTTTCTATAAGTAATTGATTGTATCACAAACTATGACATGCGCACAGATCGTAAGGTCTACTGTTCAGCGATAAAAA
CTAATCTTTTCACCGGATTATTCGATTACAATGCCCACTGCATTTTCACCCCCACCCCCCCACACACACACACAAGAA
>Ag-Fgf8
MLIFRLLQAASLAYLCSLLVEMVSCQSLSAGLEKGAPEILKNHTPSKDIRRTYKLYNQCSSKHLQIIGKSVNARGLPNSPNVN
LSFVSVHTRHNPNAMNIVGQKSGRFLCFNKKGKLIARFNGRKKLCLFKEDFSEDHYTVLNSLYNTSWYVGFNRKGKPLRGNQY
SKSKLRDCFHFVKRDHSYVQLEDSRRPWPGPPIKDPTKLKDLLTGRRTHIGRTAKNRPRPPR-

>DN76096_c8_g1_i1_(Ag-Fgf17)
CACCAACCAGAGACGGACGGCGGTAGTTGCATCCCCCCTTCTTTGGGAACATGGTGCCGGGCTCGTAGAGGGATCGGCGATCA
GCTCACTGGCACCTGGTTCCGACCTGCGGGACGGACGTCTGTTTCTGAGTCCTTGCGAAGCACCTCCTATGGGATACCTACCT
ACCTGGCCATGATACAAACATCCCTGACTACTGTGACCCCGACGATTGCCGGCGAACAGCCTGAGCAAATGCCACCAACCTAA
TGCCACATGTTTATACCCAAGTTTCTTCAAATTGTCGTGCTAGCGTTTTTGAGTGCACTGCTCATGGTGTCGTGTCAAGGGGT
AGCCGAAGTCCTCGAGAGGGGAGCCGAGAAACTGCTGAAGAAGGACAACGCCAGTGGCCTTCGTCACTACAAGCGTCAATACA
AGCTTTACAATCACTGCAGTGGAAGACATATACAAATCATTGGACAATCAGTCACTGCACAAGGAAAAGAAGACAGTCCTTTC
ACCGACGTGCTTTTAGAAACAGTACACTACCACAACAAATTGGGTGTCAACATCATCGGACAAAGCAGCGGACGTTACTTGTG
TTTCAACAAGAAAAGCAAGCTCATCACAAGGTTCTCCGGCAGAAGTCCAAGGTGTTTATTTGATGAGCAATTCAGCCCAGACG
CCATGTATTCTACACTGAGATCTGTCTACAATCCTGACTGGTACGTCGGTTTCAACCGCAAGGGGAAGCCCTTGCTAGGTGCA
GACCATGCTAGTAGTAGTCATCGTGAGCGTTGTTTCTACTTCACGAAAAGGGACCACACATACCTCGATTTTCAGGAGCACCG
GCATCCAGGACCTCCTATGCCAGACACCCATAAATTGCGGGACTATTTGAGAGAGCAGCGTCTCAAGAGGCGTCGACGGCAGG

```
TCCTGATGTCAACGGAAGCTGCCTTTCGACCTTCAAGATGAAATACGAACGAAACATTTCTAGGTGTTGAAATTAACACCTCT
ATCACGTTCCACCTTAGGTCACAAGCACGACCGATCTGTTGCCACTTTCACGTTAGCATTCACCGAAAACTGTTTGTTATTCG
CGCTCAGCTGTGCTCGCTCGCCGTGTCCAGCAAGATTCTCAGGTGTCCAGGACGTGGAAACAAAACACGAAGTGTCAAAAGGA
TTATAAGTCTTGTTTGACCTCTATAGTTCAAGGGAACTCCATTCGCACCTTGCAAACTAGTTTATTGGGCCCTCGTAGACAGG
CCTGTTTGTGCGCTAAAGAATCTCAAAAGTCTCAGTGTCTTTTATTTCAGCTTTTGGTAAATGACACTTTGTTGATTTCCCGG
AAAGGAAGAGAATATTTGTACATATTACCCCATTCCGCAAATTTAAAATAGTAATAAATGTGCTGCCAGCGTTCTGTTGGAGT
GACTTCCATTACATTTGTGCTGACGGAACTTTTGATATTAGAATTAACAAACTGTGAAGGACATTATCCGTTGTGAAAAAATC
AAAGTGTGTAGCATTTGATAGTGGGTCTACTTATATAGCTGTTTGATGTTTTTTTAAGAAAAGACTTTTGTTAAAAGGAAATA
TTTATAGAATGTTTGTGTTTGAAAGAACTCTGCGCTGTAAAATGTGTTGAATTAGCAACGTCGCACCTTCTGTCACAGAGATA
AAAATAGCAGGGTTTCCTATTTAGAATTCGTGAAGTGAATTTTTTTTTTTTTTCAGCTGAAGGTGTTCGTCACACCTTTGGAA
GAATTAGTTTCCAAACACCTGTACAGAAAATCCTACAATGCCAAAAGGCATCATTTGATATCCGAACAACTACAGAAAATGGC
ATTCTTCGTTAATATGAAATGTTCCATATTGTTTAGAGACGAAAGTGCTTTTTTTGTTAATCAGAAACACTGCTTTAGTCATT
AATATTTCAATTTTGGAACATCTTTGACATACGATGGATTATTAATTACCAAGTGAAACCTTTCAGTAAATGTGTGCTTAACC
GCTTTTAGTGAAATAAAGCAACACCAAACACACAGCAGA
>Ag-Fgf17
MFIPKFLQIVVLAFLSALLMVSCQGVAEVLERGAEKLLKKDNASGLRHYKRQYKLYNHCSGRHIQIIGQSVTAQGKEDSPFTD
VLLETVHYHNKLGVNIIGQSSGRYLCFNKKSKLITRFSGRSPRCLFDEQFSPDAMYSTLRSVYNPDWYVGFNRKGKPLLGADH
ASSSHRERCFYFTKRDHTYLDFQEHRHPGPPMPDTHKLRDYLREQRLKRRRRQVLMSTEAAFRPSR-
```

```
>TRINITY_DN82175_(Ag-dof)
GTGGGTAACCCTTTCCGAACTTCGCTCCGGGTGGCTAGAATCGATTCCAAACACGGGAGTGTAAATAACATTCGCTGGAAGGGAATTCGTTTCG
GAGATAGGCACTTGTGAACTGGGAAACGTTTGCTCAGAGCTGCGGTGTCCGTTTTTCCCAGAAAGAGTTGCTAGTGTGTTCCGGCTCATGCGGC
GATTTGCTCACGCCGCATTAGCAAGCATTCTCACCAGGAGGCAATCATAATCCCTCCGCAGGAAAACACCGAAGATGCCCAAAAAAGGTAATCG
AAACAACATGGCTCCGCCCGATTTGACGTTACTTTTCAGCGGCGAAGTCTCAGAGTGGGACAGCTATCTGACTCGACAGTTTGAGAATTCGTGC
GATCCTGCAGTAATCGAACACGAGCAGATCGAAGAGGTCTCTTTTCCCATCAGCACATCCCAAAGCCGGCACTTCACGGACGCCAGAGCGGTGT
TAGTGATCCTGTCGCCGTCGTTCTTGGACTTTCTCGATCGACATGCGGACGAAGGATTTCAGATCGGCAAGCTTTTGAAGCCGAGCCGGACGGT
GGCTATGTTGTGCGGTGTCCAGCCGTCAGACGTCTCGTCCTTCCACAAGGCTGCCCTCGTTACGTATGACGACTGGCCACGTATCGTGGCCCGC
GACAAAGATGTCAATATGGTGCGAAGTGTCGTGGACACCGTTCAGGAGCTTCTCACGAGATCCAAAGAGAACGAACGAATACACAAACCAAAGT
TTAAGCTCATTCCTCGAAAAGTTAATGAGGGAAATGGGAAGGTTTTTGTTATTCTAAACAAGCCTCTCGAAGACTCTAAAAAAGTGAAAATCAC
ATTGGAAGGACATGGTGTGAAGGAAGTTCCTGTCAAGAGAAGGAATCCCTACACTCTGCAGTTTGTAGTGCCAGATATTTGCATGCAAGTACTG
AGTGTCCTGAATGTTCATGTACGGTGCAATGATGATCTACTTGGAGTAAGCCAGCTTAAGTGTGAATCCCGAATGGGAGAACTGTATAATCTCC
TTTCATCCATCAATTCCCCATATGAGTTTATATGTCATACTTTAGGGATGGCAACTTTATATGAAGTTGATGAGCACATGGTGACAGCCTTTCG
GAAGAATCTGCCACCATCAGGATTCAACCTTCTGCATGTGGAGGAAAAGACTGAAAAGAAGAAAAGTAAAGAAGAATTTCCCACACTACTCCAC
TTTGCTGCTCGATATGGGCTGTGCAGCCTTTGTAAAGAGCTGCTTCGGTGCCCTGGTTCACAAACAGCTTGCTCCTTGAAGAATTCCAGAGGTC
TGACCCCTGCCCAGATGGCTAATCATGCTAGTCATAATGAAATTGCAACTATACTTGAGAATTTTCAGAAGTCTGTCACTACTTCTGAGAAAAC
AAAAAGCACCCCGGTTGCACAACCTTCAACTGTAGTATATGAACAAATGCAGGTAGAGTCACCTGCTGAAAATTATGACATTCTTCCTTCAAAC
CCTGCAAGACTTGTGCTTCCACCTACTCCTACATCTGAAGAAGAAGAAGCAGTTCTTGGAAAAGAGTGTGAATATTTGAAAATGATGGGAACTA
GTCCATTTGATTCAGAAACTTTTGAACCTCACAAAAAACTCATACTGGGTTTTGACCACAAGAGAGACGAGAGCGAAGAAGTTCTGCTTTCCTC
TCTAAGTGGCGACAGACCACCTTCACCACAAAGAATTTCTCAGGGAGATAAGGCAAAATTAGCATTGCTTGAAAGTCAAGCAAAGCTAATTGAA
ATTATGGAAGCATACAAAAAAGGAGCATCCTTGATTGAAGTAGAAAAACTGCATAATGAATGGAAAGCTATGTATCAAGTATCTGATGTAGAAG
CAAAGAGCACTTTAGAAGGCCTAAGGAACATGTATGCTAAAGCACAGAAAGCAAAAATCAACAAGAAACAGTCATCAAGCTTTTCTGAACTGCG
ACAGTTTTTGGGCTCTAAACTTCGAAGAAAAGATAGTGTTGGCTCAAAGAGCAATTCCAGTCGAAAGAGTTCCAAAGAGTTGAGCTGTATAGAA
AAACTGTGTCTTGAAGTTGAAGGATTGAGGCCTGTCAGTACACTAAGCATCATGTCAAACACAAGCAGCTCAAGTGGCTCATCCTTTGACCGAC
TGAGTTCTGTGAGTGGTGGATCTGACAGTGGGAATCATTCTGATTATGGTGAAGAAGCAAACCGCTTAAGGGAGGCAGGAAAGTTTGAATCAAG
ACTTACACATCCAATGGGATTAAATGGAAACTTGCTGACATCACAACCAAGCAACATTCCATTGTCATTGAATAGTAAGCCTCCTGCACCACCA
CCTCGTCCAATAAAACCACCTTCATTCATTAATAAATATGAAAATAGGATATATTCTAATAGCTTGGATCAGACAGCTTATTCTGTGCCACATA
AGCTTCCTACTTCTTCGAGTACCCCTCCGGGAAAGTCTCTGGACTATGTCATGCCAGACTGCACTCTGGGGAGTGGAAACACTTTCAAGTCTAA
GAAGAGTGCCATGTCAAAATCAGCCGACGATCTGCTGTCAGTTTCAAGTGGGGACTGTAAGATTTTGAACGCCTGTGATTTGTATGACACACCA
CCCCCTGTGGGAAACCTCTTCTTGAAACAGCAAACTACAAGTGTGGGTAGTGACATGCCTAAGGTACCTACCAGTCCCACCCATAGCTGTGGAC
TAAGTTTCTTAAATGCAGCAAAGAATGAATGCTATGGTATGGACTATGACTTAGTTCCACCTCCTCTTCCAGTGTCAGTGCCTCCAAGAAGACC
AAATTTTCCAAAATATGAAAGGAGCAGCAGCTTATCCAATTATGATATTCCCAAATTTGCTCCTCCTACTCCTACGTCGAAAGAAATCCTTGAA
GCAAATTATGATTTCCCACGATTAGATCAAAACAGTTTGTGCAGTTGTTTATTGAAACCAAGAAAGGAGTCCCTTCCTCCATGTTGTTGTTCGG
CAAGTTCTTATGATGTGCCATTGTCACCACGAAAATATAGTCTTCCCAACACACCTCTAAGTGATGCCCACTCAGGAAAGAGCTTATCAACATA
CTATAATATGCATTTTGGTAGGAACTCATGTAGCAGTGATGAACAGTGTGACAGTATTCCTGATGAACCTCCCCCGCCTCCACCACAAGAGCTA
AGTGATGAGGAAGAGCATTACAAATTAGTTGGTATTCCTGTACCTGTTTCTGGTGCTAATTTAACAAACATTGCATCCAAATGTTGAGTGTTCC
AACACTGATAACTCACAAAGACCAAAGAACTGTATTTGTGTACATCTATGTATTGTGATTCAGCAGAAATTTCTTGTGTGCCATTTAATGTATT
ATGGTTTTTGAAGATTATTTTTCATTCAATTTTGTAAAAAGTAACTATGATGTAAAGTCACTATAACTATTAATCACCTTGGTAGCATTCTTC
CTTTTGTTTATATGGACATAAAAGTTCGATTTGTGTTGGATTATGTCAGATGTTATTAAAAATGGACTTTTTTTGTATTAAGGTAAATGAAATA
TATTTCTGTCAGATTCCATGTAAATAGTCTTTTAAGTTTTTCAAATCTTGTCAGGTTACAGATCTCATTCAGCAAATACCAAATATCAGTTTTT
GTTGAGATAAACAGTAGTATATGACCCACAGAAAGGAATATAAAAGGAGCAAACTGACATATTTAAGTAGTTGTGATATAAATAAAATAACTAT
TATCTAGTGTTTGCTATTTTTTAGAGTGGTTGTTAATTAGATGTAAAGTTGCCCAGTTTAGGCATACCTATTAGTGATGCATTCAGGAAGTGCA
TTCACATGTGAGCGATATCAGGTAAACTATATTCTTTATATACAGGGTGCTTGGTAAAGTCCCAAGTCGCCCGAATGACTTGGGAGCGATGATG
GCTAGGGCTGTGGAGGTTGGGTAGGGGTGGGGTGGGG
>Ag-dof
MPKKGNRNNMAPPDLTLLFSGEVSEWDSYLTRQFENSCDPAVIEHEQIEEVSFPISTSQSRHFTDARAVLVILSPSFLDFLDRHADEGFQIGKL
LKPSRTVAMLCGVQPSDVSSFHKAALVTYDDWPRIVARDKDVNMVRSVVDTVQELLTRSKENERIHKPKFKLIPRKVNEGNGKVFVILNKPLED
SKKVKITLEGHGVKEVPVKRRNPYTLQFVVPDICMQVLSVLNVHVRCNDDLLGVSQLKCESRMGELYNLLSSINSPYEFICHTLGMATLYEVDE
HMVTAFRKNLPPSGFNLLHVEEKTEKKKSKEEFPTLLHFAARYGLCSLCKELLRCPGSQTACSLKNSRGLTPAQMANHASHNEIATILENFQKS
VTTSEKTKSTPVAQPSTVVYEQMQVESPAENYDILPSNPARLVLPPTPTSEEEEAVLGKECEYLKMMGTSPFDSETFEPHKKLILGFDHKRDES
EEVLLSSLSGDRPPSPQRISQGDKAKLALLESQAKLIEIMEAYKKGASLIEVEKLHNEWKAMYQVSDVEAKSTLEGLRNMYAKAQKAKINKKQS
SSFSELRQFLGSKLRRKDSVGSKSNSSRKSSKELSCIEKLCLEVEGLRPVSTLSIMSNTSSSSGSSFDRLSSVSGGSDSGNHSDYGEEANRLRE
AGKFESRLTHPMGLNGNLLTSQPSNIPLSLNSKPPAPPPRPIKPPSFINKYENRIYSNSLDQTAYSVPHKLPTSSSTPPGKSLDYVMPDCTLGS
GNTFKSKKSAMSKSADDLLSVSSGDCKILNACDLYDTPPPVGNLFLKQQTTSVGSDMPKVPTSPTHSCGLSFLNAAKNECYGMDYDLVPPPLPV
SVPPRRPNFPKYERSSSLSNYDIPKFAPPTPTSKEILEANYDFPRLDQNSLCSCLLKPRKESLPPCCCSASSYDVPLSPRKYSLPNTPLSDAHS
GKSLSTYYNMHFGRNSCSSDEQCDSIPDEPPPPPPQELSDEEEHYKLVGIPVPVSGANLTNIASKC
```

**Dysdera silvatica**

```
>TRINITY_GG_25065_c0_g2_i1_( Ds-Fgf1)
ATTAAATATAATTCTGGATAAATTCTGGAAATAGAATTCAAATGTAAACATTTGAAGTTCTATTGGAAATCGACATATTAGACCAGCTATAGCT
ATTTTCAATGCACCCAGATGAAAAATTAAAAATAATGCAAAAAAATGTTTTTATCATTACTAAAAAGTGCTGAATAGATATACGCTTTATAAAC
TGCGCCAATCTTTGATAAAATATAACCTATAAAATAAAACGTTCACAAATAATTTAAAAAATGCAACAATTGAAAAACATTTGCATGTTTTTCT
GTAAATTGTAAAAATTCTACTAGCGAACTAAGATCATGATCCAAAACGACAGCTACATCTTTTAAATCTTATCTCCCAACATTTAAAAGCTAAT
ATAGCTCCTAAACACTTTGAACATATAAGCTTCATAAACATGATGAACGAAATGAAGAAAGCTCCTAGAAATTTTAGGGCAACTTTCAAATCAT
GCAAATCACAGTAATTGATTGGACATCTCTGTATCATGCCTCACAACCTTTTCCTATATTAACTTACGTCACACAGTGACATATTCAGATGTTA
GCACCAAGCTAAGATCTAAAATCTTTGGTTGATTGTGGTATTCCTCACATGGACAAATTTGTTTCACAGTAACAACAGTTAACCCAGTTACATT
GTCACGGATACCTTCTTAAAATTAAAGATGACGATAAAACACTGGAAACACCGAGATTTGGTTTAATCAGCAGCAGTATCAACATCAGTATGTT
CTTGGATAAGCCGATCTTGGCAGAAAACGGACAGCCTTTTGACCTGGTCTTGTTTTATTTCCCTGTTTGGTTTTGCCAGATTTCTTGATTCCCA
ATAGGTATTCTTTTTCAGGATAACCTTCTTTCCATATATAGTAATTGAATGGCTCATCAACACGTTCAAGCCAAAGGGTACAATCCATATTCGG
GTCTCTCTCGCCATAAACTTTTCCCTTGTTGTCCATCGCGATGTAAAAGCCACTGTTACGTCCTTGAAGTTGAACTTCTCCAGGCATGTTTGAG
TGGAACTCTAGCACAGCATCCATATGAAAACGG
>Ds-Fgf1
RFHMDAVLEFHSNMPGEVQLQGRNSGFYIAMDNKGKVYGERDPNMDCTLWLERVDEPFNYYIWKEGYPEKEYLLGIKKSGKTKQGNKTRPGQKA
VRFLPRSAYPRTY-


>TRINITY_GG_36900_c0_g1_i1_(Ds-Fgf8)
ATAGTTTGCTCATAGAAATGGTATCGTCCCAAAGCCTCTCCGCTGGTCTTGAAAGAGGTGCTCCTGACATCTTGCGTAACAATACTGTGTCTCA
GTACATCCACAGAACGTATCAGTTGTACAATCAGTGTAGCGGGCGGCATGTACAGATCATCGGCAAAGCCGTCAACGCCAAGGGAACTGCGCAT
AGTCCAAATGTGAATTTAAGCTTCGTTTCTGTACATGTACGCCATCATCCCAGCGCCATGAGCATTGTTGGTCTGAAGTCAGGACGTTTTCTGT
GTTTCAACAAAAGGGGAAAACTTATAGCCAGGTTCAATGGGAAGAAAAG
>Ds-Fgf8
SLLIEMVSSQSLSAGLERGAPDILRNNTVSQYIHRTYQLYNQCSGRHVQIIGKAVNAKGTAHSPNVNLSFVSVHVRHHPSAMSIVGLKSGRFLC
FNKRGKLIARFNGKK


>TRINITY_GG_24745_c0_g1_i1 len=262 path=[1:0-261] [-1, 1, -2]_(Ds-Fgf17)
GGAGAATGATATATTTGCCAAAGGAACTAATAATTACAAGAGACGATATAAGCTGTATAATCACTGTAGCGAACGGCAAGTCCAGATTTTATCC
GAAGGGGTCAATGCAAAAGGCAGCAAAGACAGTGTATTCACTAATATTATAGTAGACACAGTCCATTCCCACTCAAACAATCACTATGGTGTCA
ACATAATTGGAGAAAAATCAGGCGTATATCTCTGCTTTAACAAAAAAGGCAAGCTGATCACAAGGATATCTGGC
>Ds-Fgf17
ENDIFAKGTNNYKRRYKLYNHCSERQVQILSEGVNAKGSKDSVFTNIIVDTVHSHSNNHYGVNIIGEKSGVYLCFNKKGKLITRISG
```
